# Physics-Grounded Evaluation to Guide Accurate Biomolecular Prediction

**DOI:** 10.1101/2025.06.30.662466

**Authors:** Ningyi Lyu, Siyuan Du, Qianzhen Shao, Zhongyue Yang, Jianpeng Ma, Daniel Herschlag

## Abstract

Deep learning has revolutionized protein structure prediction, with function prediction on the horizon^1,2^. Biomolecular properties, including structure and all aspects of function, emerge from atomic-level interactions and the probabilities of their formation^3,4^. Learning the physical rules that govern these probabilities allows models to deliver accurate structure predictions and may enable extrapolation beyond the training data—a capability needed to predict the many biologically important functional properties where comprehensive data are not readily attainable^5^. Current structure-based models follow a training logic primarily focused on matching atomic coordinates, rather than atomic interactions and their probabilities. It remains unknown whether the models have learned the physical rules that underlie atomic interactions, the extent of their knowledge, and the prediction errors that arise from limits to this knowledge. We found that state-of-the-art structure prediction models, AlphaFold2, AlphaFold3, and ESMFold, capture basic energetic principles but show pervasive biases in the conformational preferences of molecular interactions. These biases manifest as widespread prediction errors, including the misassignment of a large fraction of side-chain non-covalent interactions—∼30% for the AlphaFold models and ∼60% for ESMFold—and an inability to reproduce experimentally derived conformational ensembles. More than half the errors made by AlphaFold2 and AlphaFold3 are in common, suggesting limitations not overcome by using different model architectures. Overall, our multifaceted, physics-grounded evaluation identified previously unknown, system-wide deficiencies in current structure prediction models. This framework is applicable to and needed for all biomolecular structure and function prediction models that deliver atomic-level structural information. The insights derived from these evaluations will allow researchers to judiciously apply current models and will guide the development of next-generation models to achieve accurate prediction of biomolecular function.

## Introduction

Accurate prediction of biomolecular behaviors––from protein and RNA folding to ligand binding, enzyme catalysis, and allosteric regulation––will transform industry and medicine. Deep learning holds great potential for achieving this goal, as exemplified by the astounding success of AlphaFold and related algorithms in protein structure prediction^6–9^. This success has led to great excitement and widespread ongoing efforts extending beyond structure to predict biomolecular function*^e.g.^* ^10–12^.

A growing number of studies have used or repurposed structure prediction models to predict aspects of protein function, including folding stability^13,14^, ligand binding affinity^15,16^, and alternative conformations^17–19^. Nevertheless, a similar number of tests have suggested a limited ability to extrapolate structure-based models to these applications*^e.g.^* ^20–22^ (Supplementary Table 1). Given the large user base of these models and rapidly expanding efforts to develop new biomolecular models, there is an urgent need for effective evaluations that can determine the capabilities and limitations of the models and thus can guide their use and their improvement. Ideally, these evaluations would use metrics that are tightly connected to the properties that reflect the model’s predictive power, rather than surrogates or indirect correlates of these properties.

Biomolecular properties emerge from atomic-level interactions. Structures entail atoms arranged to form interactions—covalent bonds, backbone and side-chain torsions, hydrogen bonds, van der Waals interactions, *etc.*—with the sum of their energetic contributions representing the minimum on an energy landscape (Fig. 1a). Structure prediction, in its most accurate form, captures this energy minimum. The same physical principles also govern functional processes: for example, ligand binding introduces new interactions, and the sum of the energetic differences between the interactions in the apo and bound states determines the ligand’s binding affinity (Fig. 1a). Learning the physical rules universal to all biomolecular properties will allow models to provide accurate predictions and enable them to extrapolate beyond their training data^23^—the latter being required to predict the many functional properties where comprehensive training data are not available or readily attainable^5,24^. However, it remains unknown whether deep learning modes have learned these physical rules, the extent of their knowledge, and the prediction errors that arise from limits to this knowledge.

**Fig. 1.**
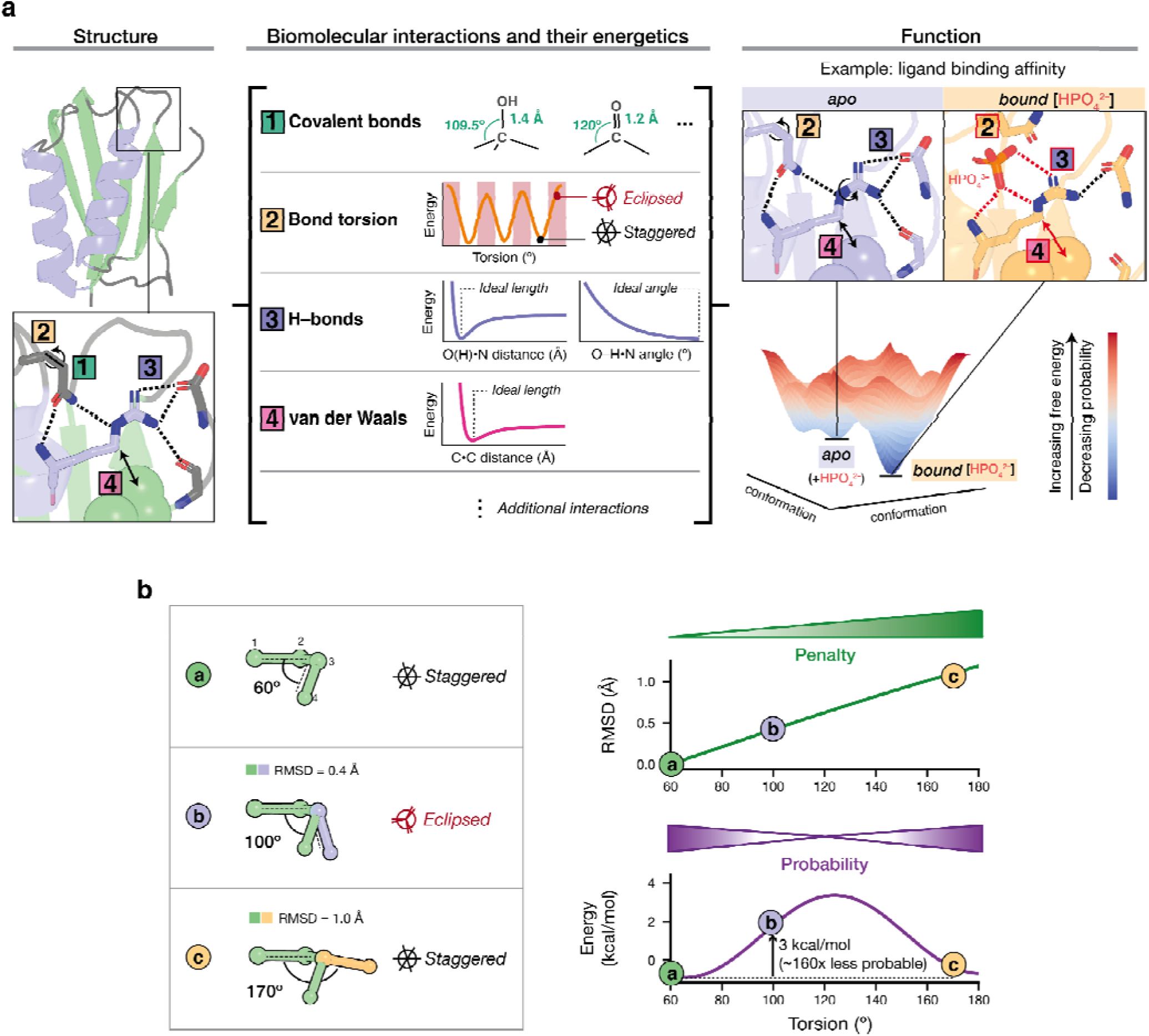
Molecular interactions and their energetics underlie protein structure and function, but model evaluation relies on distance-based metrics rather than interactions and their probabilities. **(a)** Structures are defined by molecular interactions (left; PDB: 2FHM) with each interaction following distinct energetic rules that determine its conformational preferences (middle). When combined, these energetic rules define the multi-dimensional conformational landscape that encodes protein function; a simplified two-dimensional energy landscape for ligand binding is shown as an example (right). Different interactions formed in the apo and the bound state of the protein and determine the free energy difference between the two states (*i.e.*, binding affinity). The example shown is acylphosphatase (PDB: 2FHM and 3BR8); red dashed lines and numbered boxes indicate different interactions formed in the bound state *versus* the apo state. The colormap indicates the free energy of the conformational states, from blue (low energy, high probability) to red (high energy, low probability). **(b)** The torsional rotation around a bond and the corresponding changes in RMSD and energy. The left subpanel shows a bond rotating from its original state (a) to states (b) and (c). This rotation is quantified by the torsion angle (defined by atoms 1–2–3–4) that changes from 60° to 100° and 170°. The right subpanel compares the changes in RMSD (top) and energy (bottom) from this rotation. The energy values are calculated from a smoothed knowledge-based energy function derived from crystallographic data (T = 298 *K*), as described in *Methods*. The green color map from light to dark green indicates the increased penalty in model assessment; the purple color map from light to dark purple indicates the direction of increased probability (lower energy).

Physics-based evaluations have been critical for developing predictive models, even for systems much simpler than the structure and function of biomolecules. Early models of small molecule generation were limited in value for drug discovery because a large fraction of the generated molecules were found to give invalid valence and highly strained geometries^25,26^. To overcome this problem, a series of benchmarks placed physics at the forefront of model evaluation^25,27,28^, thereby guiding the development of subsequent models that nearly fully eliminated valence errors^29,30^ and greatly reduced high-energy molecular arrangements^31,32^.

Biomolecular models are challenged to learn more complex interactions than small molecules as well as higher-order interaction networks, yet the current evaluation paradigm does not incorporate physics. Indeed, assessments of structure-based models near-universally use Cartesian distance based metrics that compare the distances between atomic coordinates in three-dimensional space, such as root-mean-square deviations (RMSD). These metrics collapse the multi-dimensional conformational differences to a single value which does not evaluate the accuracy of individual interactions and energy terms*^e.g.^* ^6–9,33^ (Supplementary Table 2). Consequently, improvements in these aggregate metrics do not necessarily lead to improvements in the functional predictions where accurate physical interactions are needed. For example, AlphaFold2 predictions were substantially improved in their RMSDs than traditional homology models when compared to the experimentally determined structures; yet, in ligand docking, these AlphaFold2 predictions did not give more accurate ligand-bound poses than the homology models^34^, suggesting critical features not captured by RMSDs.

To determine the capabilities and limitations of deep learning biomolecular models and guide their application and development, we provide a generalizable, physics-grounded framework to systematically evaluate whether models have learned the biomolecular energetics that define structure and function. We applied this approach to evaluate three state-of-the-art structure prediction models—AlphaFold2, AlphaFold3, and ESMFold—interrogating >3.4 million molecular interactions across 3939 structures. We found that the models have extracted energetic properties for multiple types of covalent and non-covalent interactions. Nevertheless, all models showed systematic discrepancies in the conformational preferences of these interactions that were previously unrecognized. These discrepancies are sufficiently consequential to result in prevalent prediction errors in molecular interactions and interaction networks—including the misassignment of ∼30% to ∼60% of side-chain•side-chain interactions—and an inability to predict conformational ensembles. This evaluation framework is generalizable to any model that delivers atomic-scale structural information, providing a new paradigm to define the capabilities and limitations of current models and guide the development of next-generation models that capture the biomolecular energetics needed for accurate and quantitative function prediction.

## Results and Discussion

### An evaluation framework based on molecular interactions and their energetics

To assess the ability of structure prediction models to reproduce molecular interactions and their energetics, we used evaluation metrics that directly map to energetic properties—covalent bond length, angle and torsion, hydrogen bond length and angle, and van der Waals distance (Fig. 1a). Fig. 1b illustrates the difference between interaction-based metrics and Cartesian distance-based metrics, using bond torsion as an example. A bond rotation of 40° from a staggered to a partially eclipsed state yields a ∼3 kcal/mol increase in energy (corresponding to a ∼160-fold less probable conformer at 298 *K*), while an additional rotation of 70° leads to a different staggered state with an energy similar to the starting state (Fig. 1b). By contrast, RMSD increases monotonically with the rotation, resulting in an increased penalty even for the lower-energy, higher-probability state (Fig. 1b).

We compared 3.4 million interactions predicted from AlphaFold2, AlphaFold3, and ESMFold to those obtained from X-ray crystallographic structures (*n =* 3949; Supplementary Table 2). These structures were part of the *Top*2018 dataset, with resolutions higher than 2 Å, and they were filtered by model quality at the single residue level^35^. Identical sequences were used to retrieve AlphaFold2 predictions from its database^36^ and to predict structures using AlphaFold3^9,37^ and ESMFold^7^ (*Methods*). The structures were confidently predicted, with an average predicted-local-distance-difference-test (pLDDT) score of 94 (SD = 4) in AlphaFold3 predictions, 96 (SD = 4) for AlphaFold2 predictions and 89 (SD = 12) for ESMFold predictions.

Below, we illustrate the evaluation framework with AlphaFold3 as the primary example and demonstrate the generalizability of our framework by comparing the performance of AlphaFold3 to that of AlphaFold2 and ESMFold. By comparing matched pairs of high-quality experimental structures and model predictions for the same sequences, our evaluations minimized discrepancies arising from errors in the reference structures and thereby more cleanly isolated errors from model prediction.

### AlphaFold extracted basic energetic properties of molecular interactions

AlphaFold3 reproduced the overall patterns of conformational preferences for covalent and non-covalent interactions. We found that the backbone torsion angles of predicted structures follow the classic Ramachandran distributions (Fig. 2a and Supplementary Fig. 1), consistent with the ability of AlphaFold3 to reproduce protein folds^6^. For side-chains, the predicted bonds showed preferences for staggered over eclipsed conformers, as expected, and the overall distributions agreed with those derived from the PDB (Fig. 2b and Supplementary Figs. 2 to 4). AlphaFold3 employs a diffusion-based architecture that directly generates atomic coordinates so that covalent geometries are not predetermined. We found that AlphaFold3 captured basic covalent bond properties, such as the ∼0.2 Å shorter C=O compared to C–O bonds (Fig. 2c, Supplementary Fig. 5 and 6, and Supplementary Table 4 and 5).

**Fig. 2.**
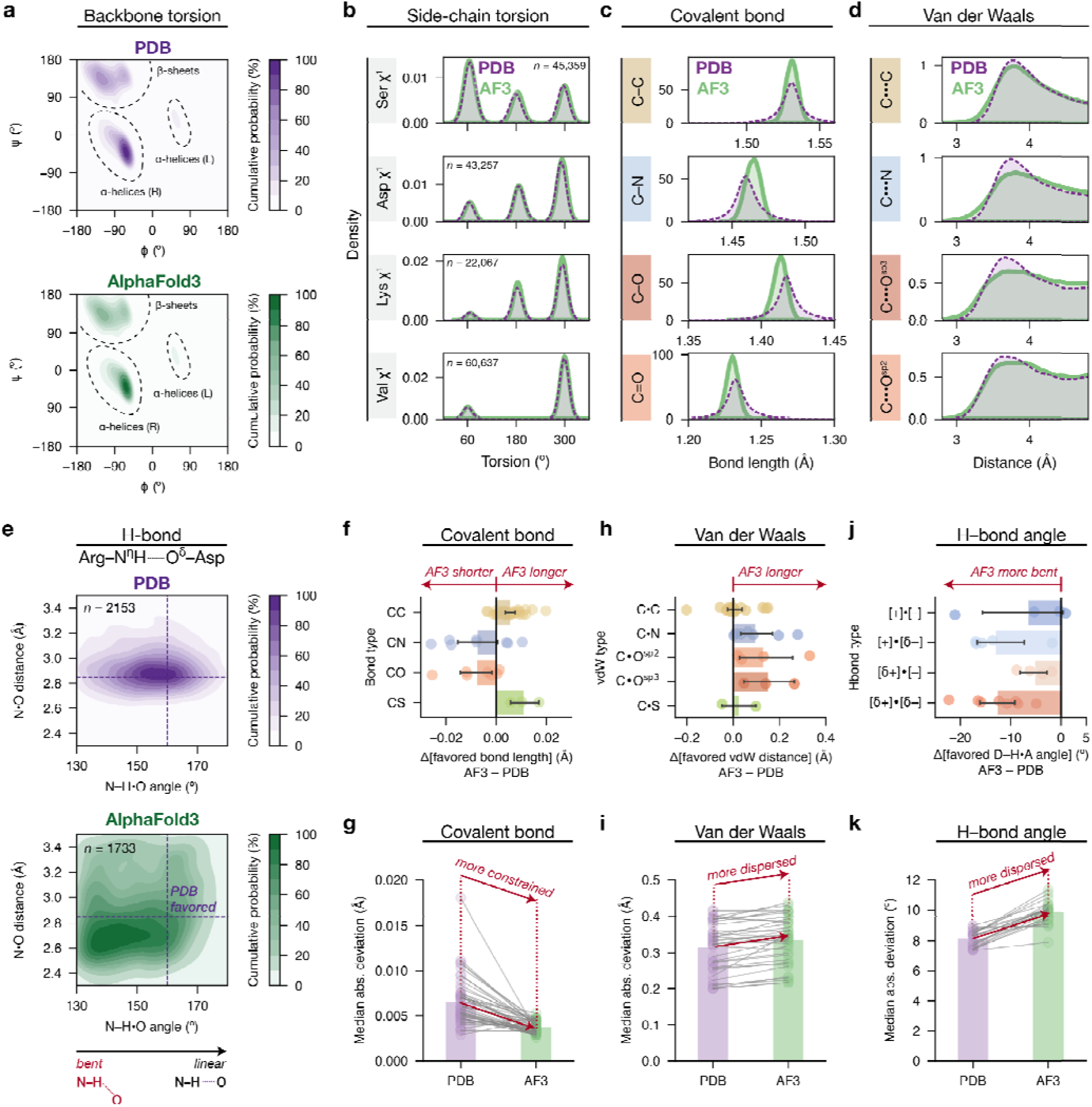
Biomolecular interaction energetics inferred by AlphaFold3. The analyses for AlphaFold2 and ESMFold gave similar results and are shown in Extended Data Figs. 1 to 3. **(a)** Probability density distributions of backbone and angles (*n* = 50,000 randomly sampled from 1, 097, 512 residues (not including glycine and proline residues; see Supplementary Fig. 1). The color gradients indicate the probability mass under the contour in percentages, with darker shades indicating more probable conformational states. **(b)** Examples of probability density distributions of side-chain torsion angles observed in the PDB (purple) and those from AlphaFold3 predictions (green). The distributions for other amino acids are shown in Supplementary Figs. 2. **(c)** Examples of probability density distributions of lengths of the specified covalent bonds. The distributions shown were taken from serine residues. The distributions for other amino acids are shown in Supplementary Fig. 5. **(d)** Probability density distributions of van der Waals distances for different interatomic contacts. Each distribution includes >75,000 van der Waals interactions (see sample sizes in Supplementary Table 3). **(e)** Example probability density distributions for hydrogen bond angles (donor–H•acceptor) and distances (donor•acceptor) for arginine•aspartate side-chain hydrogen bonds. The color gradients indicate the probability mass under the contour in percentage, with the darker shades indicating more probable conformational states. This distribution includes Arg•Asp residue pairs that form a single hydrogen bond between their side-chains (excluding bidentate hydrogen bonds) to ensure an unambiguous comparison. Additional hydrogen bond distributions are shown in Supplementary Fig. 9. **(f)** Differences in the most favored covalent bond lengths in AlphaFold3 predictions *versus* the PDB distributions. Each circle represents the peak of the length distribution for a specific covalent bond type based on the amino acid and atom identities; the bond types and values are given in Supplementary Table 4. **(g)** Differences in the spread of the distributions for covalent bond lengths in AlphaFold3 predictions *versus* the PDB distributions, quantified by the median absolute deviation (MAD). Each circle represents the MAD for a specific covalent bond type; data are provided in Supplementary Table 4. **(h)** Differences in the most favored van der Waals distances in AlphaFold3 predictions *versus* in the PDB distributions, separated by the atoms involved. Each circle represents the peak of the length distribution for a specific van der Waals interaction based on the amino acid and atom identities; the bond types and values are given in Supplementary Table 6. **(i)** Differences in the spread of the distributions for van der Waals distances in AlphaFold3 predictions *versus* in the PDB distributions, quantified by the median absolute deviation (MAD). Each circle represents the MAD for a specific type of van der Waals interaction, and the data are provided in Supplementary Table 6. **(j)** Differences in the most favored hydrogen bond angles in AlphaFold3 predictions *versus* in the PDB distributions, for the following hydrogen bond types: charge•charge ([+]•[–]), charge•dipole ([*δ*+]•[–], [*δ* –]•[+]) and dipole•dipole ([*δ* +]•[*δ* –]). Each circle represents the peak of the angle distribution for a specific hydrogen bond type, and the data are provided in Supplementary Table 7. Only hydrogen bonds with sp^2^ hybridized donor atoms were included so that the hydrogen positions and angles can be confidently calculated (*Methods*). **(k)** Differences in the spread of the distributions for hydrogen bond angles in AlphaFold3 predictions *versus* the PDB distributions, quantified by the median absolute deviation (MAD). Each circle represents the MAD for a specific type of hydrogen bond, and the data are provided in Supplementary Table 7.

AlphaFold3 partially captures the conformational preferences of non-covalent interactions. Its predicted van der Waals interactions follow Lennard-Jones-like energetics, where a specific inter-atomic distance is preferred for each interaction type and shorter-than-ideal distances are increasingly disfavored (Fig. 2d, Supplementary Fig. 8 and Supplementary Table 6). Hydrogen bonds in AF3 have particular distance and angle preferences, although these preferences differ from those in the PDB training set (Fig. 2e, Supplementary Fig. 9 and Supplementary Tables 7 to 8).

Thus, AlphaFold3 has extracted conformational and energetic properties of molecular interactions. Nevertheless, notable discrepancies remain, which we describe and quantify below.

### Discrepancies in the conformational distributions derived from AlphaFold3

The peak(s) in each conformational distribution represents the most favored state(s) and the breadth corresponds to the range of conformers and their relative probabilities. Differences in the peaks and the breaths of the distributions when compared to the PDB would correspond to inaccuracies in the conformational preferences learned by the models and an inability to reproduce at least a subset of the experimentally observed states. To evaluate differences between the AlphaFold3 and PDB distributions, we compiled distributions for each interaction type and compared the peaks and the dispersions of the distributions to that from the training data. We identified differences for nearly all covalent and non-covalent interactions (Fig. 2f to h).

We found that the favored covalent bond lengths differ, on the order of ∼0.01–0.03 Å for multiple bond types between the AlphaFold3 and PDB distributions, with the direction of change dependent on atom and bond type (Fig. 2f and Supplementary Table 4). While the differences are subtle, bootstrap analyses showed that they are highly significant (Supplementary Fig. 7). Similarly, covalent bond angles showed deviations on the order of ∼1–3° (Supplementary Fig. 6 and Table 5). In addition, the AlphaFold3 distributions were consistently narrower than those derived from the PDB distributions, as quantified by lower median-absolute-deviation values (Fig. 2g and Supplementary Table 4 to 5).

Larger differences were found for non-covalent interactions. For van der Waals distances and hydrogen bond lengths, the peaks differ by ∼0.1 Å or more for side-chain interacting partners for AlphaFold3 relative to the PDB peaks, except for C•C van der Waals interactions, where the peaks agreed (Fig. 2h, Supplementary Fig. 8 and Table 6). AlphaFold3 hydrogen bonds adopt more bent geometries compared to the linear hydrogen bonds found in the PDB (Fig. 2j and Supplementary Table 7) and in small molecule structures and quantum mechanical calculations^38–40^. The AlphaFold3 non-covalent distributions were broader than those from the PDB—*i.e.*, these interactions are more varied in AlphaFold3 predictions than in the PDB (Fig. 2i and k and Supplementary Tables 6 to 7). The increased breadth, in addition to the altered peak positions, was particularly pronounced for hydrogen bonds (Supplementary Fig. 6).

Overall, differences were found within and across different types of interactions, and the direction of change is specific to residue and atom types, indicative of systematic errors rather than random noise. The observed discrepancies could arise from how the model is constructed and trained, or from limitations in the training data—*e.g.*, from errors in the low resolution structures in the PDB, from insufficient data, or from both. As covalent bonds are typically constrained when solving X-ray structures^41^ and are highly abundant, their distortions more likely arise from issues in the model rather than the data. To investigate the consequences of these conformational biases on model performance, we next examined how they manifest in the prediction of individual interactions in each protein structure and of conformational ensembles.

### Widespread prediction errors of side-chain interactions

Protein structure––more fundamentally, the probability of a conformational state in an ensemble––is determined by the sum of the energetics from all the interactions that are present (Fig. 1a). The discrepancies in the overall conformational preferences identified above could alter the balance of forces so that the model predicts conformational states with different sets of interactions compared to the corresponding PDB structures. Alternatively, the discrepancies may have less severe consequences, maintaining the same set of interactions but with some geometric distortions.

To distinguish between these possibilities and learn more about the models’ capabilities and limitations, we first compared each bond torsion angle between AlphaFold3 predictions and PDB structures, as these rotations underlie changes in non-covalent interactions as well as more extensive rearrangements. We then evaluated the energetics consequences of the identified torsion deviations as well as the accompanying changes in hydrogen bonds and van der Waals interactions. This systematic approach allowed us to parse individual interactions, providing a multi-dimensional physics-based breakdown of the physical terms that underlie structures.

Our comparisons focused on buried residues (relative solvent accessibility <25%, 410,000 residues and 2.2 million interactions; Supplementary Table 3) as their conformations are determined by the sum of their interactions with surrounding protein groups and so can be used to evaluate the energetic interplay of these interactions. Additionally, buried residues are typically the most confidently modeled from the X-ray diffraction data^42,43^, thus minimizing errors from X-ray modeling and refinements in our comparisons.

### Assessing backbone and side-chain bond torsions

Among 1,277,419 backbone torsion angles, 99% of the *ψ* and *ϕ* angles match their PDB references within 30° (Fig. 3a). Correspondingly, 91% of the backbone amide groups are predicted in the correct secondary structure (Supplementary Table 9). The agreement is higher for residues forming alpha helices and beta sheets (97% and 96%) than for residues outside of these secondary structures (85%; Supplementary Table 9). High accuracy of backbone torsion angles is consistent with the excellent performance of AlphaFold models in predicting protein folds in prior evaluations based on backbone or C atom RMSDs^6^.

**Fig. 3.**
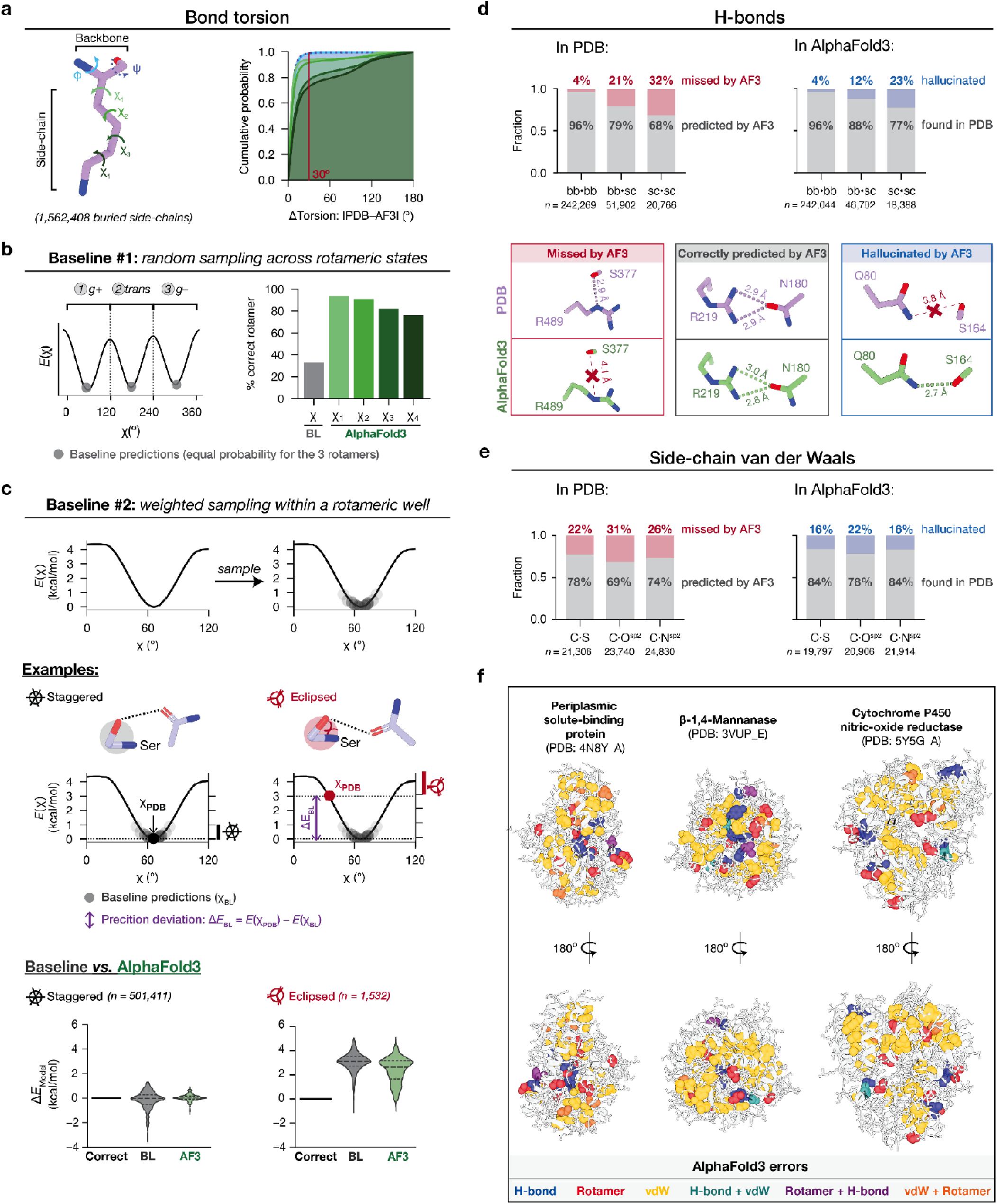
One-to-one comparisons for interactions in AlphaFold3 predictions *versus* the corresponding PDB structures. See Extended Data Figs. 4 to 5 for AlphaFold2 and ESMFold results. **(a)** Left: definition of backbone and side-chain torsion angles, using a lysine residue as an example. Right: Cumulative distribution for side-chain torsion angle differences between AlphaFold3 predictions and the corresponding PDBs. **(b)** Prediction accuracy for rotameric states of AlphaFold3 compared to a context-free baseline model that randomly samples torsion angles according to the PDB distribution. Left: the baseline model samples three rotameric states that correspond to the local energy minima (*gauche-, trans, gauche+*) with near-equal probabilities; thus, the model predicts the correct rotamer ∼1/3 of the time. The three rotameric states are mapped on an energy function *E*(χ), obtained by inverting the PDB torsion angle distribution *f*(*χ*) using the Boltzmann relationship (*Methods*). Right: percentage of correct side-chain rotameric states predicted by AlphaFold3 compared to the baseline. **(c)** Comparison of AlphaFold3 to a second context-free baseline model in predicting staggered and eclipsed bonds. Top: baseline model that samples within a rotameric state. The low-energy staggered conformations are more likely sampled than the eclipsed conformations. Middle (examples): expected baseline model accuracy in predicting staggered and eclipsed conformations reported by the energetic deviation of the PDB *versus* AlphaFold-predicted torsional state [Δ*E*_BL_ = *E*(χ_PDB_) - *E*(χ_model_)], obtained by mapping the torsion angles onto the PDB torsion energy function *E*(χ). For a staggered PDB bond (*e.g.*, the serine example shaded in grey), the baseline model predicts the correct energy (Δ*E*_BL_ ∼ 0) as it most likely samples a staggered conformer. For an eclipsed conformer constrained by nearby molecular interactions in the PDB (*e.g.*, the serine example shaded in red), the baseline model is expected to give a positive deviation with the deviation scaling proportionally with the energy of the torsion angle [*e.g.*, Δ*E*_BL_ ∼= 2.5 kcal/mol when *E*(χ_PDB_) = 2.5 kcal/mol]. Bottom: AlphaFold3 (light green) compared to baseline (gray) for energy deviations from staggered (bottom left) and eclipsed (bottom right) PDB references. For reference, a theoretical model that predicts all bonds correctly would have Δ*E*_correct_ = 0 and is shown as the black lines. **(d)** Comparisons of buried backbone•backbone (bb•bb), backbone•side-chain (bb•sc), and side-chain•side-chain (sc•sc) hydrogen bonds in the PDB structures and in AlphaFold3 predictions. The gray bars indicate the numbers of hydrogen bonds in common in the PDB structure and the corresponding AlphaFold3 prediction (“correctly predicted by AF3”). For the PDB hydrogen bonds, the fraction that were not found in the AlphaFold predictions are shown as red bars (“missed by AF3”); for the AlphaFold3 hydrogen bonds, the fraction that were not found in the PDB structures are shown as blue bars (“hallucinated”). Representative structures for each group are shown below [PDB: 2W3P_B, 1OFL_A and 1KGN_C, from left to right (purple) and their corresponding AlphaFold3 predicted structures (green)]. **(e)** The same comparisons as in panel d for side-chain•side-chain van der Waals interactions. These interaction distances are within 0.4 A of the ideal or “peak” value (*Methods*), so that the “missing” category includes both interactions that are too short (clashes) or too long (no interactions formed). Steric clashes represent a small (0.11%) percentage of the interactions (see also Supplementary Fig. 8). **(f)** Examples of prediction errors in buried side-chain interactions mapped on protein structures for the misassignment of rotameric states, hydrogen bond and van der Waals partners, and combinations of these errors, shown in different colors. Buried residues with correctly assigned interactions are shown as white sticks.

Bond torsion accuracy is lower for side-chains, and decreases for bonds extending further from the backbone, with 94% to 77% of the predicted *χ*^1^ to *χ*^4^ torsion angles falling within 30° of the corresponding PDB structure (Fig. 3a). The decreasing accuracy may arise from the larger conformational space occupied by longer side-chains, from the increased number of interactions they can form, or from their lower abundance and thus less training opportunity.

### Side-chain torsion deviations indicate altered energetic balances in AlphaFold

The favored bond torsions of each buried residue are determined by local molecular forces, and differences in predicted torsions indicate altered energetics of the surrounding environment. Rotameric side-chains can occupy three energetic minima or “wells” (*gauche–*, *trans*, and *gauche+*)^44^, and rotation to an alternative well can result in steric clashes and accompanying rearrangements of the surrounding atoms. Rotation within the same rotameric wells, while giving smaller RMSDs than rotation to a different well, can have substantial energetic consequences––*e.g*., a 40° torsional rotation from staggered to partially eclipsed is destabilizing by ∼3 kcal/mol, corresponding to ∼160-fold difference in probability at room temperature (Fig. 1b). Thus, the eclipsed conformers provide a sensitive test for the model’s ability to accurately capture energetic interplay. Nevertheless, eclipsed bonds are rare and their effects would be masked by typical comparisons that combine all bonds together (*n* = 1532 buried eclipsed bonds, compared to *n* = 501,411 buried staggered bonds, as described below). To overcome this limitation, we evaluated the predicted bond torsions in two stages: (1) across rotameric wells, and (2) within each rotameric well. To quantify the model’s ability to capture the local energetic balance, we compared model predictions to those made by “baseline” models that contain no information about surrounding forces.

To predict rotameric wells, the constraint-free baseline model randomly samples between the *gauche–*, *trans*, and *gauche+* conformers, and thus predicts the correct rotameric state one-third of the time (Fig. 3b, baseline #1). AlphaFold3 gave a substantial improvement from this baseline, correctly predicting the rotameric state for 94%, 92%, 78%, and 73% of the *χ*^1^, *χ*^2^, *χ*^3^ and *χ*^4^ bonds (Fig. 3b).

For the second stage of testing, we selected bonds where AlphaFold3 predicted the correct rotameric well to test the model’s ability to predict the preferred torsion angles within each well. We compared AlphaFold3 to a second baseline model, this time randomly sampling torsion angles within each well, weighted by the probability of occurrence of each angle in the PDB reference dataset (Fig. 3c, upper, baseline #2) so that the probability of sampling an eclipsed bond would be much lower than sampling a staggered bond, as it would be without any forces from the protein context. We used the Boltzmann relationship to convert probabilities into knowledge-based energies^45,46^ to provide a simple evaluation metric in kcal/mol (*Methods*). Because the baseline model most frequently samples the low-energy, staggered states, it is expected to give at most small deviations for staggered bonds in the PDB (Δ*E*_BL_ ≈ 0; defined in Fig. 3c, middle). By contrast, high energy, eclipsed states are rarely sampled by the baseline model and thus unlikely to be predicted correctly by chance––*e.g*., an average deviation of Δ*E*_BL_ = 3 kcal/mol is expected for the partially eclipsed state shown in Fig. 3c (middle) as the model preferentially predicts staggered states.

For staggered states that are within 1 kcal/mol of the minimum, the baseline model and AlphaFold3 both gave Δ*E* values centered around zero (Fig. 3c, lower). AlphaFold3 gave a tighter distribution, indicating improved accuracy over the random baseline model. To test eclipsed states, we selected bonds that are 2.5 to 4.0 kcal/mol higher in energy than the minimum in the PDB structures. The baseline model gave the expected deviation of Δ*E*_BL_ = 3.1 kcal/mol (SD = 0.7; Fig. 3c, lower) for these bonds. AlphaFold3 also deviated from the PDB, with average prediction deviations of Δ*E* = 2.4 (SD = 1.1) kcal/mol. This deviation is substantial but less than that for the baseline model deviation (Fig. 3c; Extended Data Fig. 4 and 5). The AlphaFold3 predictions captured 6% of the eclipsed bonds within 0.5 kcal/mol of the PDB values, suggesting that certain structural contexts may be reproduced more accurately but that most are not for these locally strained states.

Overall, AlphaFold3 outperformed the baseline model in predicting the correct rotameric wells, where errors would involve relatively large movements and steric factors. Nevertheless, the balance of forces responsible for favoring states within a particular rotamer was not well captured. This limitation may arise because bond rotation within a well is highly sensitive to the accuracy of the local energetic balance, despite small changes in atomic positions and thus gives only a small penalty from the training metrics used in model development.

### AlphaFold3 misassigned about one-third of side-chain non-covalent interactions

Rotations of the side-chain bonds are expected to perturb the non-covalent interactions they are involved in. To test the consequence of torsional deviations in non-covalent interactions, we performed one-to-one comparisons for hydrogen bonds and van der Waals interactions. Because these interactions involve residue pairs, the tests were performed in two steps: (1) determining whether the same residue pairs form interactions in AlphaFold3 and PDB structures, and (2) quantifying the extent of geometric deviations of the interactions that were in common.

Our comparisons included ∼440,000 hydrogen bonds where the donor heavy atom is sp^2^ hybridized, as we could unambiguously assign the hydrogen position for this subset of hydrogen bonds (Supplementary Table 3). Nearly all (96%) backbone•backbone hydrogen bonds were predicted between the same atom pairs of the same residues as seen in the PDB. The accuracy decreased substantially for hydrogen bonds involving side-chains—with 21% and 32% of the backbone•side-chain and side-chain•side-chain hydrogen bonds, respectively, predicted to form between incorrect partners by the AlphaFold3 models (Fig. 3d, left). AlphaFold3 also hallucinated hydrogen bonds such that 4%, 12% and 23% of the backbone•backbone, backbone•side-chain and side-chain•side-chain hydrogen bonds, respectively, were not present in the corresponding PDB structures (Fig. 3d, right).

Analogous comparisons for side-chain van der Waals interactions were performed by identifying carbon•heteroatom pairs within 0.4 Å of their ideal van der Waals distances (*Methods*). These comparisons revealed similar trends as observed for hydrogen bonds. Across different atom types, from 15% to 31% of the van der Waals interactions in the PDB are missing in AlphaFold3 predictions, and 14% to 22% of the model-predicted van der Waals contacts were hallucinated (Fig. 3e).

Even when AlphaFold3 predicted the correct interaction pairs, substantial geometric deviations were found. About 39% of the side-chain distances for correctly-predicted hydrogen bond donor•acceptors deviated by more than 0.2 Å from the corresponding PDB distances, and 32% were at least 20° more bent (Extended Data Fig. 6a). Similarly, among the ∼130,000 correct van der Waals pairs, about 32% of the AlphaFold3 predictions deviated >0.2 Å from the PDB (Extended Data Fig. 6b).

As the structure predictions correspond to the unliganded, apo states, while the PDB reference set contains both apo and ligand-bound structures, we assessed whether the errors might arise from the subset of comparisons made between predicted apo structures and ligand-bound references. Including only apo reference structures in the comparisons to the model predictions gave similar error rates for each interaction type (Extended Data Fig. 7), suggesting that the subset of ligand-bound structures is not predominantly responsible for the observed error rates. Nevertheless, a priority for developing next-generation models will be testing and removing any errors that might emerge from a mixture of apo and ligand-bound structures in the training set; separately treating apo and bound structures would also be needed for the models to learn ligand binding interactions so that they can be accurately predicted.

### Incorrect predictions are distributed throughout the structures and across the dataset

To determine whether there are idiosyncratic features associated with the errors that could help pinpoint specific model limitations, we tested whether errors are enriched in certain regions of the proteins or associated with certain types of protein structure.

Mapping the incorrect predictions onto each protein structure, we observed that incorrect interactions appeared to be distributed throughout each structure rather than clustered in particular regions (Fig. 3f). Quantitatively, the extent of clustering of incorrectly predicted residues within each protein was indistinguishable from that of randomly selected residues (Extended Data Fig. 8). The randomly distributed rather than clustered errors suggest multiple individual cases of inaccurate energetic balances, rather than a single or small number of epicenters that propagate errors.

We next determined the propensity for different groups of proteins across the dataset to be more or less prone to errors. Incorrect interactions were found in nearly all structures across different structural classes, and the number of incorrect interactions in each structure scaled with protein size, suggesting that the prediction errors were not associated with specific proteins (Extended Data Fig. 9).

In addition, the prediction errors were not identified from AlphaFold confidence scores, as residues involved in incorrect interactions gave average pLDDTs of 92 (SD = 8) (Extended Data Fig. 10). Altogether, these results indicate system-wide, non-idiosyncratic side-chain prediction errors that are not identified as suspect by the model.

### AlphaFold sampling produced highly restricted ensembles compared to experimental ensembles

Conformational ensembles are the most direct emergent properties of energy landscapes and provide an independent test for the extent to which the models reproduce protein energetics. Moreover, ensemble generation is a rapidly growing frontier of predictive models^47–50^, an important direction as ensembles provide the states and probabilities that are ultimately needed to provide quantitative prediction of function^51–53^ (Fig. 1a).

AlphaFold3’s diffusion-based architecture is well suited for ensemble generation because each reverse-diffusion trajectory samples from an internal probability distribution^9,54^. To evaluate AlphaFold3’s ability to generate accurate ensembles, we curated reference experimental datasets collected by non-cryogenic multi-temperature (MT) X-ray crystallography, as these data reveal a wider range of conformational states than X-ray structures determined at cryogenic temperatures^55–60^. We collected high-resolution MT datasets (1.0–2.2 Å) for nine proteins that were determined above 200 *K* and had been refined into multi-conformer models (Fig. 4a and Supplementary Table 10). We repeatedly sampled from AlphaFold3 for each of these sequences initializing the predictions with random seeds, and we compared the resulting bond torsion distributions to the corresponding MT datasets.

**Fig. 4.**
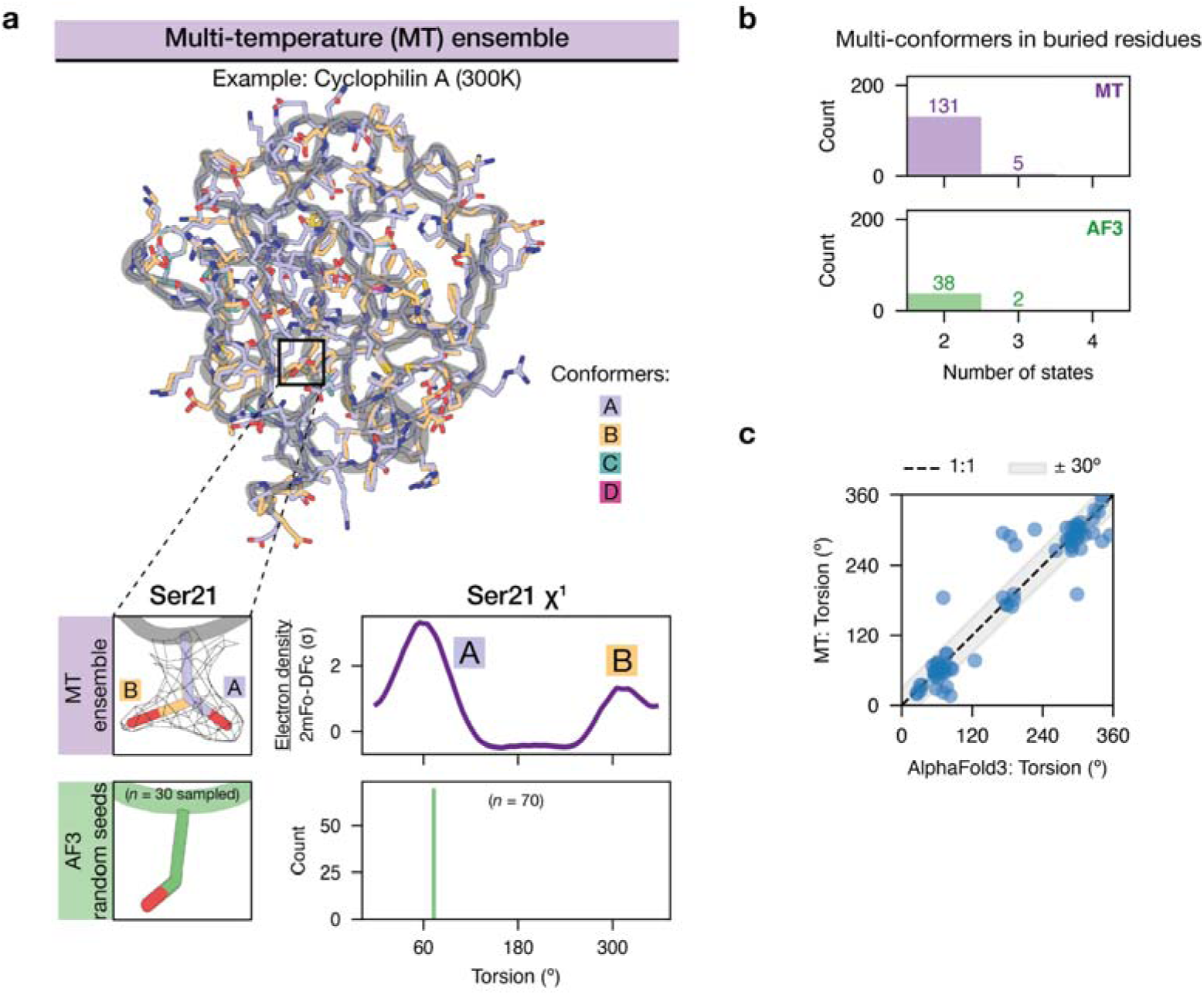
Comparisons of multi-temperature (MT) crystallographic models and AlphaFold3 samples generated by random seeding. **(a)** Top: multi-conformer ensemble model of Cyclophilin A refined from an X-ray dataset collected at 300 *K* (PDB: 4YUM; resolution = 1.5 Å). Bottom: an example of the comparison performed for the multi-conformer side-chain of Ser21 from Cyclophilin A. The electron density map around Ser21 (2mFo-DFc; purple line) shows two conformers corresponding to modeled A and B conformers. The torsion distribution for this residue from the AlphaFold3 sampling is shown below in green. **(b)** The number of torsion angles with multiple states in the MT models *vs.* from AlphaFold3 sampling found in buried residues (<0.25 relative solvent accessible area, *n* = 1363 total buried bonds). **(c)** One-to-one comparisons for the torsional angles predicted by AlphaFold3 and found in the MT model for buried residues where AlphaFold3 predicted multiple conformational states (*n* = 82 conformers from 40 sidechains). The dashed line indicates 1:1 or perfect agreement, and the gray shaded area indicates a range of ±30°.

We found 136 side-chain bonds that clearly occupy two or more conformational states (a total of 270 conformers) across the 1363 buried side-chain bonds in the MT multi-conformer models (Fig. 4b). For most of the side-chains exhibiting alternative conformers, AlphaFold3 sampling generated only a single conformer (96 of 136) with limited to no variations in the sampled distributions, suggesting an overly restricted probability landscape in the model (Fig. 4a to b). The same sampling procedures using AlphaFold2 generated similarly highly restricted distributions (Supplementary Fig. 10). Nevertheless, for the subset when multi-conformer states were predicted by AlphaFold3, we found high precision in predicting the multi-conformers found in the MT models (38 out of 40; only 2 hallucinated), and the majority of these conformers are in close agreement (Fig. 4c).

Overall, these results suggest that the AlphaFold models sample from highly restricted distributions, resulting in nearly deterministic predictions for the majority of the residues even when using random seeds. Emerging ensemble-generation models that perform ensemble-reweighting may be able to improve the number of alternative states and the accuracy of those states^e.g.,50^. Our physics-based evaluation framework, combined with high-quality experimental ensemble models, provides the foundation for systematically assessing these emerging AI models to guide their improvement.

### Comparisons across multiple models revealed limitations and strengths of each model

Structure prediction models have been developed using different model architectures and training strategies. To test the generalizability of our evaluation framework and to determine the strengths and weaknesses of different models, we also evaluated AlphaFold2^6^ and ESMFold^7^, two widely used models with different structure and sequence modules than AlphaFold3.

### Limited capture of the conformational preferences of noncovalent interactions

AlphaFold2 and ESMFold both showed biases in the conformational preferences for noncovalent interactions, as was observed for AlphaFold3 (Fig. 5a and Extended Data Figs. 1 to 3). ESMFold was shown to give less accurate overall structures than the AlphaFold models in prior evaluations^61^. We found that ESMFold reproduced the Ramarchandran map and the torsion angle preferences with similar accuracy as AlphaFold2 and AlphaFold3 (Supplementary Figs. 1 and 4), but it showed highly dispersed distributions and large deviations in the preferred conformations for noncovalent interactions, compared to the AlphaFold models (Fig. 5a and Supplementary Figs. 8 and 9). Correspondingly, the ESMFold error rates for noncovalent interactions are substantially higher, reaching ∼60% for sidechain hydrogen bonds (Fig. 5b and Extended Data Fig. 5). These observations are consistent with prior evaluations and suggested that its major sources of errors are non-covalent pairwise interactions.

**Fig. 5.**
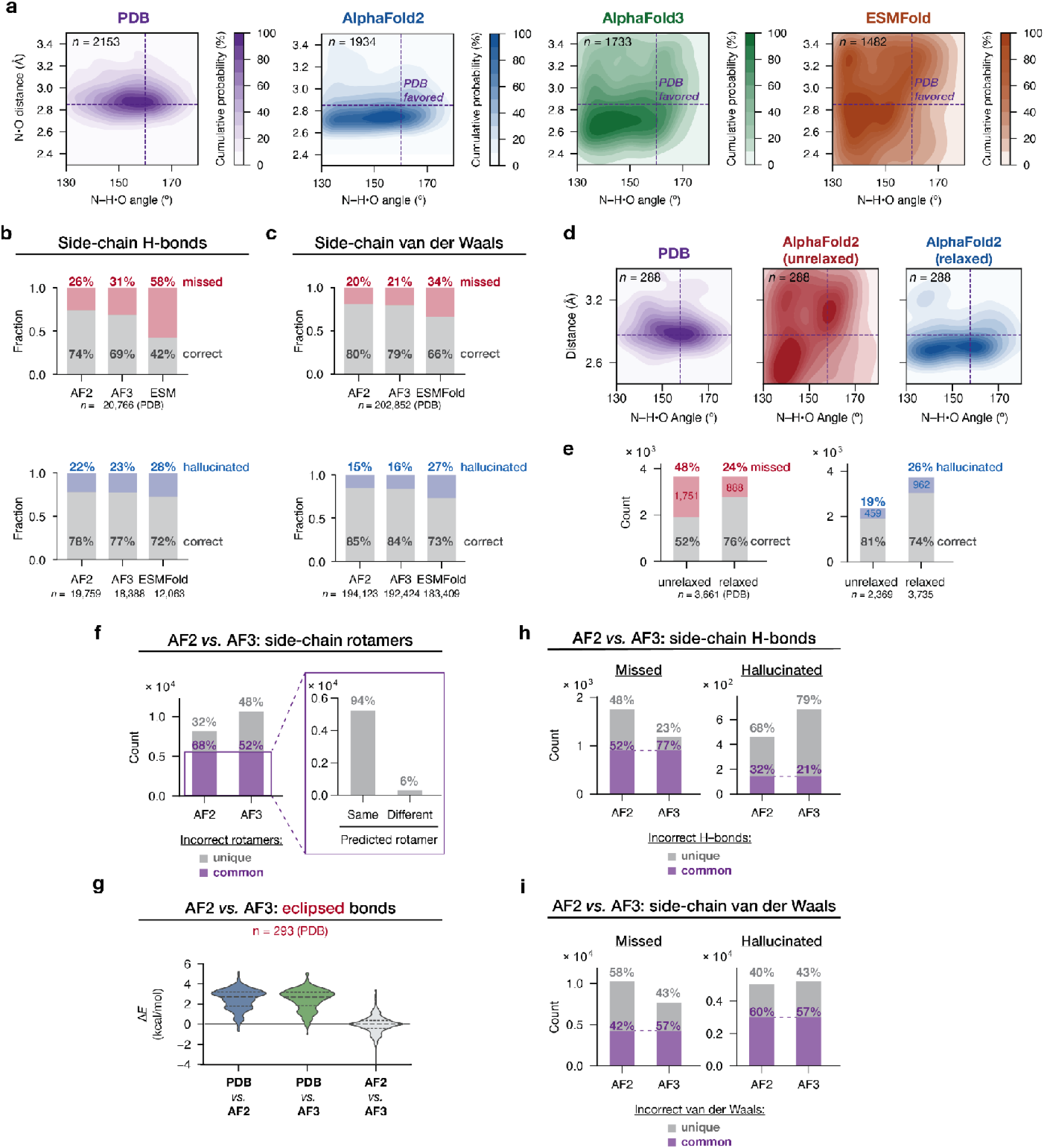
Comparison of AlphaFold2, AlphaFold3 and ESMFold performance in predicting interactions. **(a)** Probability density distributions for hydrogen bond angles (donor–H•acceptor) and distances (donor•acceptor) for an example—arginine•aspartate side-chain hydrogen bonds—in the PDB (purple), and in the AlphaFold2 (blue), AlphaFold3 (green) and ESMFold (orange) predictions. The color gradients indicate the probability mass under the contour in percentage (10% each contour), with the darker shades indicating more probable conformational states. This distribution includes Arg•Asp residue pairs that form a single hydrogen bond between their side-chains (excluding bidentate hydr gen bonds) to ensure an unambiguous comparison (*Methods*). Additional hydrogen bond distributions are shown in Supplementary Fig. 9. **(b)** Comparison of the accuracy of side-chain hydrogen bonds predicted by the three models. The gray bars indicate the fraction of hydrogen bonds in common between the PDB structure and the corresponding model prediction (“correct”). In the upper subpanel, the fraction that were found in the PDB but not found in model predictions are shown as red bars (“missed”), and the total number of hydrogen bonds (*n*) correspond to the number of hydrogen bonds found in the PDB structures. In the lower panel, the fraction of hydrogen bonds that were found in model predictions but not found in the PDB structures are shown as blue bars (“hallucinated”), and the total number of hydrogen bonds found for predictions in each model is annotated below the bar. **(c)** Comparison of the accuracy of side-chain van der Waals interactions predicted by the three models. The color scheme used is the same as in (b). **(d)** Probability density distributions for hydrogen bond angles (donor–H•acceptor) and distances (donor•acceptor) for Arg•Asp side-chain hydrogen bonds in the PDB (purple), in the unrelaxed AlphaFold2 (red) and relaxed AlphaFold2 (blue) structures. There are 288 of these hydrogen bonds in the unrelaxed AlphaFold2 dataset (containing only apo structures); the PDB and relaxed AlphaFold2 dataset are larger but they were resampled to the same sample size (*n* = 288) for direct comparisons. The color gradients indicate the probability mass under the contour in percentage (10% each contour), with the darker shades indicating more probable conformational states. Additional hydrogen bond distributions are shown in Supplementary Fig. 11. **(e)** Comparison of the accuracy of sidechain hydrogen bonds before and after AMBER relaxation. The grey bars indicate the count of hydrogen bonds in common between the PDB structure and the corresponding model prediction (“correct”). In the upper subpanel, the number of hydrogen bonds that were found in the PDB but not found in model predictions are shown as red bars (“missed”), and the total number of hydrogen bonds (*n*) correspond to the number of hydrogen bonds found in the PDB structures. In the lower panel, the number of hydrogen bonds that were found in model predictions but not found in the PDB structures are shown as blue bars (“hallucinated”), and the total number of hydrogen bonds for each model is annotated below the bar. **(f)** Rotameric states errors in AlphaFold2 and AlphaFold3. Left: the number of incorrect rotamers that were common (purple) and unique (grey) errors. Right: within the shared subset of errors, number of common *versus* different predicted (incorrect) rotameric states. **(g)** Energetic deviation (Δ*E*) for eclipsed side-chain bonds torsions for PDB conformers to the AlphaFold2 predictions; for PDB conformers *versus* AlphaFold3 predictions; and for AlphaFold2 *versus* AlphaFold3 predictions. **(h)** Side-chain hydrogen bond errors in AlphaFold2 and AlphaFold3 that were common (purple) and unique (grey). Left: hydrogen bonds identified in the PDB structures but not in the corresponding model predictions. Right: hydrogen bonds hallucinated in each model prediction. **(i)** Side-chain van der Waals errors in AlphaFold2 and AlphaFold3 that were common (purple) and unique (grey). Left: van der Waals interactions identified in the PDB structures but not in the corresponding model predictions. Right: van der Waals interactions hallucinated in model predictions.

AlphaFold2 produced tighter distributions and more accurate conformational preferences than AlphaFold3 and ESMFold (Fig. 5a, Extended Data Fig. 3, and Supplementary Figs. 8 and 9). AlphaFold2 also showed a slightly higher accuracy in reproducing the noncovalent interaction pairs in the PDB, with 26% missed sidechain hydrogen bonds, compared to the 31% and 58% in AlphaFold3 and ESMFold, respectively (Fig. 5b to c). A simple model for AlphaFold2’s higher accuracy is that the AMBER force field relaxation step performed at the end of the AlphaFold2 pipeline but not the other two models improved its performance. We therefore asked whether this relaxation step contributed to higher accuracy, and if so, to what extent.

### Force field relaxation alters conformational preferences in AlphaFold2 and partially rescued misassigned noncovalent interactions

To determine the effects of the AMBER force field relaxation step on AlphaFold2 predictions, we selected the apo structures in our PDB dataset (*n* = 1056) and performed AlphaFold2 predictions with no relaxation, and then compared the resulting structures to the relaxed AlphaFold2 predictions. We found large differences in the overall conformational preferences. Without relaxation, the conformational distributions of sidechain hydrogen bonds are dispersed, similar to those produced by AlphaFold3 and ESMFold (Fig. 5d and Supplementary Fig. 11), indicating that force field relaxation is responsible for the higher accuracy of AlphaFold2.

To quantify the extent to which relaxation rescued misassigned side-chain interaction pairs, we performed the one-to-one comparisons for relaxed *versus* unrelaxed AlphaFold2 structures. Relaxation of the AlphaFold2 structures led to a substantial decrease—from 48% to 24% missing side-chain•side-chain hydrogen bonds (Fig. 5e). Nevertheless, the number of hallucinated hydrogen bonds increased after relaxation—from 19% to 26% of the total hydrogen bonds found in the predicted structures (Fig. 5e). Smaller effects were observed for van der Waals interactions, where relaxation rescued 8% of the 30% missing interactions and also 6% of the hallucinated interactions (Supplementary Fig. 12).

Carrying out the same relaxation on the AlphaFold3 structures gave the same trends as for AlphaFold2 but a smaller beneficial effect—a decrease from 32% to 22% missing side-chain•side-chain hydrogen bonds; once relaxed, the AF2 and AF3 structures gave similar amounts of errors (Supplementary Fig. 12).

Overall, our comparisons revealed a considerable effect of a simple force field relaxation step in modifying the overall conformational preferences of interactions, including partial rescue of incorrect sidechain interactions. However, even with relaxation, ∼20% of sidechain interaction errors persisted for both AlphaFold models. Resolving these errors may require more extensive rearrangements than relaxation can provide and/or improvements in the accuracy of force fields.

### Common prediction errors by AlphaFold2 and AlphaFold3

AlphaFold3 employed a different structural module than AlphaFold2, which could lead to distinct strengths and weaknesses of the two models. We therefore compared the errors made by AlphaFold2 and AlphaFold3, prior to effects from relaxation, to identify potential differences. Of the 8171 and 10,661 mis-assigned side-chain rotamers in unrelaxed AlphaFold2 and AlphaFold3 predictions, respectively, most errors (5551) were in common. Intriguingly, unrelaxed AlphaFold2 produced slightly less errors in predicting rotamerics states than AlphaFold3, in contrast to its higher error rates for non-covalent interactions. This behavior may arise because its structural module employs a “global rigid body frame” representation and generates structures via rotation of the torsion angles^6^, which may be more effective in learning and reproducing rotameric preferences than pairwise non-covalent interactions, whereas the all-atom diffusion-based structural module of AlphaFold3 may better capture non-covalent interactions.

For nearly all the common errors (94%), the models predicted the same incorrect rotameric states (Fig. 5f, right). In addition, AlphaFold2 and AlphaFold3 also made near-identical (and mainly incorrect) predictions for the eclipsed, high-energy bonds found in the PDB (Fig. 5g).

Mis-assigned noncovalent interactions also tended to be common in AlphaFold2 and AlphaFold3 predictions. PDB hydrogen bonds and van der Waals interactions that were missing in AlphaFold3 predictions were also mainly missing in AlphaFold2 predictions (77% and 57%), with AlphaFold2 missing more interactions (Fig. 5h). Similarly, a large fraction of hallucinated hydrogen bonds and van der Waals interactions were common for AlphaFold2 and AlphaFold3 (21% and 57%, respectively; Fig. 5i). After relaxation, while the errors made by AlphaFold2 were partially corrected, the remaining errors were still largely in common with AlphaFold3 (Supplementary Fig. 13).

Overall, we found a preponderance of common errors made by AlphaFold2 and AlphaFold3, regardless of force field relaxation. This commonality could arise from using AlphaFold2-Multimer predictions to augment the training set for AlphaFold3^9^ or from a higher complexity of the energetic balance of certain sites that makes them more challenging to predict, such as in the case of eclipsed bonds. Identification of these persisting errors underscores the value of a systematic evaluation approach in revealing specific limitations of each model to guide the next iteration of model development.

### Conclusions and Implications

Effective evaluations drive model development by identifying the capabilities and limitations of a model—what the model can reliably do, what it cannot do, and what improvements are needed to expand its capabilities. Current evaluations of deep learning biomolecular models often reduce the complex features of a structure to distance-based metrics; while valuable for their simplicity and conciseness, these metrics are limited in their ability to determine what the models can reliably predict and in their ability to identify how to improve the models. We developed a systematic and generalizable evaluation framework that overcomes these limitations, providing a multifaceted assessment of structure prediction in the natural language of structure and function—molecular interactions and their energetics. Our evaluation revealed previously unknown capabilities and limitations of current structure prediction models. The models extracted basic energetic properties for molecular interactions, including covalent bonds, bond torsions, and van der Waals interactions. Nevertheless, the energetic rules learned by the models were flawed, leading to pervasive errors in side-chain interactions in the predicted structures. These incorrect interactions and interaction networks amount to nearly 30% of all side-chain hydrogen bonds and van der Waals in the AlphaFold models and 60% in ESMFold. These errors may be responsible for the limited applicability of AlphaFold predictions for downstream tasks such as ligand docking and mutational effects^22,34,62,63^ (Supplementary Table 1). All three models evaluated here suffer from limitations in reproducing pairwise non-covalent interactions, underscoring a general deficiency that is not unique to a certain model architecture.

Corrections for these interaction errors can be attempted via force field relaxation or molecular dynamics simulations of the predicted structures, yet these “corrections” also require rigorous evaluations via the framework presented here or an analogous tool. We found that relaxation rescued many of the errors, but >20% errors remained and additional hallucinated interactions were introduced. Future model development can apply the same approaches to determine the extent to which different model architecture, training data, and subsequent auxiliary steps improve model performance.

The conformational biases and the accompanying prediction errors we identified in the AI models might be overcome by improvements at two levels: (1) using higher quality data in model training and (2) using training approaches that extract physical information more efficiently from the given data. Ensemble data obtained from MT X-ray, other perturbative X-ray approaches^9,64^, and cryo-electron microscopy^65^ at high-resolution provide rich information about energy landscapes and may help models learn energetics. Further, incorporating physical knowledge, such as adding more comprehensive terms that describe interactions and their energetics in the training objectives and weighing these terms more heavily, may help minimize overfitting to distance-based metrics. Most generally, carrying out model development in a stepwise fashion while rigorously evaluating the impact of each step on model performance will allow future models to achieve the greatest predictive power.

The multifaceted, physics-grounded evaluation framework developed here can be applied to any model that generates atomistic structural information and can be readily expanded to include interactions involving non-amino acid groups (nucleic acids, solvent, and ligands). This generalizable evaluation approach will aid the evolution of deep learning models from memorization and interpolation of structural data to the extraction of fundamental physics universal to all biomolecular behaviors. Ultimately, learning the physical rules that govern molecules will allow models to extrapolate beyond the training data and to predict functions of more complex systems, especially for cases where biological insights are needed and the scale of the training data required for interpolation may be unattainable.

## Acknowledgements

We thank Shawn Costello, Jimin Yoon, and Patrick Almhjell for feedback on the manuscript. We thank Stephanie Wankowicz, James Fraser, Markus Braun, and Sarah Rauscher for helpful discussions. We thank anonymous reviewers for their helpful comments and suggestions. We are grateful for the support and suggestions from the Herschlag and Ma lab members. We would also like to thank the following individuals for their help generating AlphaFold3 predictions using the online AlphaFold server before the code was released: Patrick Almhjell, Krystal Brodsky, Abigail Chiang, Shawn Costello, Lauren Hagler, Albert Lee, Ben Ma, Allison Rabbani, John Shin, Garrett Wang, Yaming Yan, Chen Zhou, and Stella Zhu.

## Funding

National Key Research and Development Program of China (No. 2024YFA1307502) (JM)

The Science and Technology Innovation Plan of Shanghai Science and Technology Commission (No. 23JS1400200) (JM)

The Research Fund for International Senior Scientists (No. W2431060) (JM)

National Science Foundation grant MCB2322069 (DH)

Stanford Center for Molecular Analysis and Design Fellowship (SD)

## Author contributions

Conceptualization: SD, DH, NL, JM

Investigation: NL, SD

Visualization: NL, SD

Funding acquisition: DH, JM

Writing – original draft: SD, NL, DH

Writing – review & editing: SD, DH, NL, JM

## Data and materials availability

All data, code, and materials used in the analysis are available at Zenodo (doi: 10.5281/zenodo.19139881).

---

**Extended Data Fig. 1.**
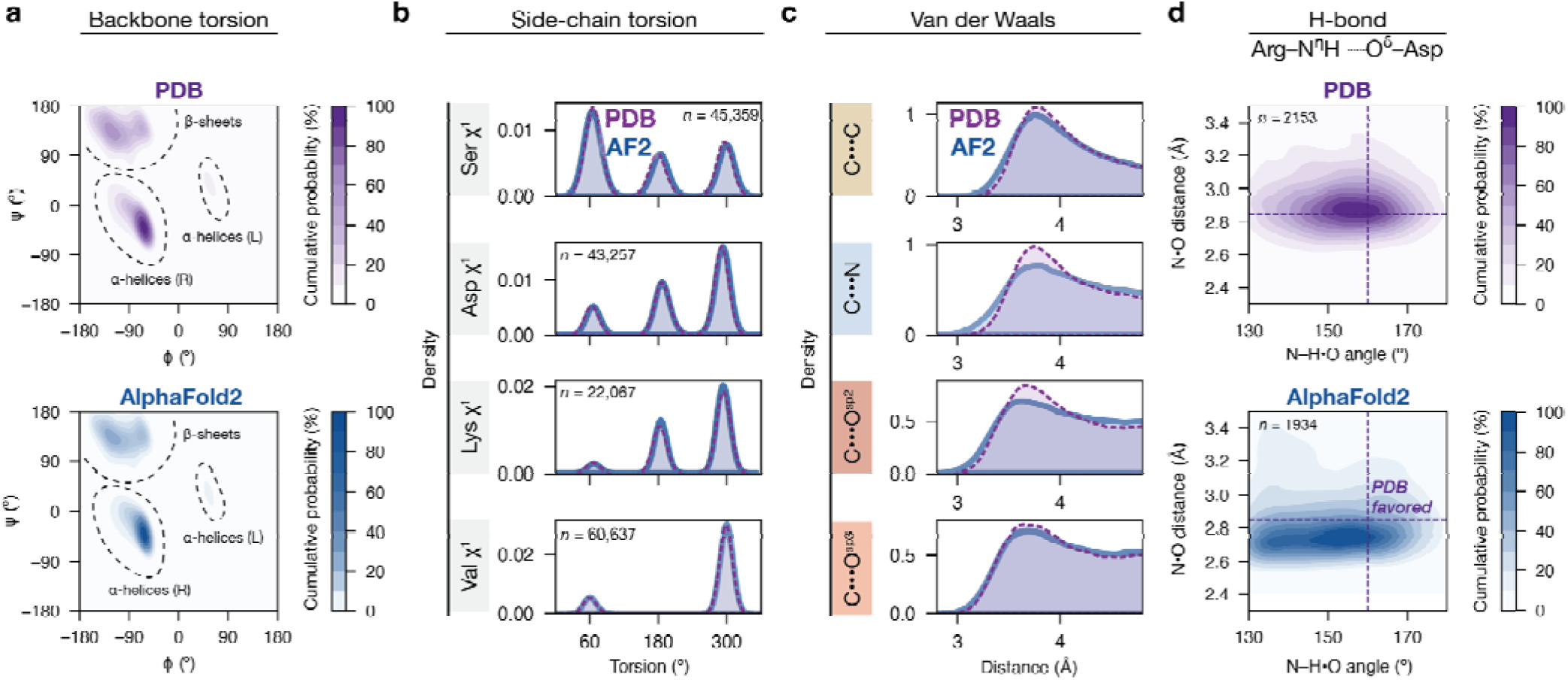
Biomolecular interaction energetics inferred by AlphaFold2. **(a)** Probability density distributions of backbone and angles (not including glycine and proline residues; see Supplementary Fig. 1). The color gradients indicate the probability mass under the contour in percentages, with darker shades indicating more probable conformational states. **(b)** Examples of probability density distributions of side-chain torsion angles observed in the PDB (purple) and those from AlphaFold2 predictions (blue); additional torsions are shown in Supplementary Fig. 3. **(c)** Probability density distributions of van der Waals distances for different interatomic contacts observed in the PDB (purple) and those from AlphaFold2 predictions (blue). Each distribution includes >75,000 van der Waals interactions (see Supplementary Table 3; additional interactions are shown in Supplementary Fig. 8). **(e)** Example probability density distributions for hydrogen bond angles (donor–H•acceptor) and distances (donor•acceptor) for arginine•aspartate side-chain hydrogen bonds. The color gradients indicate the probability mass under the contour in percentage, with the darker shades indicating more probable conformational states (10% per contour). This distribution includes Arg•Asp residue pairs that form a single hydrogen bond between their side-chains (excluding bidentate hydrogen bonds) to ensure an unambiguous comparison. Additional hydrogen bond distributions are shown in Supplementary Fig. 9.

**Extended Data Fig. 2.**
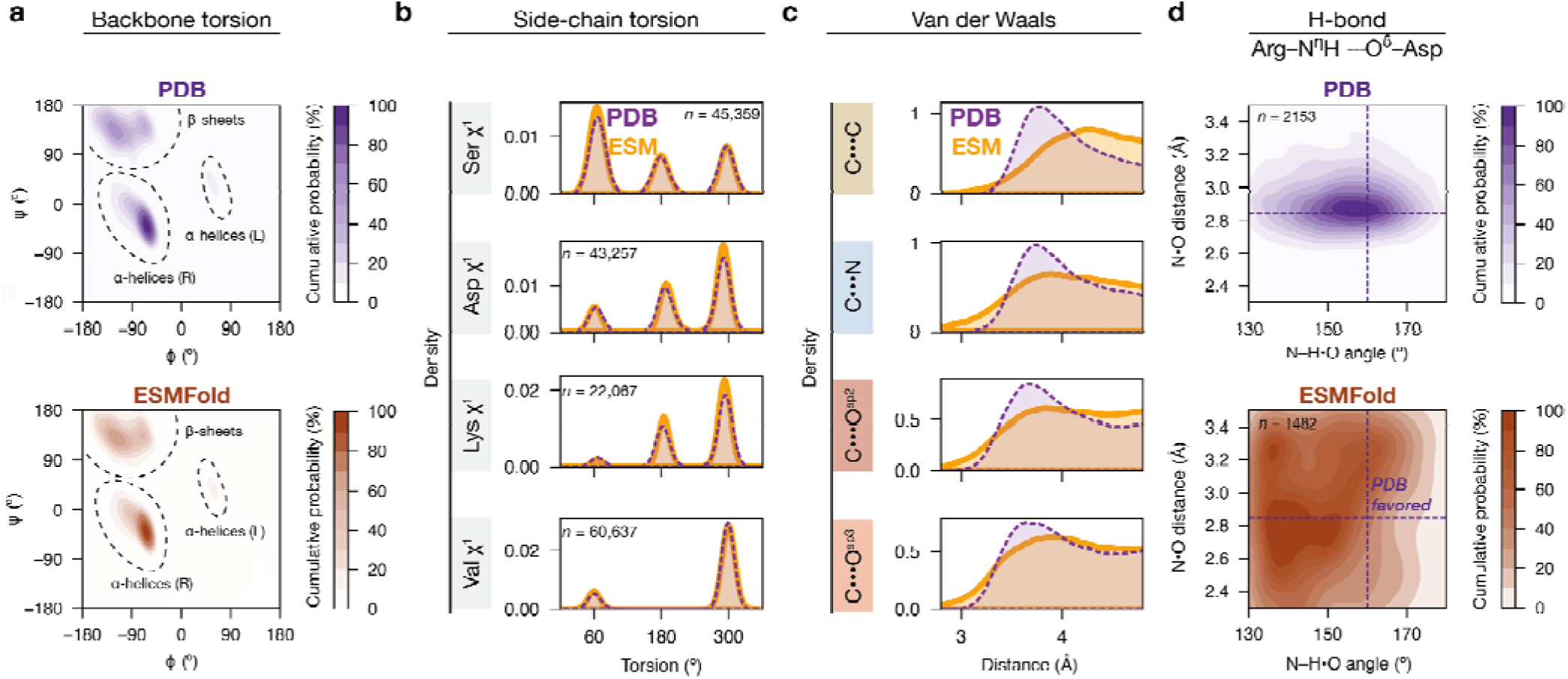
Conformational preferences for biomolecular interactions in ESMFold predictions. **(a)** Probability density distributions of backbone and angles (not including glycine and proline residues; see Supplementary Fig. 1). The color gradients indicate the probability mass under the contour in percentages, with darker shades indicating more probable conformational states. **(b)** Examples of probability density distributions of side-chain torsion angles observed in the PDB (purple) and those from ESMFold predictions (orange); additional torsions are shown in Supplementary Fig. 4. **(c)** Probability density distributions of van der Waals distances for different interatomic contacts observed in the PDB (purple) and those from ESMFold predictions (orange). Each distribution includes >75,000 van der Waals interactions (see Supplementary Table 3; additional interactions are shown in Supplementary Fig. 8). **(e)** Example probability density distributions for hydrogen bond angles (donor–H•acceptor) and distances (donor•acceptor) for arginine•aspartate side-chain hydrogen bonds. The color gradients indicate the probability mass under the contour in percentage, with the darker shades indicating more probable conformational states. This distribution includes Arg•Asp residue pairs that form a single hydrogen bond between their side-chains (excluding bidentate hydrogen bonds) to ensure an unambiguous comparison. Additional hydrogen bond distributions are shown in Supplementary Fig. 9.

**Extended Data Fig. 3.**
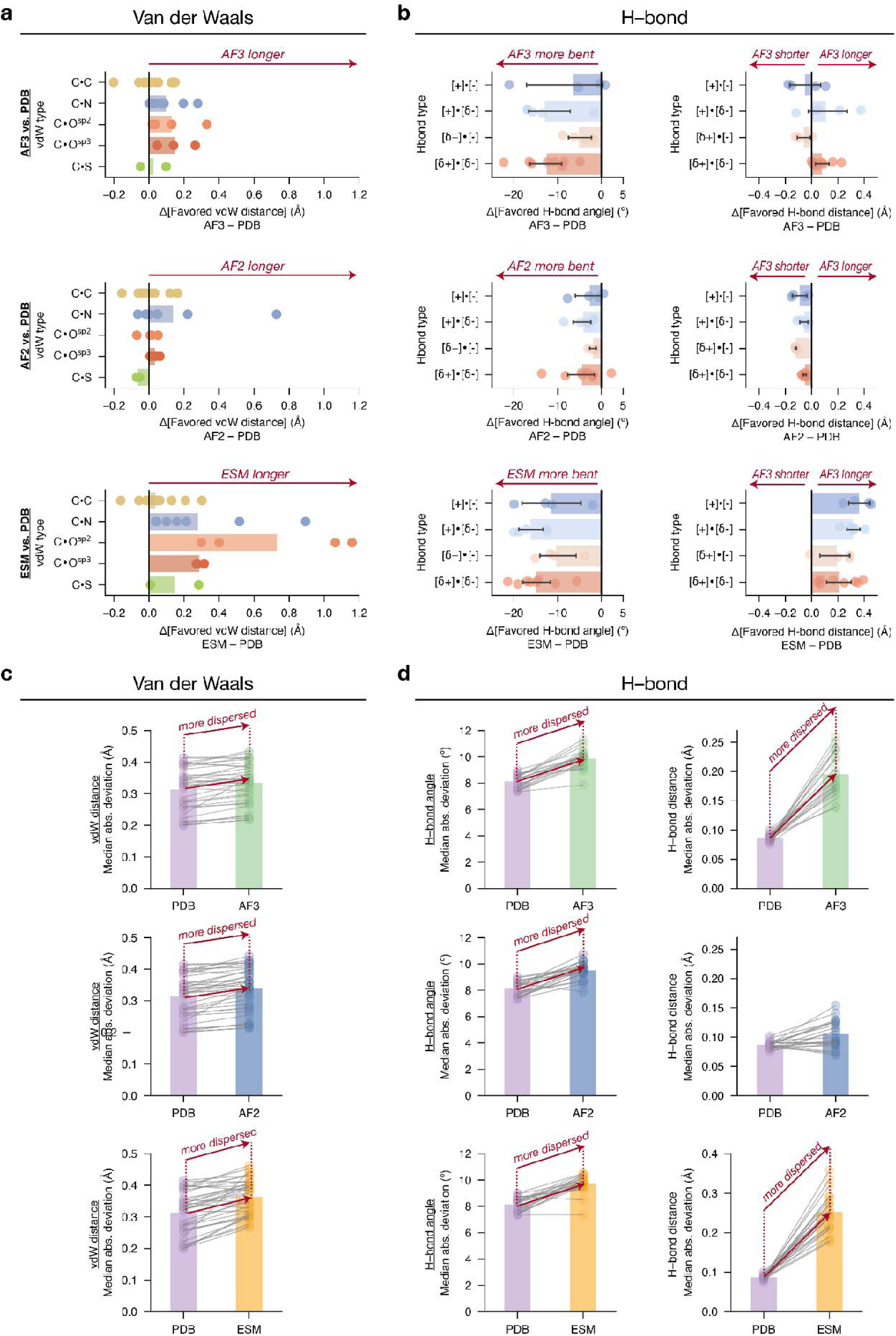
Differences in the conformational preferences for interactions in the PDB and in AlphaFold2, AlphaFold3 and ESMFold predictions. **(a)** Differences in the most favored van der Waals interaction distances in model predictions *versus* in the PDB distributions. Each circle represents the peak of the length distribution for a specific van der Waals type based on the amino acid and atom identities; data provided in Supplementary Table 6). **(b)** The same plot as (a) for hydrogen bond lengths and angles. Hydrogen bond types include charge•charge ([+]•[–]), charge•dipole ([*δ*+]•[–], [*δ*–]•[+]) and dipole•dipole ([δ+]•[δ–]); data provided in Supplementary Tables 7 and 8. **(C)** Differences in the spread of the distributions for van der Waals interaction distances in model predictions *versus* in the PDB distributions, quantified by the median absolute deviation (MAD). Each circle represents the MAD for a specific van der Waals interaction type based on the amino acid and atom identities; data provided in Supplementary Table 6. **(D)** The same plot as in (C) for hydrogen bond lengths and angles; data provided in Supplementary Tables 7 and 8.

**Extended Data Fig. 4.**
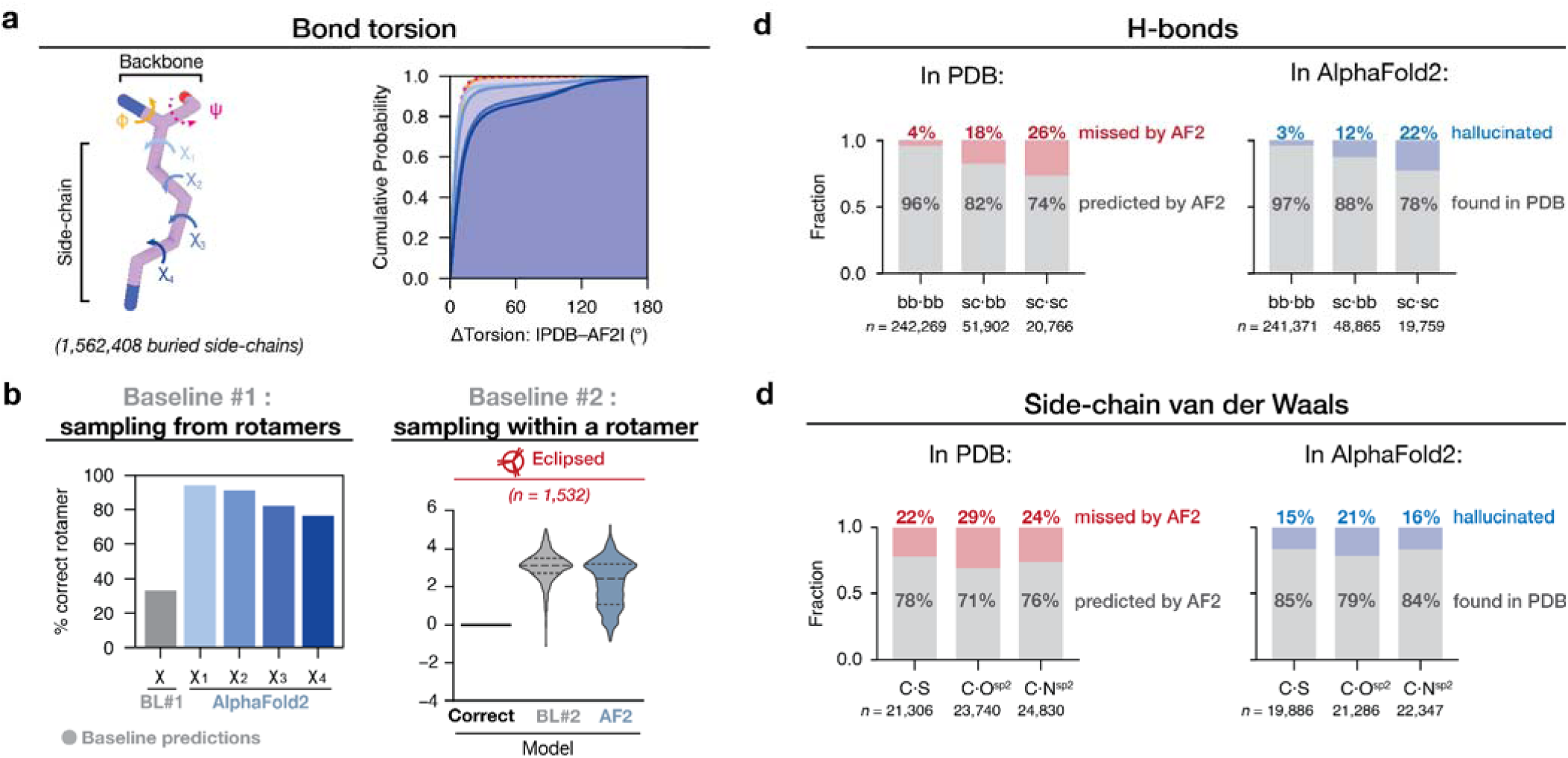
One-to-one comparisons for interactions in AlphaFold2 predictions *versus* in the corresponding PDB structures. **(a)** Left: definition of backbone and side-chain torsion angles, using a lysine residue as an example. Right: Cumulative distribution for side-chain torsion angle differences between AlphaFold2 predictions and the corresponding PDBs. **(b)** Left: percentage of correct side-chain rotameric states predicted by AlphaFold2 (light to dark blue) compared to a context-free baseline model (baseline #1) that randomly samples torsion angles according to the PDB distribution (gray). R ght: AlphaFold2 (blue) compared to the second baseline model (gray) for energy deviations from eclipsed PDB references. See main text Fig. 3c for descriptions of the baseline model. For reference, a theoretical model that predicts all bonds correctly would have Δ*E*_correct_ = 0, shown as the black lines. **(c)** Comparisons of buried backbone•backbone (bb•bb), backbone•side-chain (bb•sc), and side-chain•side- chain (sc•sc) hydrogen bonds in the PDB structures and in AlphaFold2 predictions. The gray bars indicate the numbers of hydrogen bonds in common in the PDB structure and the corresponding AlphaFold2 prediction (“ predicted by AF2”). For the PDB hydrogen bonds, the fraction that were not found in the AlphaFold2 predictions are shown as red bars (“missed by AF2”); for the AlphaFold2 hydrogen bonds, the fraction that were not found in the PDB structures are shown as blue bars (“hallucinated”). **(d)** The same comparisons as in panel c for three types of side-chain•side-chain van der Waals interactions.

**Extended Data Fig. 5.**
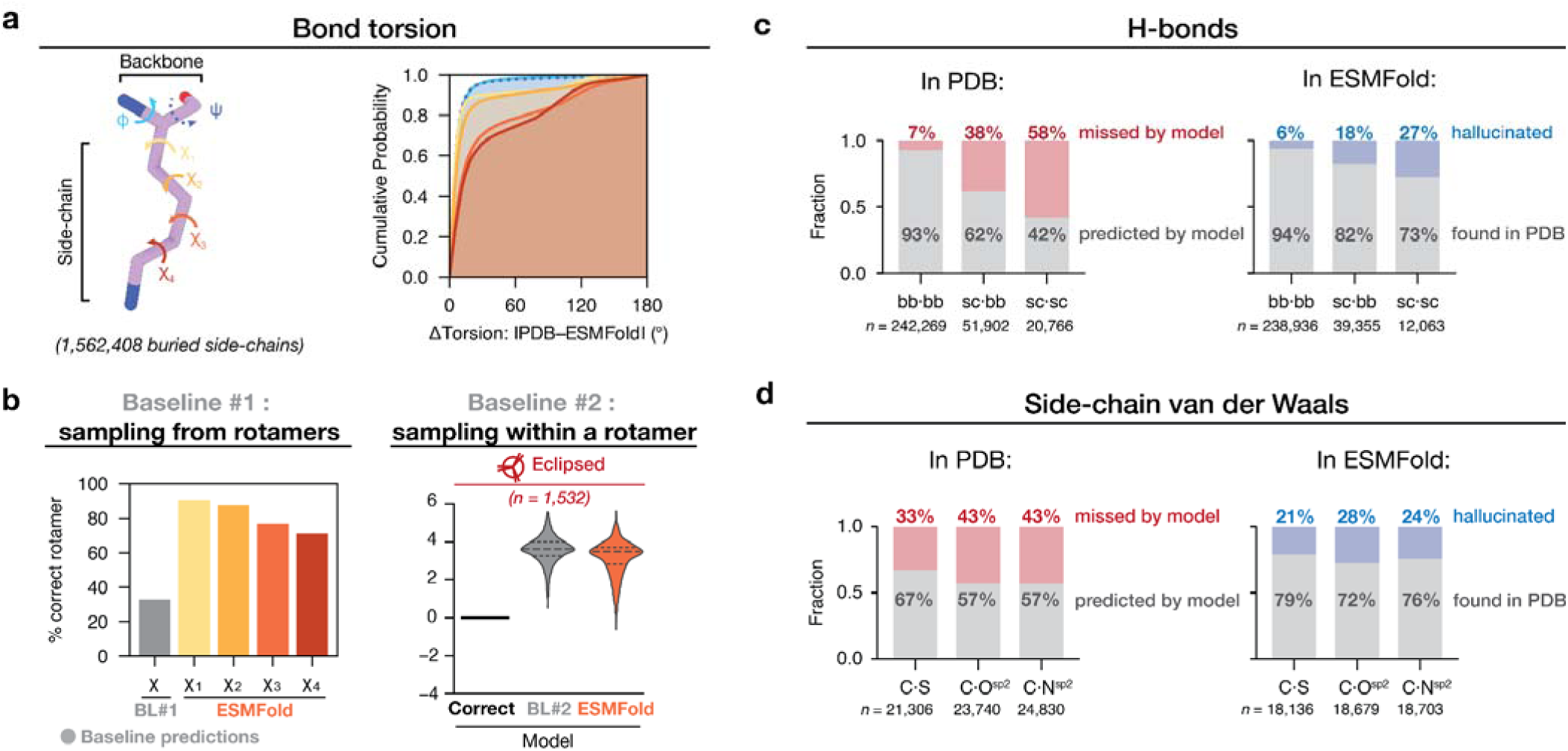
One-to-one comparisons for interactions in ESMFold predictions *versus* in the corresponding PDB structures. **(a)** Left: definition of backbone and side-chain torsion angles, using a lysine residue as an example. Right: Cumulative distribution for side-chain torsion angle differences between ESMFold predictions and the corresponding PDBs. **(b)** Left: percentage of correct side-chain rotameric states predicted by ESMFold (yellow red) compared to a first context-free baseline model that randomly samples torsion angles according to the PDB distribution (gray). Right: ESMFold (orange) compared to the second baseline model (gray) for energy deviations from eclipsed PDB references. See main text Fig. 3c for descriptions of the baseline model. For reference, a theoretical model that predicts all bonds correctly would have Δ*E*_correct_ = 0, shown as the black lines. **(c)** Comparisons of buried backbone•backbone (bb•bb), backbone•side-chain (bb•sc), and side-chain•side-chain (sc•sc) hydrogen bonds in the PDB structures and in ESMFold predictions. The gray bars indicate the numbers of hydrogen bonds in common in the PDB structure and the corresponding ESMFold prediction (“ predicted by model”). For the PDB hydrogen bonds, the fraction that were not found in the AlphaFold2 predictions are shown as red bars (“missed by model”); for the ESMFold hydrogen bonds, the fraction that were not found in the PDB structures are shown as blue bars (“hallucinated”). **(d)** The same comparisons as in panel d for three types of side-chain•side-chain van der Waals interactions.

**Extended Data Fig. 6.**
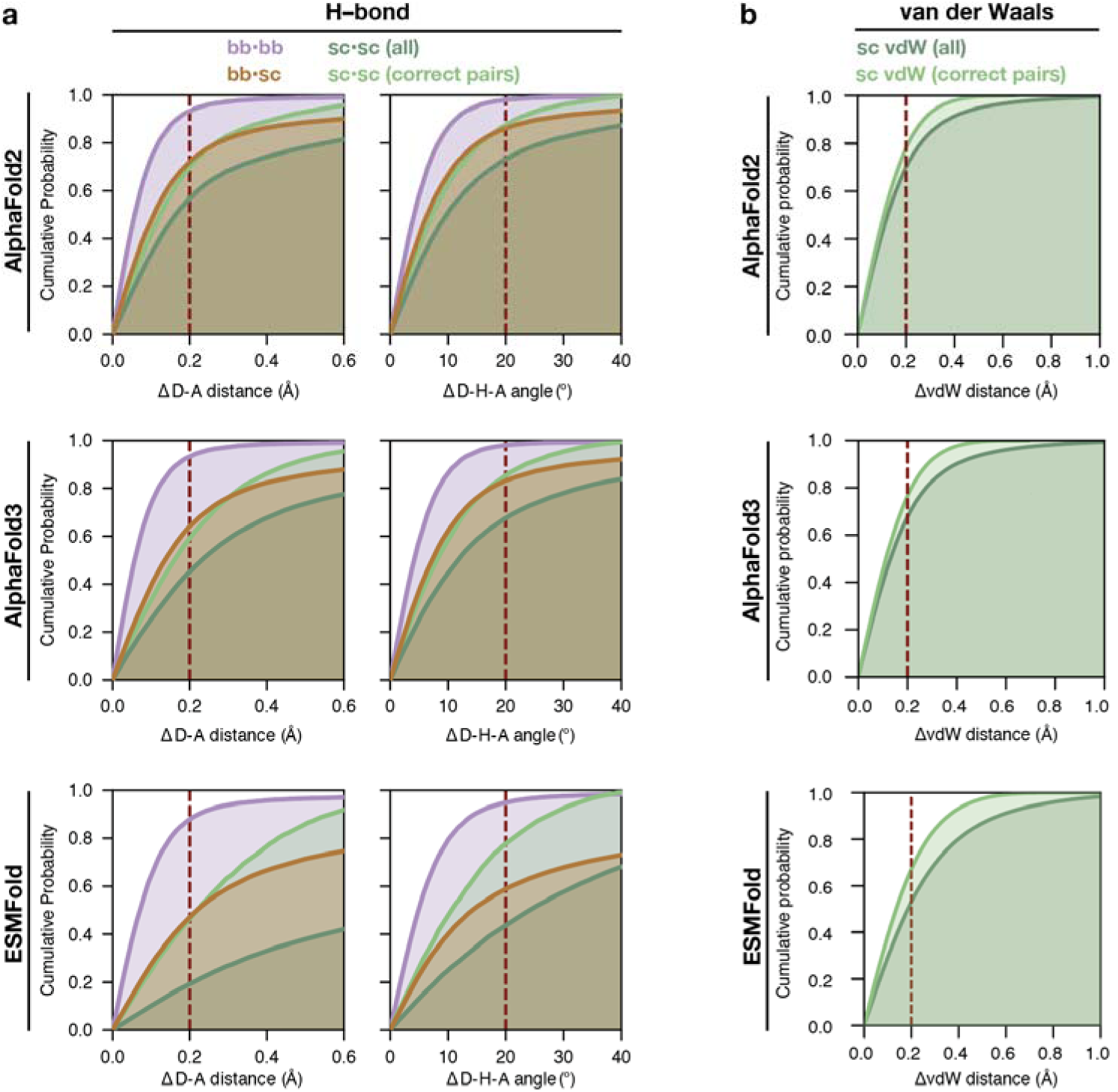
Cumulative distribution plots of the deviations of each non-covalent interaction geometry predicted by AlphaFold2, AlphaFold3 and ESMFold, compared to that in the corresponding PDB structures. For each interaction found in the PDB structures, the same atoms were identified in model predictions and the geometric parameters involving these atoms were calculated and compared. **(a)** Cumulative distribution plots for hydrogen bond distance and angles [“bb⋅bb”: backbone⋅backbone, *n* = 242,269; “bb⋅sc”: backbone⋅side-chain, *n* = 51,902 ; “sc⋅sc (all)”: all side-chain⋅side-chain, *n* = 20,766; “sc⋅sc (correct pairs)”: side-chain⋅side-chain groups predicted correctly by the model, as illustrated by the gray-shaded regions in Fig. 3d (*n* = 15,341 for AlphaFold2; 14,239 for AlphaFold3; 8,756 for ESMFold). The red line marks a geometry deviation threshold to visually guide comparisons (0.2 Å for donor⋅acceptor distance and 20° for donor–H⋅acceptor angle). **(b)** The same plots as in (a) for side-chain van der Waals interactions [“sc vdW (all)”: atom pairs forming van der Waals contacts in the PDB reference (n = 234,528); “sc vdW (correct pairs)”: van der Waals contacts that were correctly assigned by model predictions (n = 164,649 for AlphaFold2; 161,308 for AlphaFold3; 134,583 for ESMFold), as illustrated by the gray-shaded regions in Fig. 3e. The red line marks a contact distance deviation threshold (0.2 Å) to visually guide comparisons.

**Extended Data Fig. 7.**
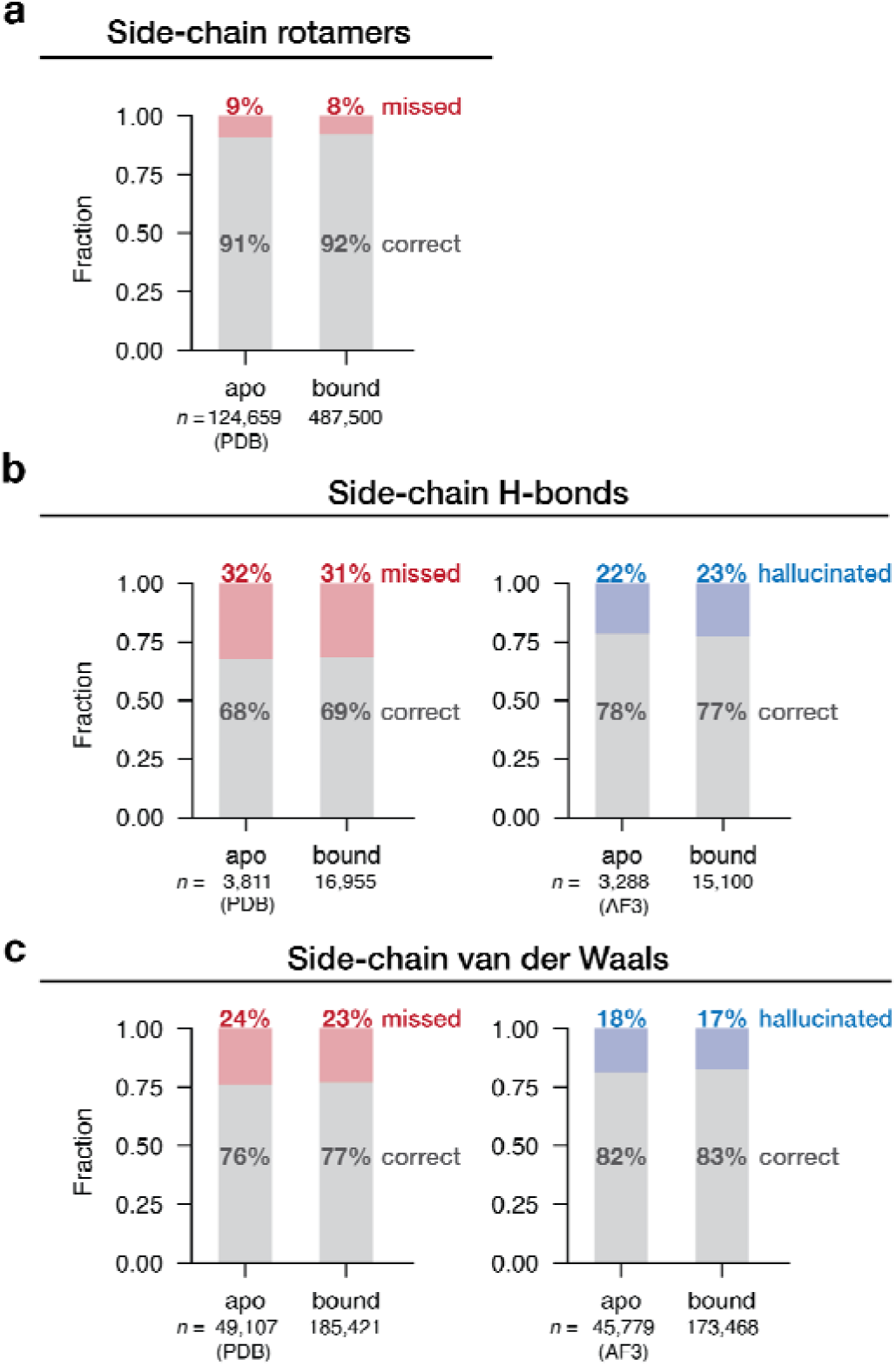
Comparison of the accuracy of AlphaFold3 predictions where the corresponding PDB structures are apo *vs.* bound to ligands or ions for **(a)** side-chain rotameric states, **(b)** side-chain•sidechain hydrogen bonds, and **(c)** sidechain•sidechain van der Waals interactions. For (a), the grey bars indicate the fraction of side-chain rotamers found in common between the PDB structures and the corresponding model predictions (“correct”). The fraction of interactions that were found in the PDB references but not found in model predictions is shown as red bars (“missed”). The annotated sample size is the total number of rotamers found in the PDB references (the same as that in the model predictions). For (b) and (c), the left subpanels are the same as in (a), showing the fraction of PDB interactions that are in common (“correct”) or missed by the models (“missed”). In the right subpanel, the fraction of interactions that were found in model predictions but not found in the PDB structure are shown as blue bars (“hallucinated”), and the total number of interactions found for each model is annotated below the bar.

**Extended Data Fig. 8.**
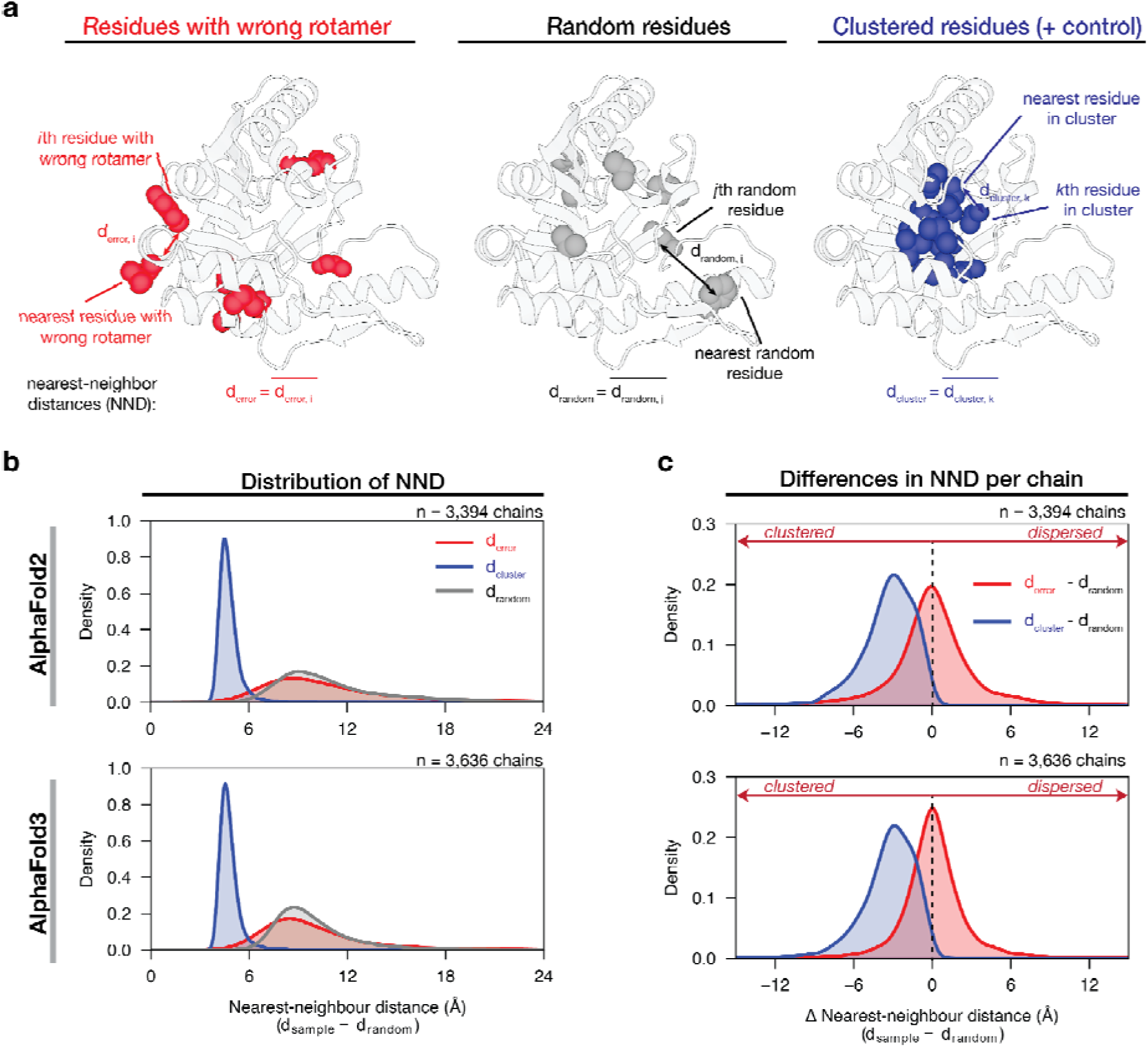
Assessing the extent of clustering of mis-assigned side-chain rotamers in AlphaFold2/3 predictions. The extent of clustering/dispersion of wrong rotamers was quantified by the mean nearest-neighbour distance (*Methods*). **(A)** Examples of nearest-neighbour distance (NND) calculation. For a protein chain (PDB: 1JAY_B), NND was calculated for all residues with mis-assigned rotameric states (d_error_, red); for a group of randomly selected rotamers with the same sample size (d_random_, gray); and as a positive control, for a cluster of residues located within a 7 Å sphere from the protein centroid (d_cluster_, blue). **(B)** Histograms of d_error_, d_random_ and d_cluster_ of all structures (*n* = 3,394 for AlphaFold2 and 3,636 for AlphaFold3; chains having < 2 residues with misassigned rotamers or having more residues with misassigned rotamers than residues without them are filtered out.) (*Method*). The NND distributions for residues with mis-assigned interactions in AlphaFold predictions resemble those of the randomly selected residues, rather than clustered residues. **(C)** The differences in NND, d_error_ – d_random_ and d_cluster_ – d_random_, were calculated for each protein chain and their distributions were plotted in red and blue, respectively. The NND differences between residues with prediction errors and randomly selected residues in each protein chain center around zero, indicating randomly distributed rather than clustered errors.

**Extended Data Fig. 9.**
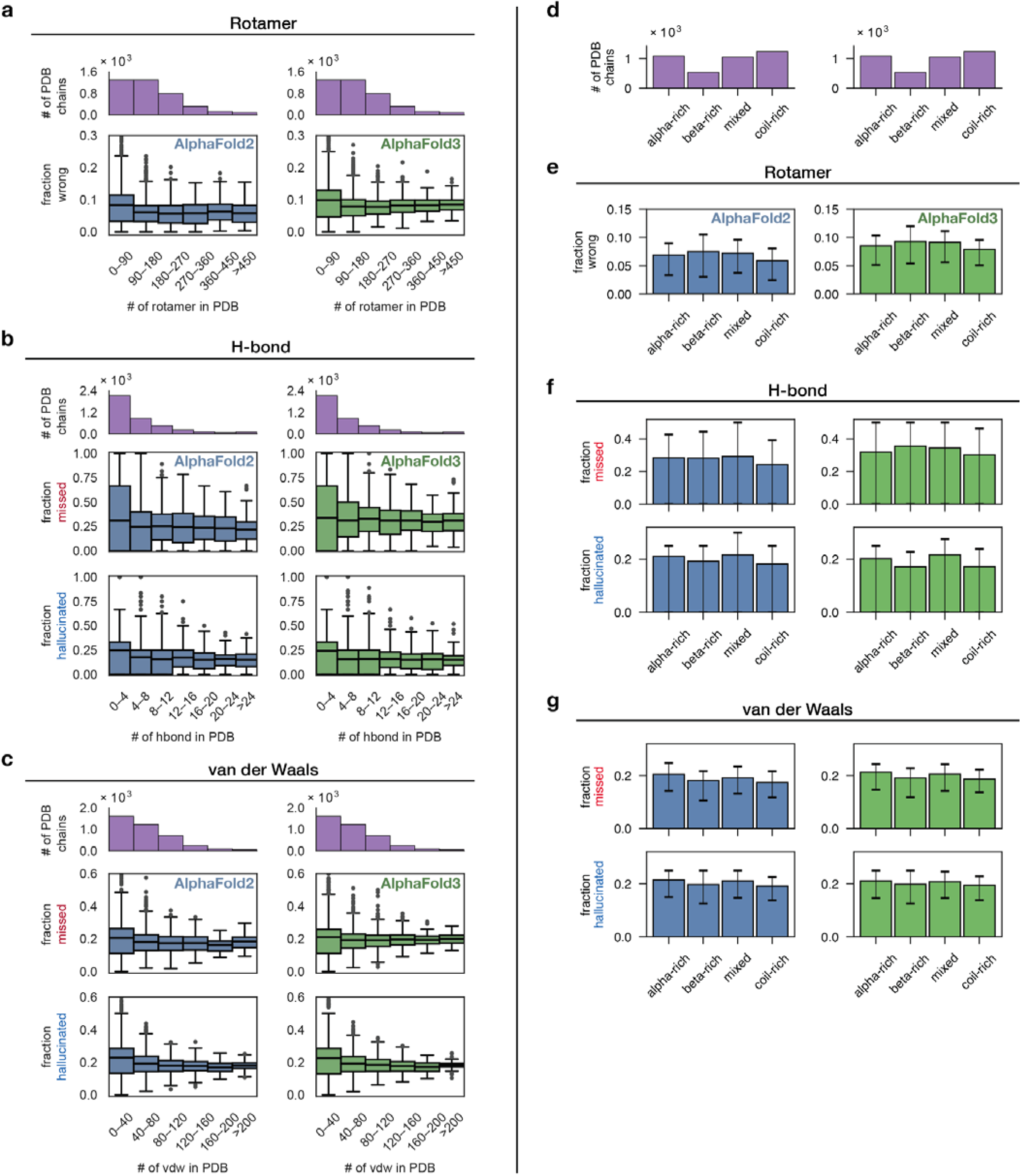
Error rates of AlphaFold2/3 side-chain interactions across proteins of different sizes (left) and across various structural classes (right). **(a)** Protein chains were binned by the number of native interactions (*x*-axis; top purple histograms). Boxplots give the mean ± interquartile range (IQR) (whiskers: 1.5 × IQR) of the per-chain error fraction in AlphaFold2 (blue) and AlphaFold3 (green) for side-chain rotameric states. **(b)** The same plots as in (a) for missed and hallucinated side-chain hydrogen bonds. **(c)** The same plots as in (a) and (b) for missed and hallucinated side-chain van der Waals contacts. **(d)** Number of *Top*2018 chains in each structural class. Structural classes were defined by the abundance residues forming different secondary structures in each protein chain (α-rich: α-helix >50% and β-sheet <25%; β-rich: α-helix <25%, β-sheet >50%; mixed: α-helix > 50%, β-sheet > 25%; coil-rich: all other structures). **(e)** Error fraction per-chain in AlphaFold2 (blue) and AlphaFold3 (green) for side-chain rotameric states in each structural class. Boxplots give the mean ± interquartile range (IQR) (whiskers: 1.5 × IQR) of the error fractions. **(f)** The same plots as in (e) for missed and hallucinated side-chain hydrogen bonds. **(g)** The same plots as in (e) and (f) for missed and hallucinated side-chain van der Waals contacts.

**Extended Data Fig. 10.**
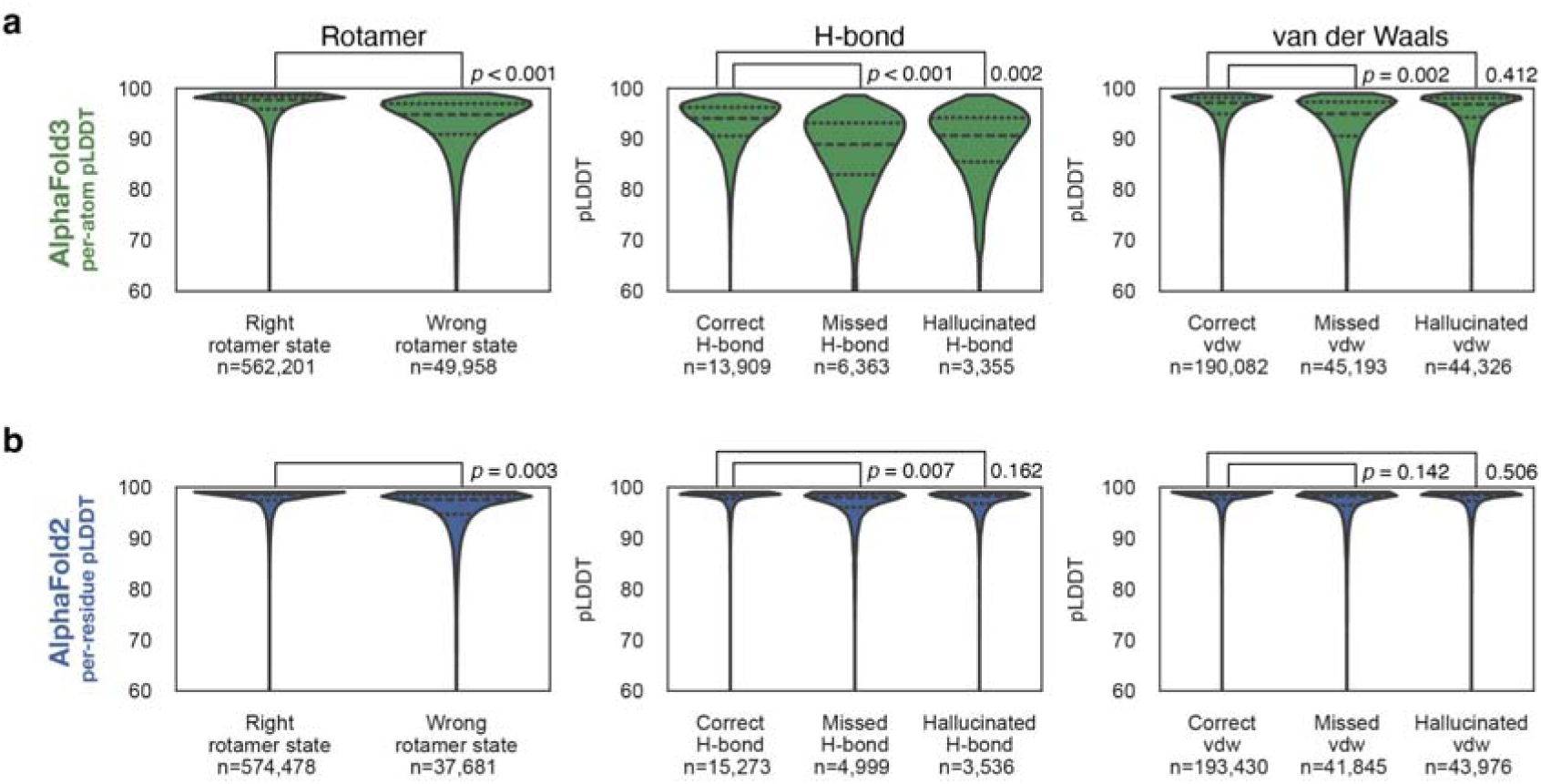
Correlation between predicted Local-Distance Difference Test (pLDDT) scores and the prediction accuracy of side-chain interactions. **(a)** AlphaFold3 per-atom pLDDT for correctly *versus* incorrectly predicted interactions. The pLDDT scores were averaged over the atoms involved in the molecular interactions (rotamer: the four atoms that define the torsion angle; hydrogen bond: the donor and acceptor heavy atoms; van der Waals: the two heavy atoms involved in the contact). **(b)** AlphaFold2 per-residue pLDDT for correctly *versus* incorrectly predicted interactions; for hydrogen bond and van der Waals interactions, the pLDDT scores were averaged over the residue pairs involved in the interactions. Statistical differences between the correct and incorrect distributions were assessed with the two-sample Kolmogorov–Smirnov (KS) tests. For each test, we report the mean *p*-value obtained from 20 independent iterations, where in each iteration we drew 100 observations at random from each distribution. The differences in the average pLDDT scores between the correct and incorrect predictions are significant but nevertheless small, and incorrect interactions involve residues with high average pLDDT scores (from 89 to 97), so that practically these scores would not allow users to identify incorrect or inaccurate interactions.

