## Supplementary Materials for "Physics-Grounded Evaluation to Guide Accurate Biomolecular Prediction"

**This file includes:**

Methods

Supplementary Figures 1 to 13

Supplementary Tables S1 to S10

References

### Methods

*Obtaining AlphaFold2, AlphaFold3 and ESMFold predictions for a subset of Top2018 structures*

The residue-level filtered *Top2018* dataset with 12,125 chains at <70% sequence homology was retrieved from ref^35^. AlphaFold2 predictions for proteins in the *Top*2018 dataset were obtained from the AlphaFold Database (https://alphafold.ebi.ac.uk/)^36^ by matching UniProt IDs. To ensure that identical sequences were used in our comparisons, the sequences of these AlphaFold2 structures were extracted using the **Bio.***PDB.PDBParser* module from the **biopython** package^66,67^ and compared with that of the corresponding experimental structures in the PDB. (The original structures deposited in the PDB were used in this step instead of the *Top*2018 structures whose sequences are incomplete due to the residue-level filtering performed in data curation.) We selected protein structures where the AlphaFold2 sequences match or cover the experimental sequences, resulting in 3939 chains. The AlphaFold2 predicted structures were then truncated to match exactly the residues in the *Top*2018 structures. AlphaFold3 predictions were generated using the AlphaFold3 server^9^ using sequences of the 3939 PDB chains (from 3848 original PDB structures). ESMFold predictions were generated using the esmfold_v1 model accessed from its official github page^68^.

The PDB files of the AlphaFold2, AlphaFold3 and ESMFold predicted structures were renumbered to match the corresponding *Top*2018 structures. For both AlphaFold2 and AlphaFold3, the predictions with the highest confidence among the five given predictions were used for the assessments, while each ESMFold run only yielded one prediction. The predicted Local-Distance Difference Test (pLDDT) scores for AlphaFold2 (per residue), ESMFold (per residue) and AlphaFold3 (per atom) were directly retrieved from the predicted PDB files.

The unrelaxed AlphaFold2 predictions were performed using the high-throughput computational modeling package, **EnzyHTP**^69,70^ (https://github.com/ChemBioHTP/EnzyHTP), using the sample script provided (run_alphafold.py). The EnzyHTP API performs the code and model parameters open-sourced by DeepMind^71,72^. Parameters were chosen to match the AlphaFold Monomer v2.0 pipeline used by AlphaFold DB^71^.

*Extracting molecular interaction geometries from experimental and predicted structures*

Covalent bonds: Covalent bond lengths between all heavy atoms and their angles in *Top*2018 and AlphaFold3 structures were measured by extracting atomic Cartesian coordinates with **Bio.***PDB.PDBParser^66,67^* and calculating their distances and angles with the **numpy** package^73^.

Bond torsions: Torsion angles (backbone 𝜓, 𝜑 and side-chain 𝜒^1^, 𝜒^2^, 𝜒^3^, 𝜒^4^ whenever applicable) of all experimental and predicted structures were calculated using the ***Ensemble*PDB*.****analyze.rotamer* module ^52^. All 𝜒 angles were normalized to the ranges defined in Shapovalov and Dunbrack^74^.

Hydrogen bonds: To calculate hydrogen bonds, hydrogen atoms were added to AlphaFold2 and AlphaFold3 predictions using the **Reduce** program^75^. To remove ambiguity in hydrogen bond identification that can result from inaccurate hydrogen atom positions, we only included donors with sp^2^ hybridized nitrogen because their hydrogen atoms can be unambiguously placed. Hydrogen bonds were then identified using the **MDAnalysis** package^76,77^ with the *analysis.hydrogenbonds.hbond_analysis* module, according to the following geometric criteria: i. donor–acceptor heavy atom distance (D•A distance) < 3.5Å; ii. donor–hydrogen–acceptor angle (D–H•A angle) > 130°. These criteria were used because various quantum-mechanical calculations have shown that hydrogen bonding interactions are significantly weakened beyond these thresholds^38–40^. Hydrogen bonds in AlphaFold2 and AlphaFold3 predicted structures that involve residues filtered in *Top2018* were removed.

Van der Waals interactions: Van der Waals contacts were identified between side-chain carbon atoms of aliphatic amino acids (Ala, Val, Pro, Leu, Ile) and side-chain non-sp^3^-carbon heavy atoms (sp^2^ carbon; sp^2^ oxygen; sp^3^ oxygen; sp^2^ nitrogen; sulfur) with a detection threshold of 5 Å. We applied the line-of-sight method^78,79^ to eliminate van der Waals interactions shielded by an additional atom or interaction. If multiple van der Waals contacts of the same type were found between a residue pair, then the contact of the shortest distance was selected.

*Comparing conformational distributions of molecular interactions*

The peaks of the conformational distributions were determined by fitting a probability density function via kernel density estimation (**scipy**.*stats.gaussian_kde*)^80^ and calculating the highest probability position. The extent of dispersion of the distributions were evaluated by calculating the mean absolute deviation (**scipy***.stats.median_abs_deviation*)^80^.

*Calculating energy functions from conformational distributions*

To quantitatively determine the agreement between the conformational distributions derived from model predictions and the *Top*2018 structures, we calculated knowledge-based energy functions. For each geometric parameter of each interaction type, we pooled all occurrences found across the 3939 chains (from *Top2018* or from model predictions) and transformed the distribution into an energy function E_PDB_(ꭓ) via the Boltzmann relationship with the **EnsemblePDB**.*analyze.energy* module^52^ which operates as described below.

The distributions were cut into a specified number of bins (*n*_bins_) and the number of samples within each bin (*n*) was calculated. The number of bins were set depending on the sample size of each distribution such that the bin widths are between 0.08 to 0.1 Å for distances and between 2 to 4° for angles and dihedrals and the resulting energy functions were not overly rugged. The probability of observing a sample in a certain bin, *P*_i_, was calculated as *n* divided by the total sample size (*n*_total_). The reference state (*E*_ref_) was defined as a random distribution where the probability of observing any state is *P*_ref_ = 1/*n*_bins_ such that *E*_ref_ = 0. The energy of each bin relative to *E*_ref_ (*E*) was calculated via the Boltzmann relationship, *E* = –k_B_*T*•ln(*P*_i_/*P*_ref_) where k_B_ is the Boltzmann constant and *T* = 298 *K*. The energy functions were saved as tables (.csv files) specifying each bin and its *E* value. The Pearson correlation coefficients *R* and *p*-values for the comparisons of model predictions *versus* *Top*2018 energy functions were calculated using **scipy**.*stats.pearsonr^80^*.

To assign a knowledge-based energy *E* for a certain geometric measurement (*x*) using the energy function, the bin containing that value *x* was found and the corresponding *E*(*x*) value was queried; if the geometric value was outside the range of geometries covered by the energy function (because the conformer was too rare to be present in the original crystallographic distributions), then the maximum *E* value was assigned.

*One-to-one comparison of individual interactions*

We selected buried residues for our comparisons based on the relative solvent accessible surface area (relASA). The relASA value and secondary structure of each residue were calculated using the Define Secondary Structure of Proteins (**DSSP**) algorithm^81,82^ implemented via the **Bio.***PDB.DSSP* module from the **biopython** package^66,67^. We used original PDB structures such that the residues filtered in *Top*2018 were included for relASA calculations. Buried residues were defined as those with relASA < 0.25. For hydrogen bonds, we also excluded those involving donors or acceptors that underwent Asn/Gln/His flipping during *Top*2018’s data curation process.

Because we renumbered the predicted PDB files such that each residue match those from the *Top*2018 file, we simply compared the covalent bonds and bond torsions for each residue of the same numbers in each protein chain.

For hydrogen bonds and van der Waals interactions, we first determined whether the same residue-atom pairs were found to form interactions in the AlphaFold predictions and in the *Top*2018 structures. For hydrogen bonds, the criteria were the same as described above (*Extracting molecular interaction geometries from experimental and predicted structures*). For van der Waals interactions, we additionally require the side-chain heavy atom contact distance to be within 0.4 Å of the ideal distance, corresponding to the peak areas of the distributions shown in Fig. 2d (3.4 to 4.2 Å for the contact pair of sp^3^C–sp^2^O and sp^2^C–sp^2^N; 3.3 to 4.1 Å for sp^3^C–sp^3^O; and 3.6 to 4.4 Å for sp^3^C–S). When ambiguous atoms (e.g., Asp/Glu carboxylate oxygens) are encountered, we ranked these non-covalent interactions by heavy-atom pair distance and compared them in order—the shortest PDB interaction with the shortest model interaction, the next shortest with the next shortest, and so on. The hydrogen bonds found in *Top*2018 but not in the model prediction were assigned as “missed” and those found in predictions but not in *Top*2018 were assigned as “hallucinated”.

To directly compare interaction geometries, we calculated the geometric parameters (using the **biopython** and **numpy** packages) for each interaction found in the *Top*2018 structures. We then located the same residue-atom pairs in the corresponding AlphaFold predictions (which may or may not form interactions) and calculated the same geometric parameters.

*Assessing the extent of clustering of prediction errors via nearest-neighbor distances*

To determine whether prediction errors tend to cluster, we compared nearest-neighbor distances of residues with incorrect predictions to (1) those of randomly sampled residues and (2) those of clustered residues in a 7 Å sphere centered at the protein centroids (Extended Data Fig. 8). In each protein chain, residues with at least one side-chain assigned an incorrect rotameric state were identified and the number of errors were calculated ($n_{error})$_._ For each residue *i* containing incorrect rotamers, we measured the distance of its C_ɑ_ to that of the nearest residue that also contain incorrect rotamers ($d_{error,i}$). We then calculated the mean of the nearest neighbor C_ɑ_-C_ɑ_ distances of each protein as $d_{error} =\frac{1}{n_{error}} \sum_{i}^{n_{error}} d_{error,i}$, a metric to quantify the extent of clustering between residues with incorrect side-chain torsions in a protein chain.

To compare $d_{error}$ to the same metric calculated for randomly selected residues, we sampled the same number of side-chain bonds ($n_{error})$ from all side-chains of the same protein and calculated the mean nearest neighbor C_ɑ_-C_ɑ_ distance as described above ${(d}_{random})$. We performed random sampling for 200 times for each structure and calculate their mean as $\underline{d_{random}}$.

To compare $d_{error}$ to the same metric calculated for neighboring residues (i.e., an upper bound for clustering), we compared $d_{error}$ to the mean nearest neighbor distance of a set of residues where at least one atom is in the 7 Å sphere centered at the protein centroids ${(d}_{cluster})$. Comparisons of $d_{error}$ to $\underline{d_{random}}$ and $d_{cluster}$ provided a quantitative measure for the extent of dispersion *versus* clustering for residues predicted to be in incorrect rotameric states (Extended Data Fig. 8).

*Evaluating AlphaFold’s ability to generate conformational ensembles*

We curated 25 multi-conformer structural models refined from X-ray crystallographic datasets determined at >200 *K* and these multi-temperature (MT) ensemble models represent nine different proteins in their apo states (Supplementary Table 10). For each protein, we performed predictions on the AlphaFold3 server using random seeds for 70 times. We used the ***Ensemble*PDB*.****analyze.rotamer* module^52^ to calculate all the bond torsions, including all modeled alternative conformers.

To determine the number of states in the MT datasets for each bond of each protein, we aggregated all the torsional angles modeled for each bond into a histogram and calculated the peaks of the histogram using **scipy**.*signal.find_peaks^80^*, taking peaks that were at least 30º apart. We performed the same calculations to find the states predicted by AlphaFold3 and compared the number of states and the peak positions to those found in the MT datasets.

*Assessing the effect of ligand or ion bound structures in Top2018 on evaluation results*

We divided the 3,939 *Top*2018 structures into two subsets: an “apo” set, defined as structures in which the only non-protein molecules were water, solvents, buffering agents, cryoprotectants, and reducing or stabilizing additives, and a “bound” set, comprising structures that contain ions or ligands. The PDB codes for the non-protein molecules allowed in the apo set are: HOH, WAT, DOD, EDO, GOL, PEG, PE4, PE5, PE8, PGE, MPD, HEZ, HPS, CIT, MES, MOP, MOPS, ADA, TRIS, BIS, ACD, FMT, PCA, BME, DTT, SOH, EDT, ACT, TRS, B3P, or CYO; all structures containing additional non-protein molecules or atoms were assigned to the bound set.

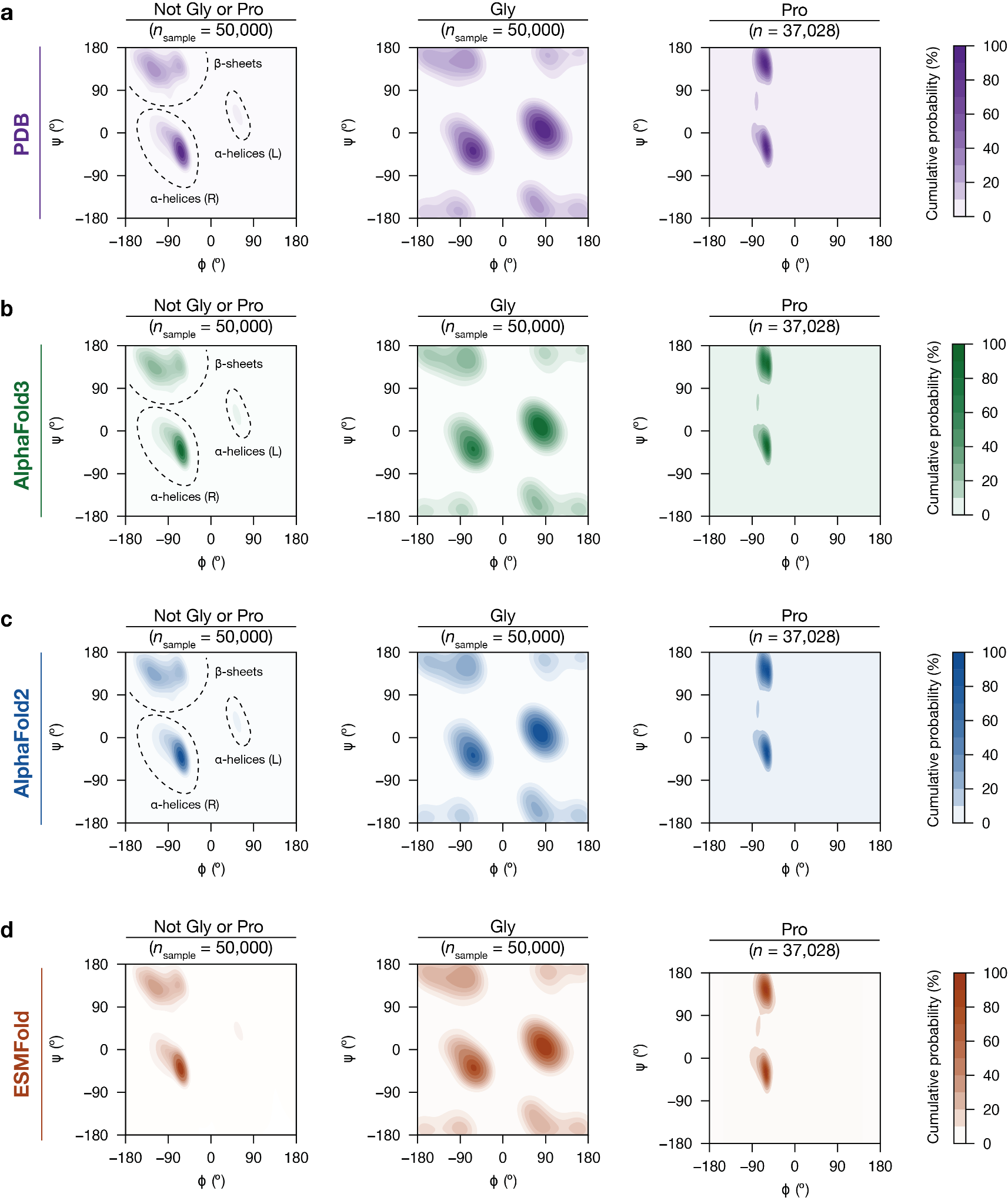

### Supplementary Fig. 1. Probability density distributions of backbone torsion angles (𝜓, 𝜙) in (a) the *Top2018* PDB structures (purple), (b) AlphaFold3 predictions, (c) AlphaFold2 predictions and (d) ESMFold predictions. The distributions were shown aggregated for all residues except glycines and prolines (left); glycines (middle); and prolines (right). The color scale indicates cumulative probability of data under the contour in percentage, where the darker shades indicate more probable conformational states. The plots show 50,000 randomly sampled 𝜓, 𝜙 angles except for proline residues where only 37,028 residues were found across the 3939 chains.

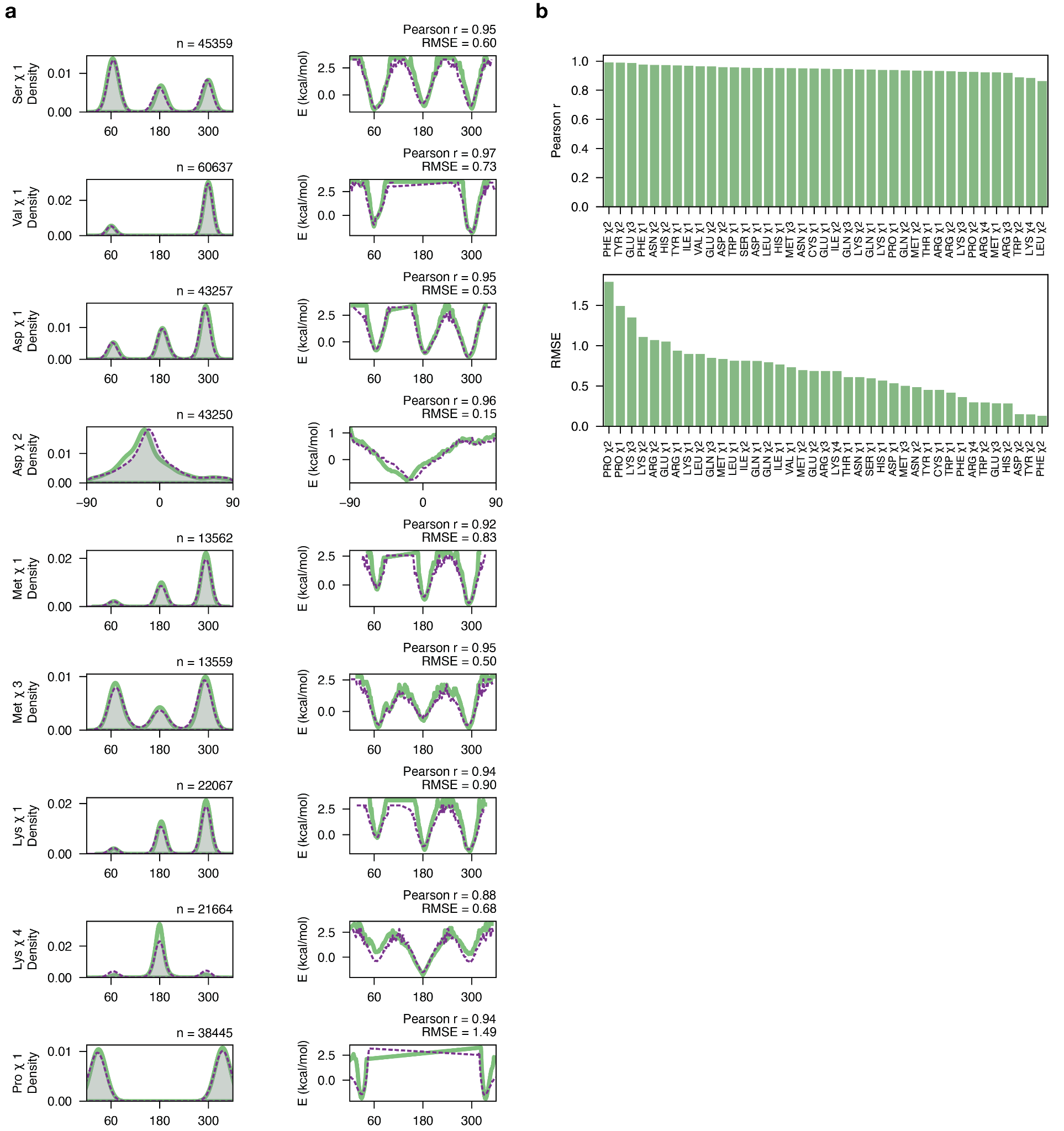

### Supplementary Fig. 2. The agreement between the PDB and AlphaFold3 predictions for side-chain bond torsion distributions. (a) Probability density distributions (left) from the 3939 AlphaFold3 predictions (green) versus the *Top*2018 PDB structures (purple). To quantitatively assess their agreement, we converted the distributions into energy functions using the Boltzmann relationship (see *Methods*) and calculated the Pearson correlation and RMSE values between the AlphaFold3 and PDB energy functions. These examples were randomly chosen, with all Pearson *r* and RMSE values (in kcal/mol) shown in panel (b). P-values are <10^–18^ for all correlations.

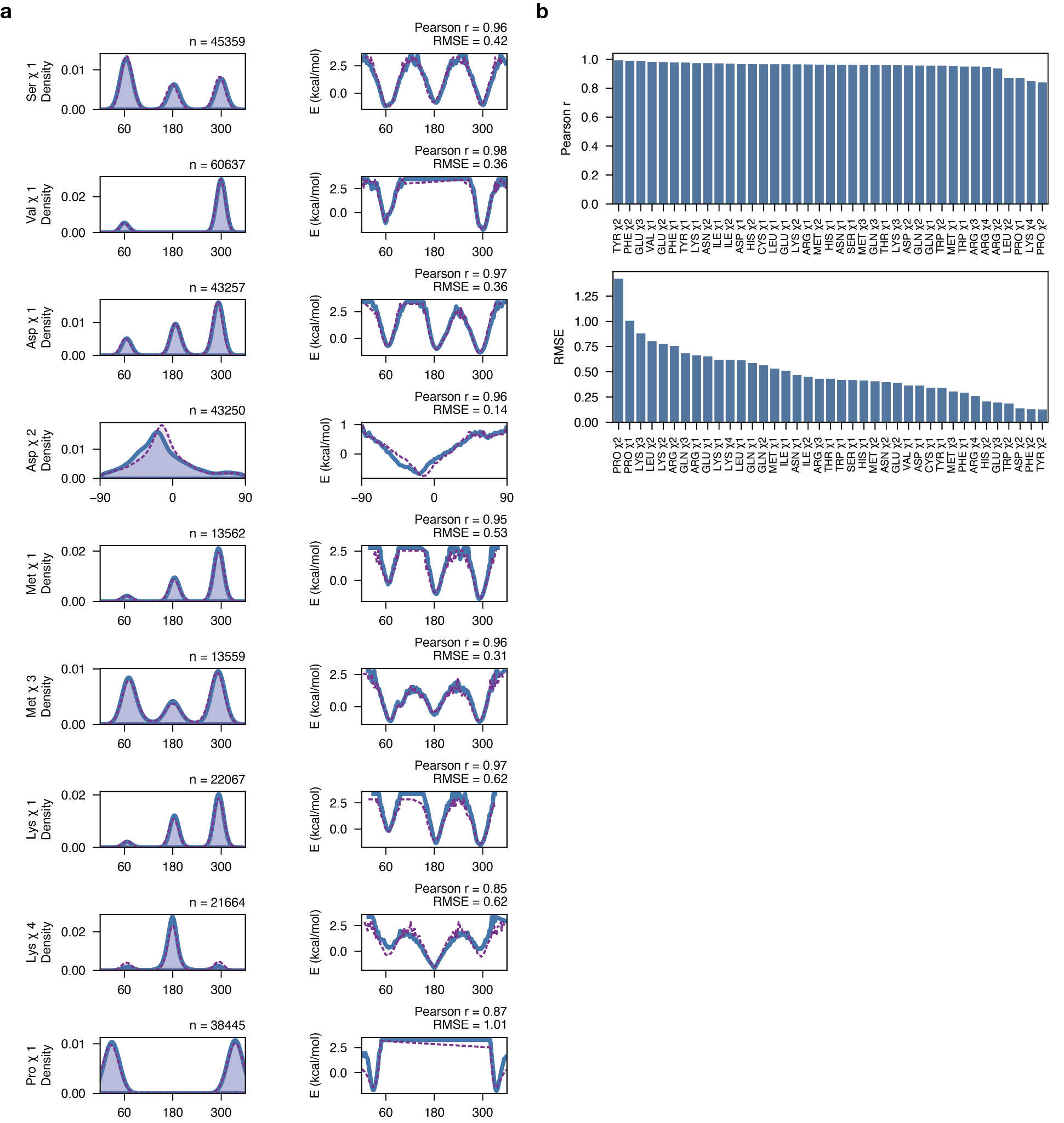

### Supplementary Fig. 3. The agreement between the PDB and AlphaFold2 predictions for side-chain bond torsion distributions. (a) Probability density distributions (left) from the 3939 AlphaFold2 predictions (blue) versus the *Top*2018 PDB structures (purple). To quantitatively assess their agreement, we converted the distributions into energy functions using the Boltzmann relationship (see Methods) and calculated the Pearson correlation and RMSE values between the AlphaFold2 and PDB energy functions. These examples were randomly chosen, with all Pearson *r* and RMSE values (in kcal/mol) shown in panel (b). *P*-values are <10^–14^ for all correlations.

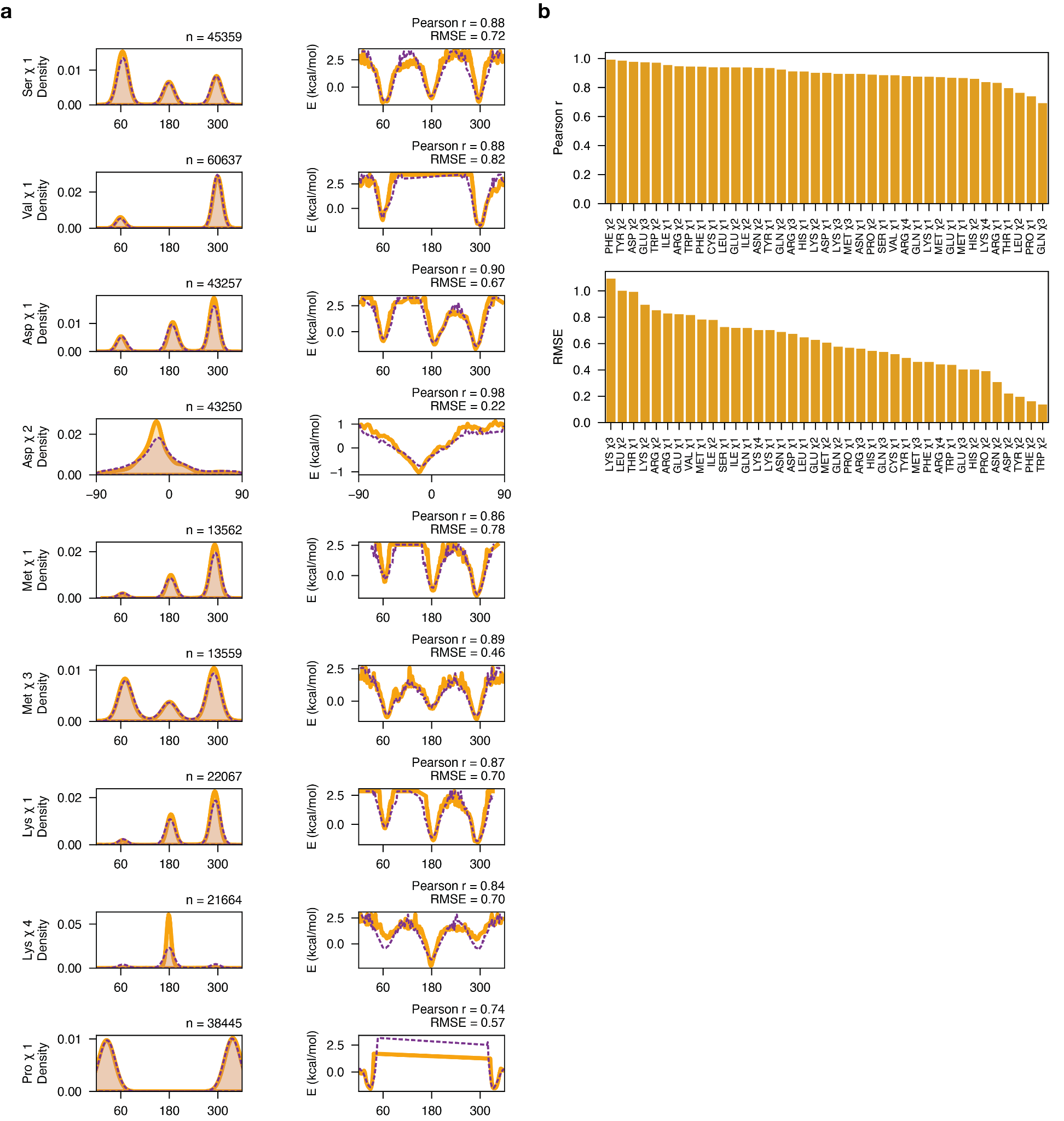

### Supplementary Fig. 4. The agreement between the PDB and ESMFold predictions for side-chain bond torsion distributions. (a) Probability density distributions (left) from the 3939 ESMFoldpredictions (orange) versus the Top2018 PDB structures (purple). To quantitatively assess their agreement, we converted the distributions into energy functions using the Boltzmann relationship (see Methods) and calculated the Pearson correlation and RMSE values between the ESMFold and PDB energy functions. These examples were randomly chosen, with all Pearson *r* and RMSE values (in kcal/mol) shown in panel (b). P-values are <10^–7^ for all correlations.

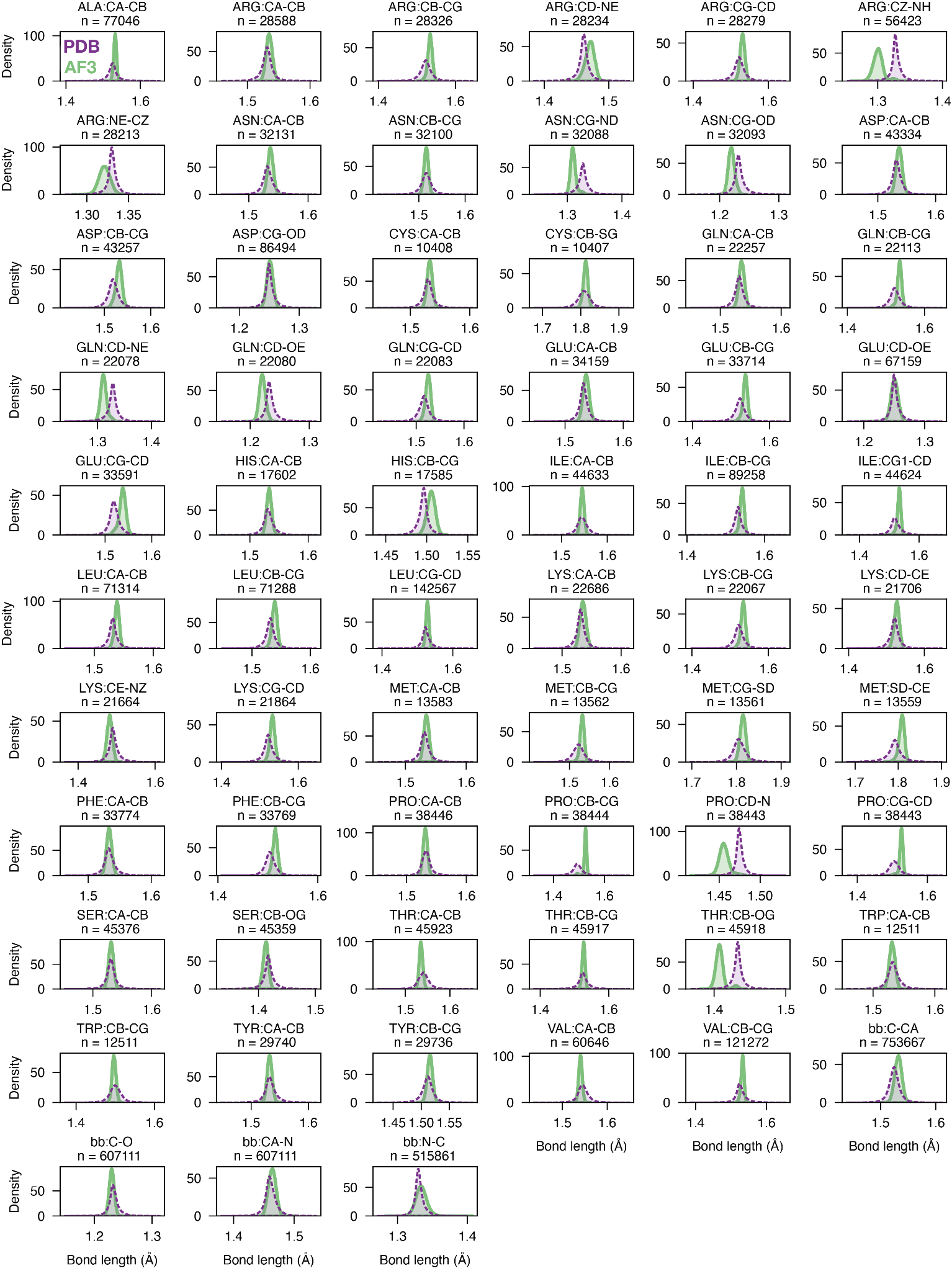

### Supplementary Fig. 5. Probability density distributions of covalent bond lengths from AlphaFold3 predictions (green) and *Top2018* PDB structures (purple). The amino acid and the atoms forming the bond are labeled on each subplot as well as the sample size. The *x*-axes use different scales for different covalent bond types to better visualize the individual distributions. Systematic deviations in the peak positions and the dispersion of the distributions are prevalent, as shown in the comparisons here and quantified in Supplementary Table 4.

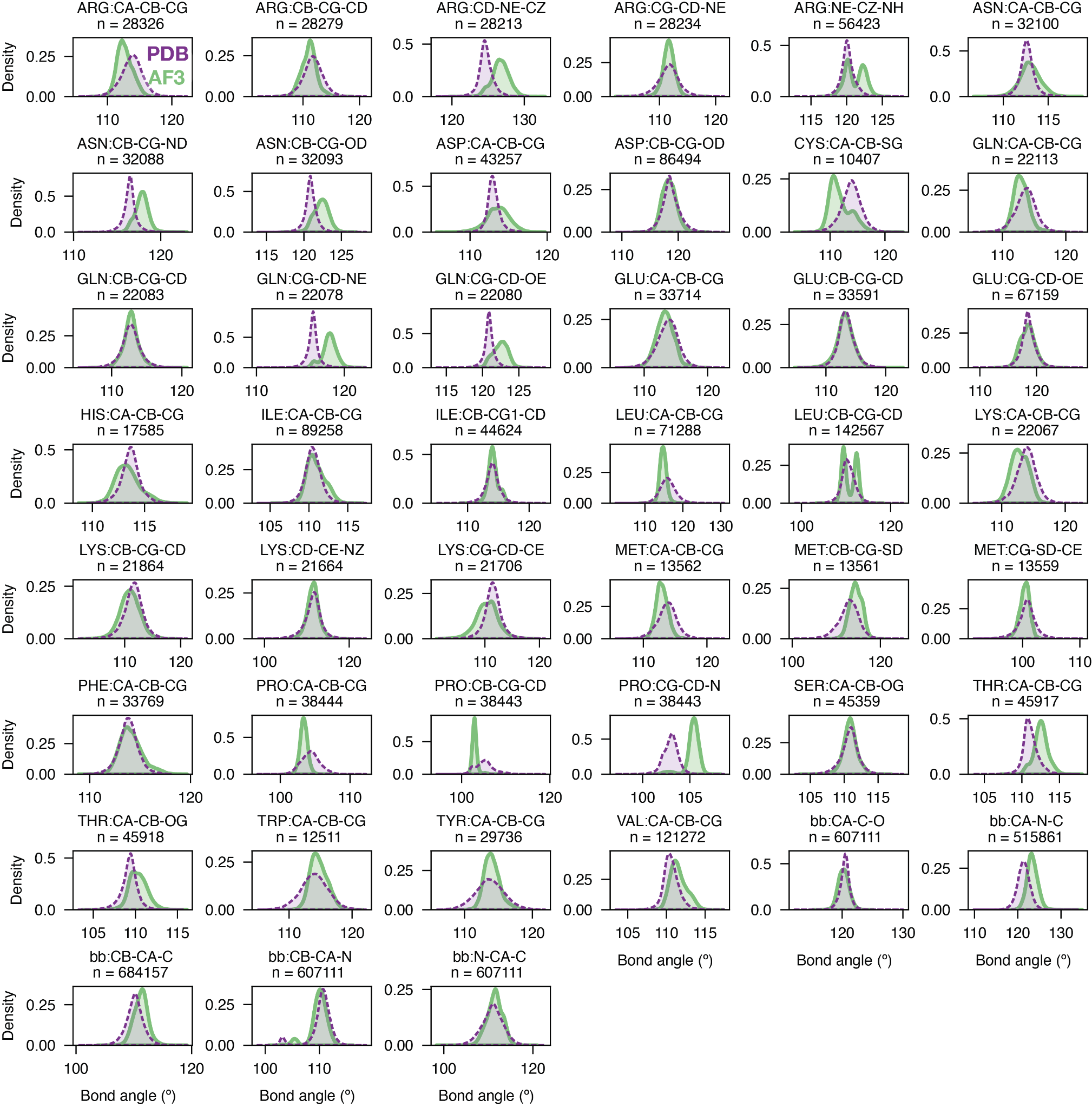

### Supplementary Fig. 6. Probability density distributions of covalent bond angles from AlphaFold3 predictions (green) and the corresponding *Top2018* PDB structures (purple). The amino acid and the atoms defining the bond angle are labeled on each subplot as well as the sample size. The *x*-axes use different scales for different covalent bond types to visualize the individual distributions. Systematic deviations in the peak positions and the dispersion of the distributions are prevalent, as shown in the comparisons here and quantified in Supplementary Table 5.

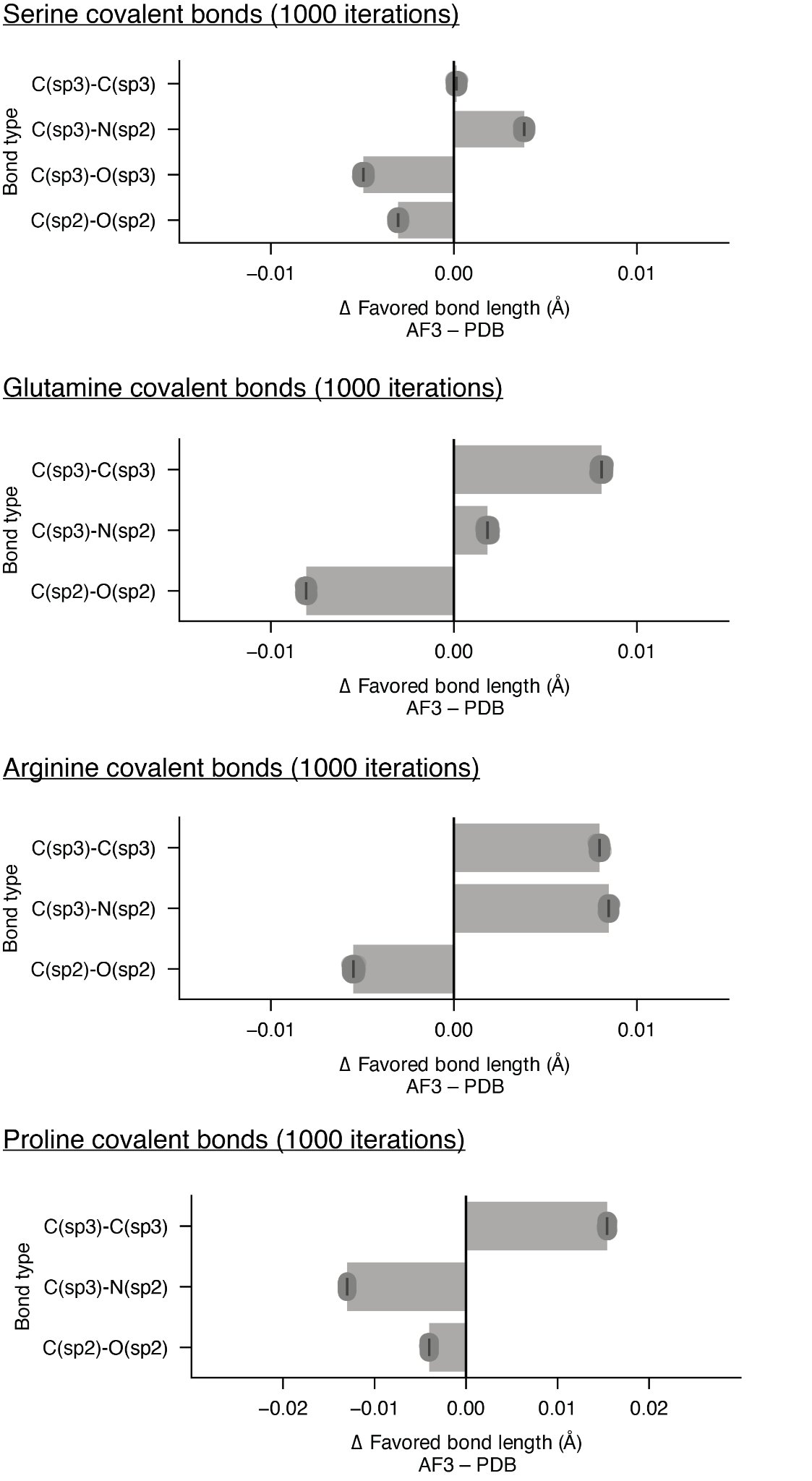

### Supplementary Fig. 7. Example bootstrap analyses of the AlphaFold3 *vs.* PDB covalent bond lengths in serine, glutamine, arginine and proline residues. For each covalent bond type, we randomly resampled 20,000 covalent bonds from the original AlphaFold3 and PDB distributions of that type and calculated the difference in the mean values (AlphaFold3 – PDB). This procedure was then repeated for 1000 iterations, giving 1000 difference values. The 1000 data points are plotted as grey circles (but only one or a few circles are visible on the plots because the results are highly consistent and thus give overlapping circles). The bars indicate the mean of the 1000 data points and the error bars indicate their 95% confidence interval (CI). The CI is extremely narrow (at the scale of 10^-4^ Å and thus appear as single lines) compared to the size of the differences; thus the bond length distortions observed in AlphaFold3 are highly robust.

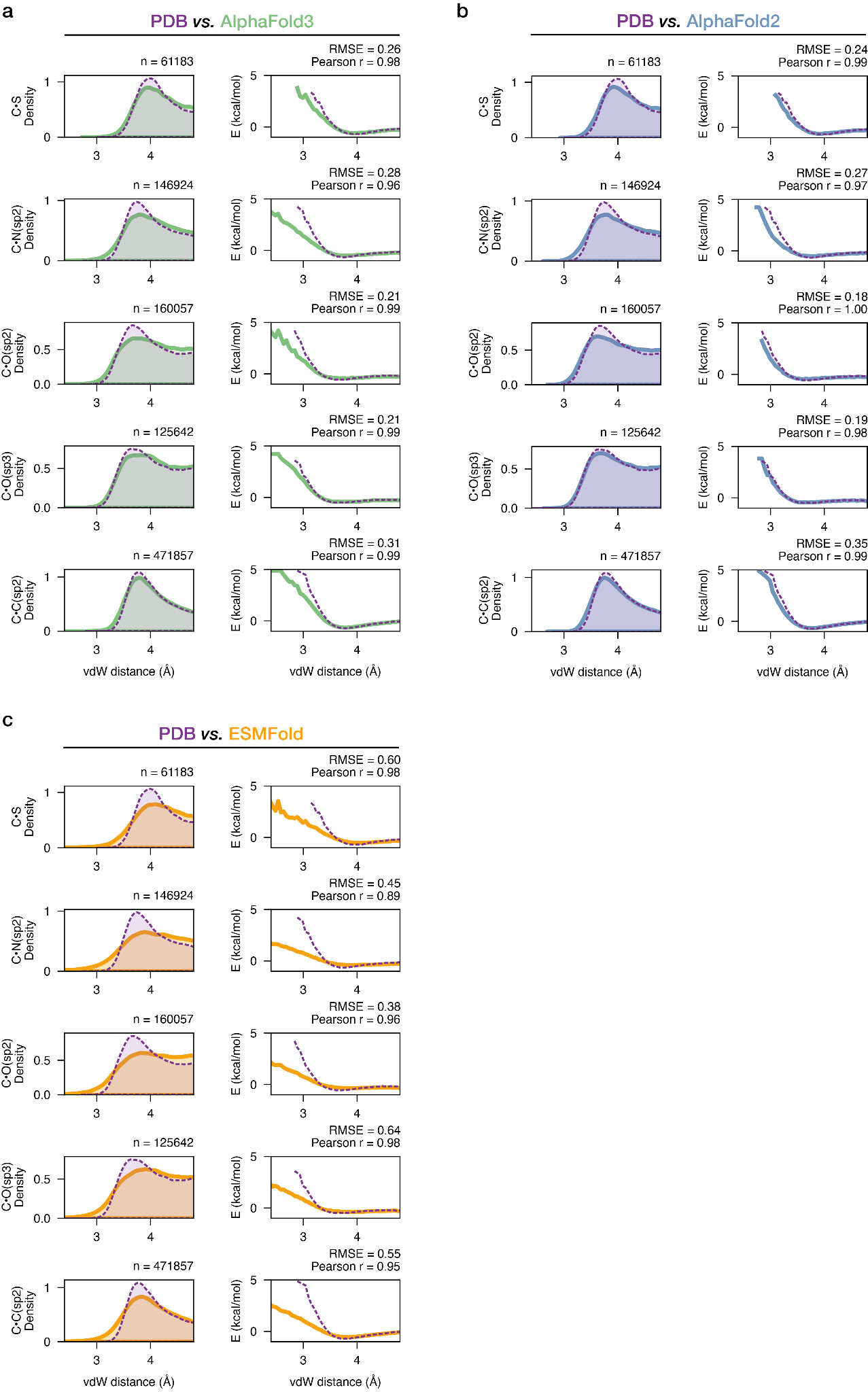

### Supplementary Fig. 8. Probability density distributions and corresponding energy functions for side-chain van der Waals interactions in (a) in AlphaFold3 predictions *versus* *Top2018* PDB structures, (b) in AlphaFold2 predictions *versus* *Top*2018 PDB structures and (c) in ESMFold predictions *versus* *Top*2018 PDB structures. The energy functions were obtained by inverting the distributions using the Boltzmann relationship (*Methods*). The models showed broader distributions that enrich shorter, sterically hindered contacts, with AlphaFold3 and ESMFold showing more pronounced differences than AlphaFold2.

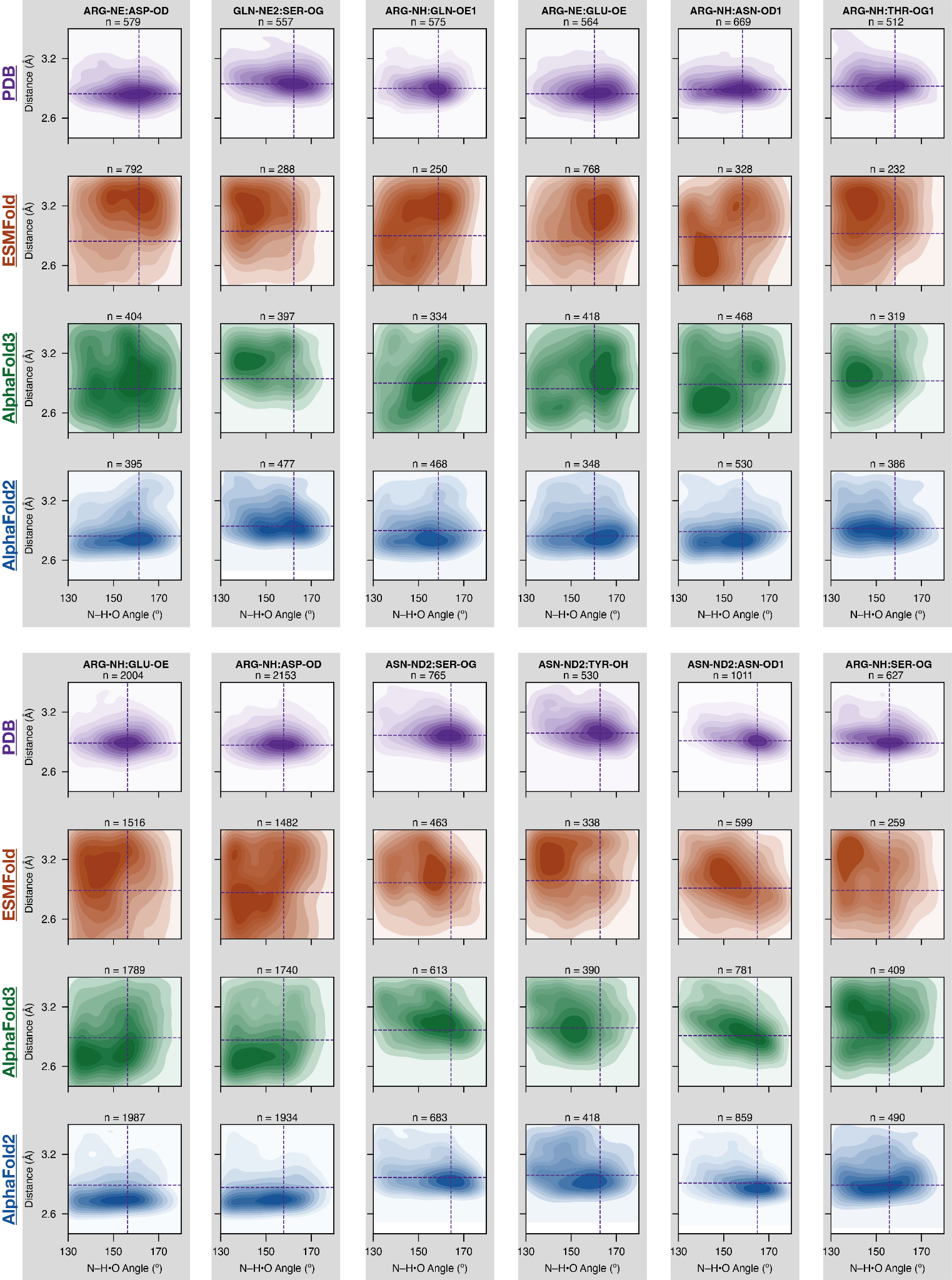

### Supplementary Fig. 9. Probability density distributions for hydrogen bond angles (N–H•O) and distances (N•O) for hydrogen bonds with sp^2^ hybridized donor atoms in *Top*2018 PDB structures (purple) *versus* those in ESMFold (orange), AlphaFold3 (green) and AlphaFold2 (blue). The color gradients indicate the probability mass under the contour in percentage (10% per contour), where the darker shades indicate more probable conformational states. The purple dashed lines indicate the position of the most populated state in the corresponding PDB distribution. For residue pairs that could form bidentate hydrogen bonds (between Arg and Glu/Asp residues), we selected pairs that form a single hydrogen bond to ensure unambiguous comparisons.

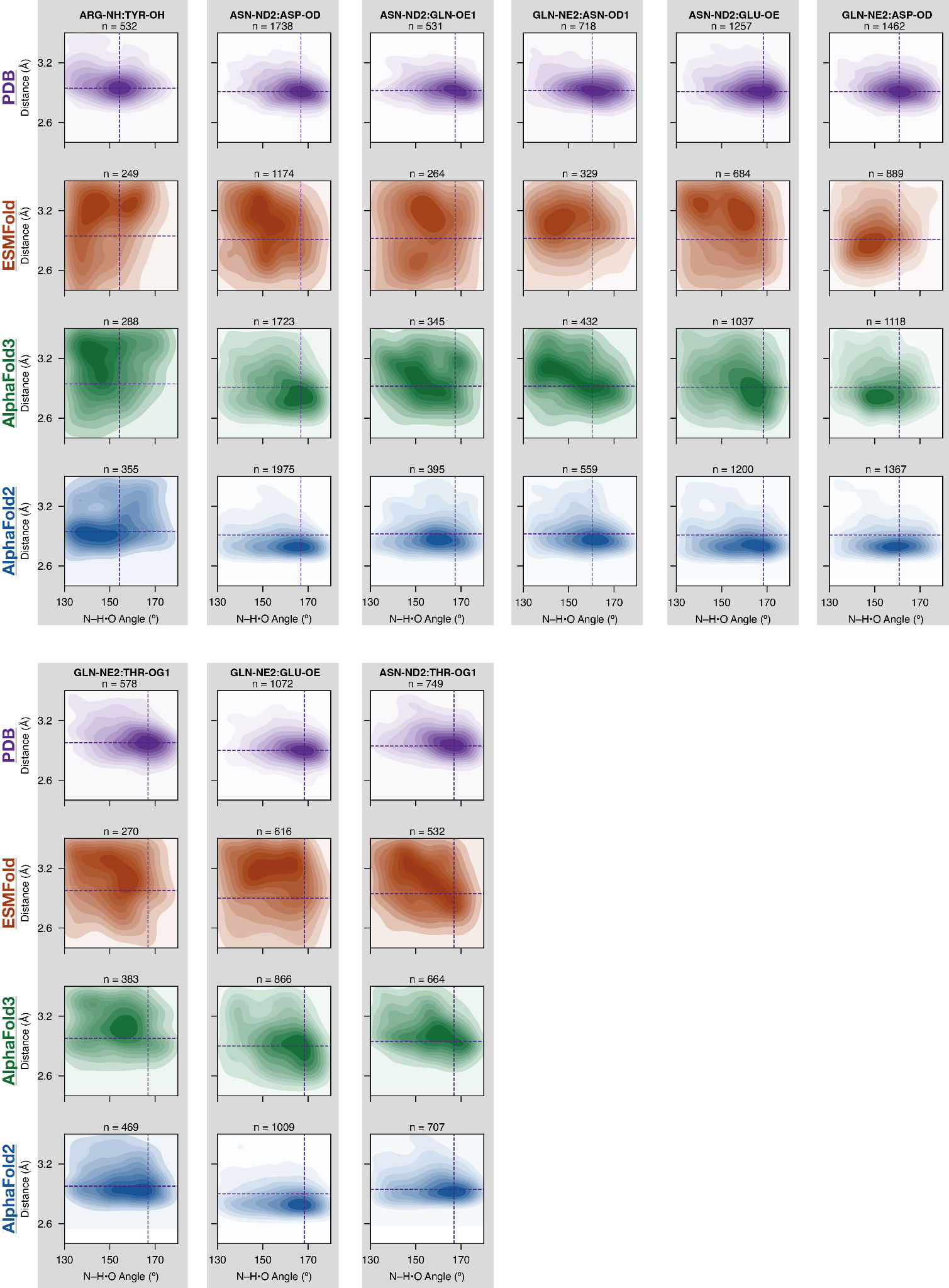

**Supplementary Fig. 9.** (continued)

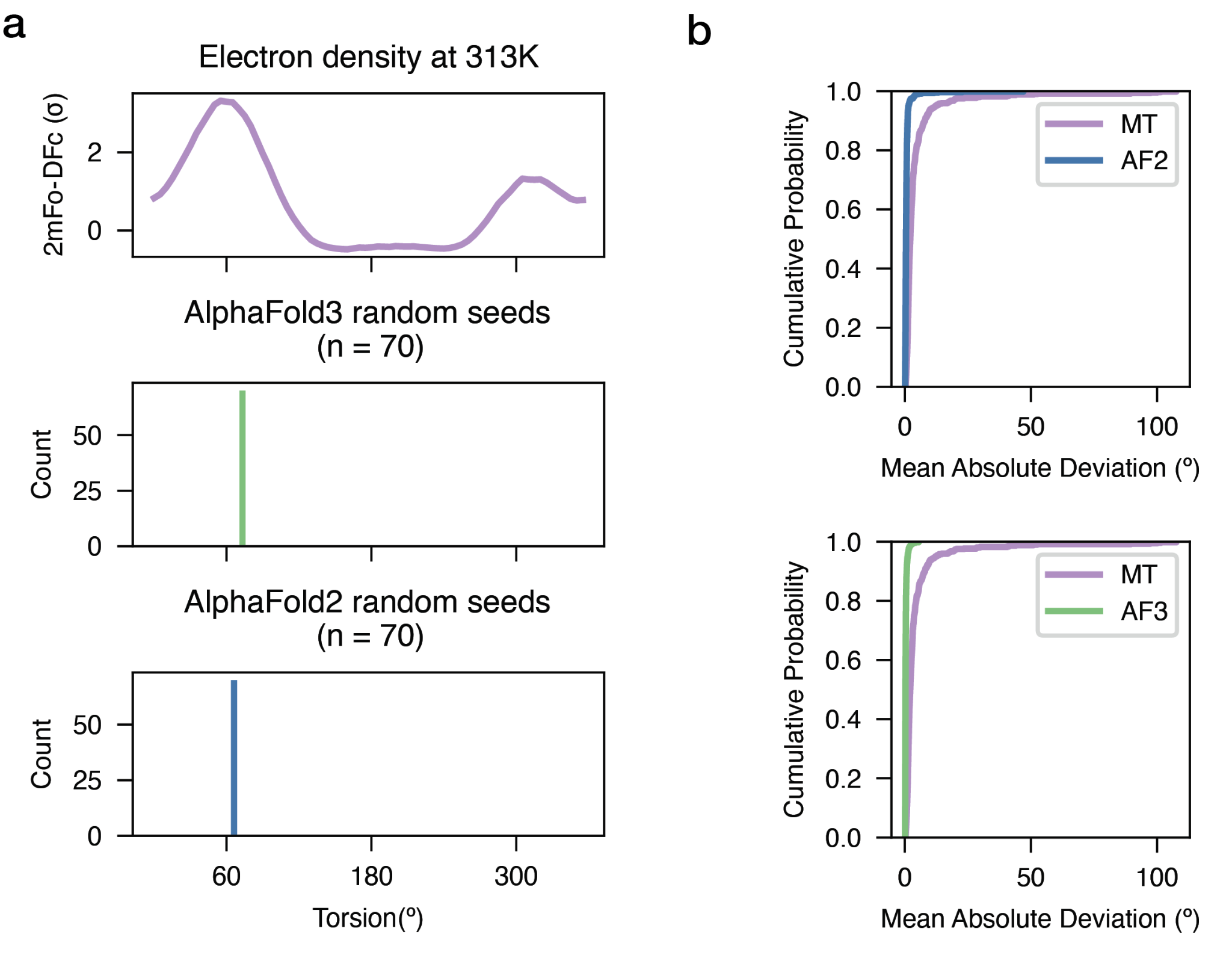

### Supplementary Fig. 10. AlphaFold2 random seed sampling for Cyclophilin A. (a) An example of the comparison performed for the multi-conformer side-chain of Ser21 from Cyclophilin A, also shown in the main text (Fig. 4A). The electron density map around Ser21 (2mFo-DFc; purple line) shows two conformers. The torsion distribution for this residue from the AlphaFold3 sampling is shown below in green and that from AlphaFold2 is shown in blue. (b) Cumulative distribution plot of the mean absolute deviation values (MAD) of all torsion distributions from MT ensemble (purple), AlphaFold2 sampling (blue) and AlphaFold3 sampling (green). Higher MAD (curves displaced to the right) indicates a higher level of spread of the distribution. Both AlphaFold2 and AlphaFold3 showed more restricted distributions (lower MAD) than the MT models (higher MAD).

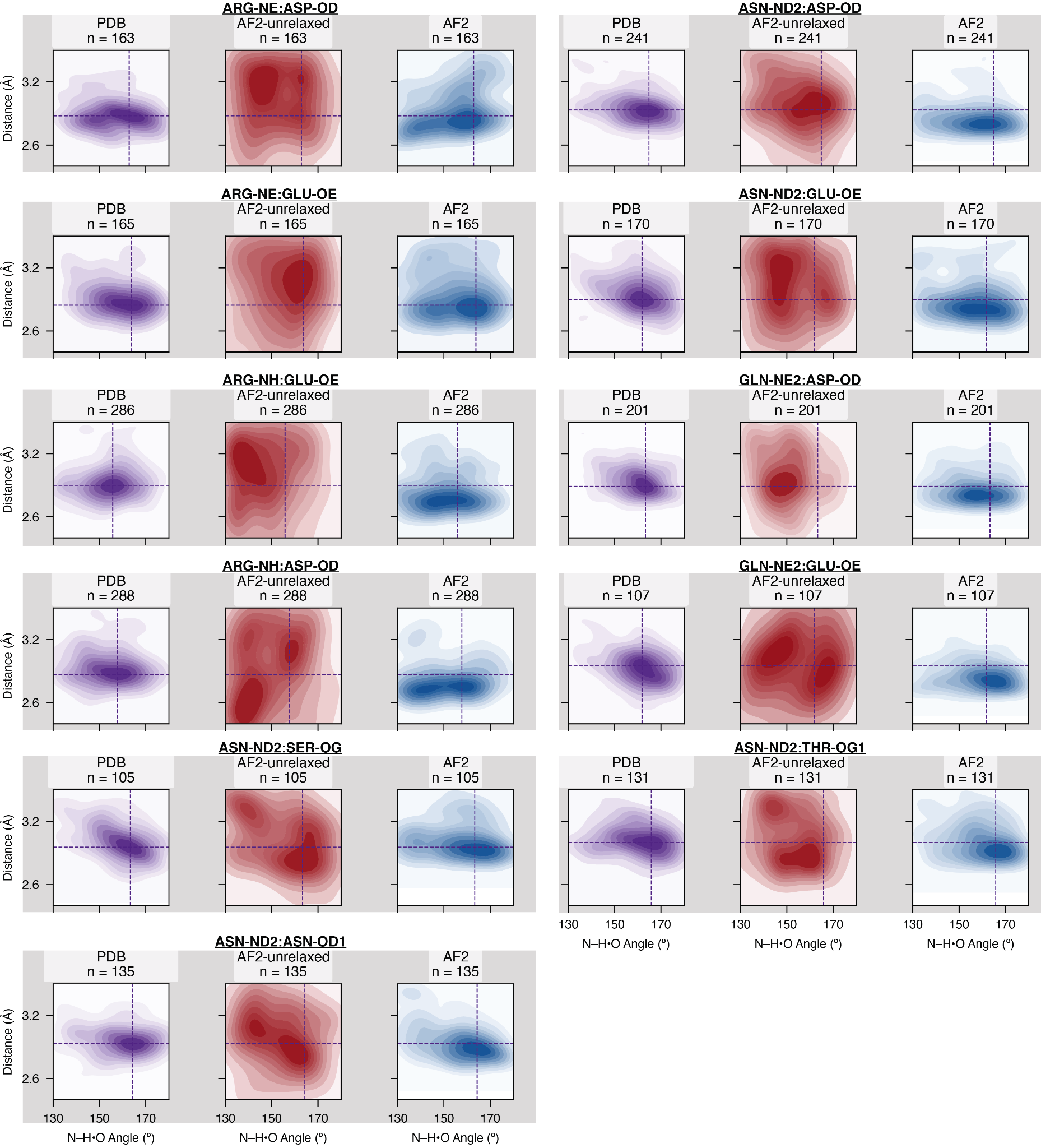

### Supplementary Fig. 11. Probability density distributions for hydrogen bond angles (N–H•O) and distances (N•O) for hydrogen bonds with sp^2^ hybridized donor atoms in *Top*2018 PDB structures (purple) *versus* those in unrelaxed AlphaFold2 predictions (red) and relaxed AlphaFold2 (blue). We performed unrelaxed AlphaFold2 predictions for the *Top*2018 PDB structures in the apo state (n = 1056) so that the sample sizes are smaller than the distributions from the full dataset shown in Supplementary Fig. 9; we randomly subsampled the PDB and relaxed AlphaFold2 predictions to the same sample size as the unrelaxed set, and only compared the distributions where the sample size is larger than 100. The color gradients indicate the probability mass under the contour in percentage (10% per contour), where the darker shades indicate more probable conformational states. The purple dashed lines indicate the position of the most populated state in the corresponding PDB distribution. For residue pairs that could form bidentate hydrogen bonds (between Arg and Glu/Asp residues), we selected pairs that form a single hydrogen bond to ensure unambiguous comparisons.

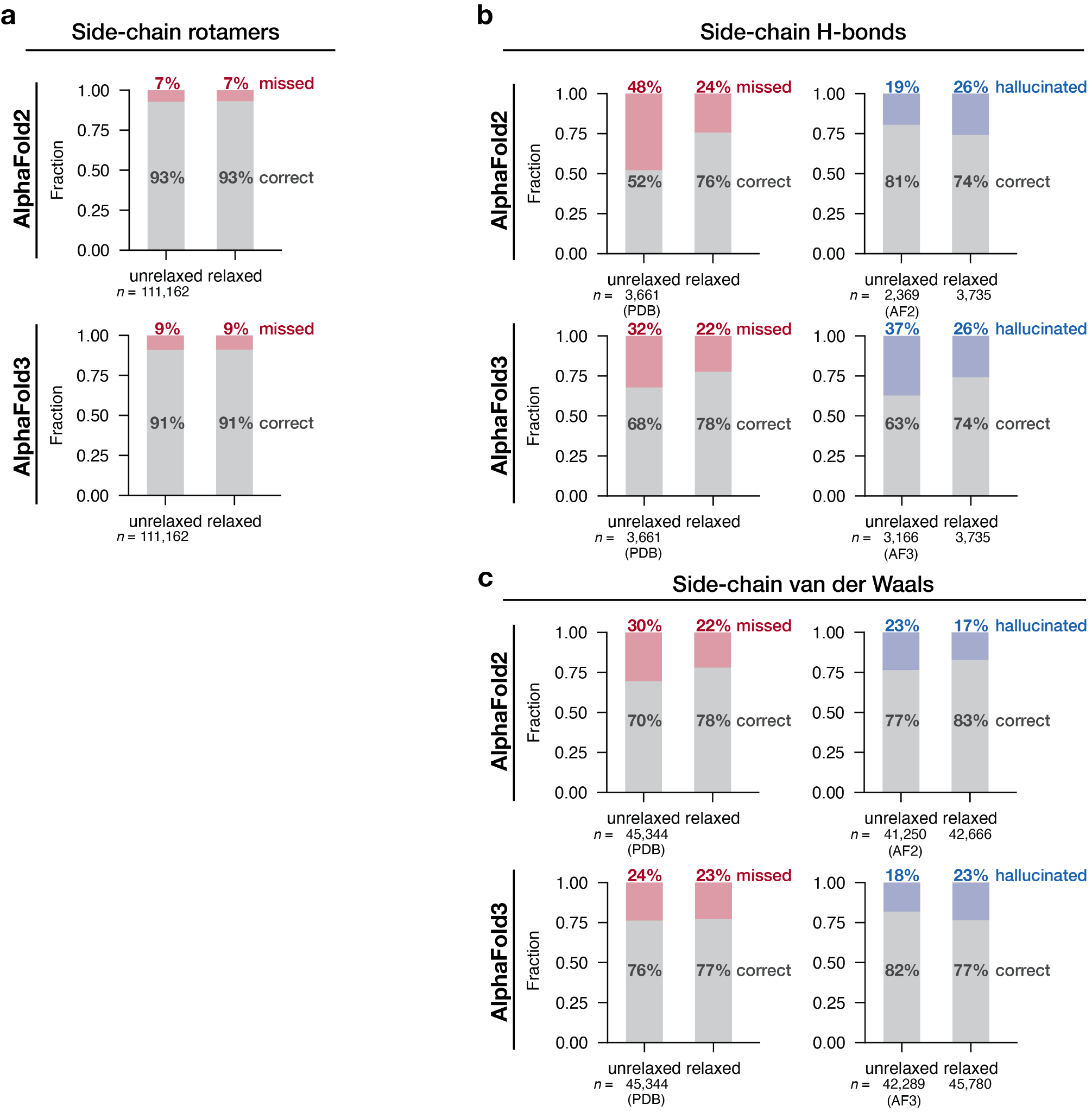

### Supplementary Fig. 12. Comparisons of the accuracy of model predictions before and after AMBER relaxation for (a) side-chain rotameric states, (b) side-chain•side-chain hydrogen bonds, and (c) sidechain•sidechain van der Waals interactions. For (a), the grey bars indicate the fraction of side-chain rotamers found in common between the PDB structures and the corresponding model predictions (“correct”). The fraction of interactions that were found in the PDB references but not found in model predictions is shown as red bars (“missed”). The annotated sample size is the total number of rotamers found in the PDB references (the same as that in the model predictions). For (b) and (c), the left subpanels are the same as in (a), showing the fraction of PDB interactions that are in common (“correct”) or missed by the models (“missed”). In the right subpanel, the fraction of interactions that were found in model predictions but not found in the PDB structures are shown as blue bars (“hallucinated”), and the total number of interactions found for each model is annotated below the bar.

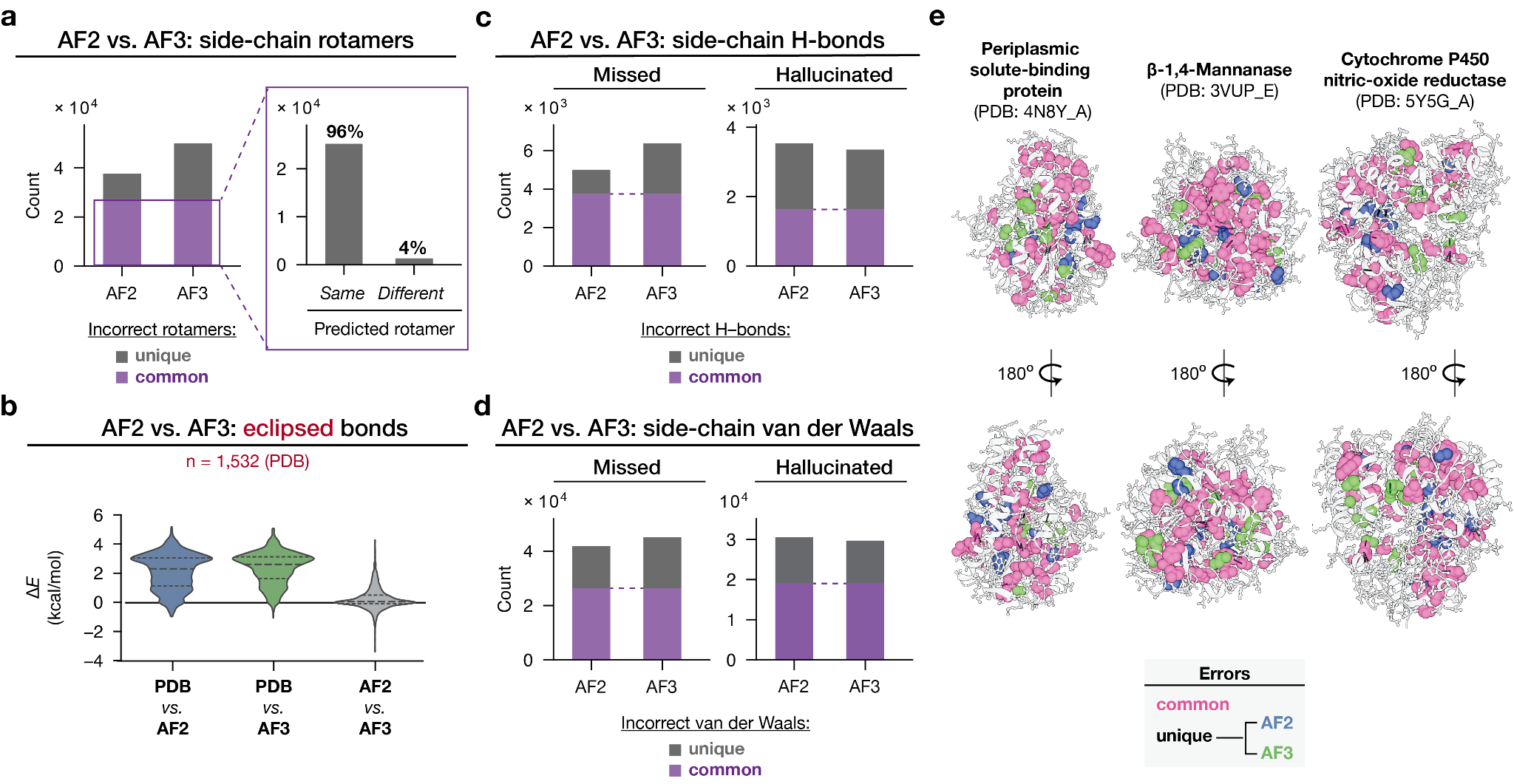

### Supplementary Fig. 13. Comparison of AlphaFold2 (after AMBER relaxation) and AlphaFold3 prediction errors. (a) Rotameric states errors in AlphaFold2 and AlphaFold3. Left: the number of incorrect rotamers that were common (purple) and unique (grey) errors. Right: within the shared-error subset, number of common *versus* different predicted (incorrect) rotameric states. (b) Energetic deviation (Δ*E*) for eclipsed side-chain bonds torsions for PDB conformers to the AlphaFold2 predictions; for PDB conformers *versus* AlphaFold3 predictions; and for AlphaFold2 *versus* AlphaFold3 predictions. (c) Side-chain hydrogen bond errors in AlphaFold2 and AlphaFold3 that were common (purple) and unique (grey). Left: hydrogen bonds identified in the PDB structures but not in the corresponding model predictions. Right: hydrogen bonds hallucinated in each model prediction. (d) Side-chain van der Waals errors in AlphaFold2 and AlphaFold3 that were common (purple) and unique (grey). Left: van der Waals interactions identified in the PDB structures but not in the corresponding model predictions. Right: van der Waals interactions hallucinated in model predictions. (e) Spatial distribution of common (pink) *versus* model-specific side-chain errors (blue: AlphaFold2; green: AlphaFold3), mapped onto three representative protein structures; 180° rotated views are shown for clarity.

### Supplementary Table 1. Literature synopsis for a selection of function-related predictions from deep-learning structure prediction models (AlphaFold2/3, RoseTTAFold, ESMFold) providing evidence for and against predictive abilities.

| **Task** | **How models were applied** | **Evidence for models’ capacity** | **Evidence against models’ capacity** |
| --- | --- | --- | --- |
| Small-molecule docking | • Using AF2 structures as receptor structures for docking^,34,62,63,83,84^ | • Docking to AF2 for new ligand discovery led to the identification of several new ligands not found when docking with experimental structures^85^. | • When docking known ligands to receptors, performance with AF2 models was similar to that with traditional homology models^34,83^, and both were worse than docking to experimental structures^62,84^.  • Side-chain misplacements in AF2 were found to hinder docking^63^; AF2 systematically underestimates binding pocket size^86^. |
| Protein-ligand complex structure prediction | • Using AF3 and RoseTTAFold-all-atoms to directly predict protein-ligand complex structures.^89^  • Using AF2 to discover cryptic pockets (*i.e.* pockets that are absent in the apo structure).^87^  • Using ESMFold-predicted structures to train a deep-learning model that predicts ligand binding sites on proteins.^88^ | • The accuracy of protein-ligand structures predicted by AF3 and RoseTTAFold-all-atoms was significantly higher than that of AutoVina^89^  •AF3 generated fewer physically implausible ligand poses than previous deep learning docking models^9,28^.  • Among 10 known proteins that form cryptic pockets, the AF2 ensembles of 6 contained pocket-opened states; in the other 4 cases, using AF2 ensembles as starting structures for molecular dynamics simulation accelerated the discovery of open-pocket states^87^. | • Accuracy of AF3 protein-ligand structures declined significantly on structures released after its training cutoff date, suggesting memorization instead of understanding of molecular interactions.^89^  • Accuracy of AF3 protein-peptide structure was found to be lower than the earlier models, AF-multimer and ColabFold.^90^  • Ligand docking accuracy for AF3 significantly drops as ligands become less similar to those seen in training^91^.  • Out of the AF3 prediction of 9 fatty-acid binding proteins, only 3 predicted the correct binding poses, with key hydrophobic stabilization interactions missing in the other 6.^92^  • The accuracy of RoseTTAFold-all-atoms predicted protein-ligand structures was significantly lower than that of AF3.^9^ |
| Protein-protein interaction | • Using AF-multimer and AF3 to directly predict protein-protein complex structures | • AF-multimer predicted accurate crystallographic homodimer geometries and its prediction confidence scores distinguished physiological homodimers from artificial complexes^93^.  • AF3 outperformed AF-multimer in antibody-antigen docking success rate and binding interface structural accuracy^9,94^. | • AF-multimer prediction quality of binding structure systematically decreases as the binding mode becomes more dynamic^95,96,97^.  • AF-multimer predicts ~40% of anti-parallel coiled-coil oligomeric states as parallel among a set of 216 experimental coiled-coil complex structures^98^.  • Binding free energy prediction using AF3 structures yielded 8.6% higher root-mean-square-error compared to that using experimental structures, when benchmarked against 8,338 experimental protein-protein binding free energy measurements^99^.  • Enhanced-sampling-based corrections can improve AF-multimer’s antigen-antibody structure prediction success rate from 20% to 43% on a set of 67 antibody-antigen structures, indicating binding interactions are recovered by physics-based downstream sampling rather than by the model itself^100^. |
| Prediction of alternative conformations | • Obtaining multiple conformations by repeated sampling AF2/3 and by increasing the depth of multiple-sequence alignment (MSA) and performing sequence clustering during the MSA^17–19,101^.  • Fine-tuning AF2 with Double Electron-Electron Resonance spectroscopy data can recover multiple alternative conformations^102^.  • Repurposing AlphaFold2’s internal side-chain representations to infer side-chain χ-angle distributions^50^. | • Several known fold-switched conformations were identified upon repeated sampling and upon deepening MSA^17,18^.  • The AF2-inferred side-chain χ-angle distributions were found to match that derived from experimental structures, as well as NMR coupling constants and molecular dynamics simulations^50^. | •AF2 did not predict two thirds of the fold-switched conformations among 92 known fold-switchers upon extensive resampling ^21,103^; AF2 predicted the lower energy conformations of fold switchers with higher confidence only 50% of the time, the same amount as expected by random chance^21^. |
| Predicting mutational effect | • Predicting stability or phenotype using the geometric deviation between the AF2 predicted wildtype (WT) and single mutant structures as a proxy ^14,104^.  • Applying structure-based variant-effect predictors (FoldX, Rosetta, DynaMut2, *etc*) on AF2 structures to predict mutation ΔΔG and comparing output scores for mutation ΔΔG obtained from traditional physics-based models^13^. | • The geometric deviation [referred to as “Effective Strain”, ES ^104^] was found to be correlated with fluorescence activity and folding free energy changes induced by point-mutations ^14,104^.  • Mutation ΔΔGscores computed by structure-based variant effect predictors from AF2 predictions correlated with deep-mutational-scanning data, and the results were similar to the scores computed from experimental structures^13^. | • The correlations in^14,104^ gave a low Pearson correlation coefficient of 0.35.  • AF2 predicted WT-like structure with high confidence when mutations well-known known to cause unfolding or fold-switching were introduced ^22,105^. The expected mutation-induced fold-switching was not recovered by molecular dynamics from AF2 predicted, WT-like mutant structures^105^.  • Rosetta-assigned mutation-induced free energy change correlated poorly with experimentally determined melting temperature (Pearson coefficient = -0.52 +/- 0.25 among 25 runs spread from -0.04 to -0.91)^106^. |
| Intrinsically disordered regions (IDRs) | • Using low AF2 prediction confidence score (pLDDT) as a signal for disorderness. | • A pLDDT threshold of 0.69 successfully identified most disordered regions across 662 proteins (precision = 0.83, recall = 0.78)^107^. | • Multiple sequence-based disorder predictors perform statistically better than AF2-pLDDT in identifying disordered residues in a set of 646 proteins that contain disordered regions^108,109^.  • AF2 assigned confident structures for 15% human IDRs^110^.  • The radiusRadius of gGyration radius of AF2 predicted IDRs substantially deviated from SAXS measured ones^111^.  • On a set of 42 protein-peptide complexes in which binding is mediated by IDRs, AF-Multimer correctly located the interaction site in only 17 complexes (~40% success)^112^, much lower than its 70% success rate on the full set of protein-protein complexes published during 2018-2021^113^.  • AF3 does not outperform AF2 on the Critical Assessment of Protein Intrinsic Disorder, third round (CAID3) benchmark^114^. |
| De novo protein design | • Repurposing and fine-tuning RoseTTAFold to become a protein structure generation model, RFdiffusion.^10^ | • RFdiffusion was able to generate a binder to influenza haemagglutinin, whose designed structure was experimentally validated with cryogenic electron microscopy.^10^ | • The binders designed by RFdiffusion bind with relatively low success rate (~20% overall success rate on five proteins with experimentally-solved receptor-binder complexes: Influenza A H1 HA, IL-7Rα, InsR, PD-L1 and TrkA).^10^ |

### Supplementary Table 2. Metrics used to train and evaluate current structure prediction models.

| **Model (year)** | **Main training loss** | **Primary evaluation metrics** | **Auxiliary loss terms** |
| --- | --- | --- | --- |
| AlphaFold2 (2021)^6^ | Backbone FAPE^a^ | Cɑ-RMSD^b^  Cɑ-lDDT^c^,  TM-score^d^ | distogram^e^, torsion angle^f^, structural violation^g^, matching experimentally resolved structures^h^ |
| RoseTTAFold (2021)^33^ | Contact + pLDDT^i^ + RMSD | Cɑ-RMSD, TM-score | N/A |
| ESMFold (2022)^7^ | Backbone FAPE | Cɑ-lDDT,  Cɑ-RMSD,  TM-score | distogram^e^, structural violation^g^ |
| RoseTTAFold-All-Atom (2023)^8^ | All-atom FAPE | Cɑ- and ligand-RMSD | distogram^e^, torsion angle^f^, structural violation^g^, masked token recovery^j^ |
| AlphaFold3 (2024)^9^ | diffusion^k^ + PDE^l^ + PAE^m^ | dock-Q^n^,  Cɑ- and ligand-RMSD | distogram^e^, matching experimetally resolved structures^h^ |

^a^FAPE (frame-aligned point error), a distance-based metric that measures the discrepancy between predicted and true atom positions after both are transformed into the local coordinate frame of a given residue, yielding a translation- and rotation-invariant error similar to aligned residue RMSD;

^b^Cɑ or ligand-RMSD, root-mean-square deviation of Cɑ or ligand atoms between aligned proteins;

^c^lDDT (local distance difference test), a per-atom metric that measures the fraction of nearby contacts with correctly predicted distances;

^d^TM (template-modelling)-score, a processed RMSD value such that it is normalized to chain length and capped by a maximum value, presumably making it more robust to local structural deviations and less biased by protein size than RMSD;

^e^An averaged cross-entropy loss over residue-pair distance bins;

^f^L2 loss on unit-circle representations of predicted backbone and side-chain torsions;

^g^Stereochemical penalties including bond-length violation loss, bond-angle violation loss, and clash / peptide-geometry violation terms, implemented with flat-bottom L1-style penalties beyond tolerance thresholds;
^h^Atom-wise binary cross-entropy loss predicting whether each atom would be experimentally resolved in a high-resolution structure;

^i^Predicted lDDT (pLDDT), model predictions of lDDT;
^j^A cross-entropy sequence reconstruction loss in which some input residue or biomolecular identity tokens are masked and the model is trained to predict the correct original token at those positions from the surrounding structural and sequence context.

^k^A noise-level-weighted combination of structures’ weighted aligned MSE term plus an auxiliary fine-tuning loss for bonded ligands / glycans
^l^A cross-entropy loss over 64 bins for the predicted aligned error between residue pairs, where the target is the alignment error of one token in the frame of another.
^m^A cross-entropy loss over bins for the absolute error in representative-atom distances between residue pairs.

^n^DockQ (protein–protein docking quality), a composite score that measures the accuracy of protein-protein contact interface, calculated by the weighted sum of three metrics: fraction of native contacts correctly predicted, ligand RMSD and interface residue Cɑ RMSD.

### Supplementary Table 3. Counts of interactions calculated from 3939 *Top*2018 PDB structures and the corresponding AlphaFold2 and AlphaFold3 predictions

| **Interaction type** | | **Count**  **(*Top*2018, all)** | **Count**  **(*Top*2018, buried*)** | **Count**  **(AF3, all)** | **Count**  **(AF3, buried)** | **Count**  **(AF2, all)** | **Count**  **(AF2, buried)** | **Count**  **(ESMFold, all)** | **Count**  **(ESMFold, buried)** |
| --- | --- | --- | --- | --- | --- | --- | --- | --- | --- |
| **Torsion** | φ | 638,629 | 410,671 | 638,629 | 410,671 | 638,629 | 410,671 | 638,629 | 410,671 |
|  | ψ | 638,629 | 410,319 | 638,629 | 410,319 | 638,629 | 410,319 | 638,629 | 410,319 |
|  | χ^1^ | 605,325 | 387,275 | 605,325 | 387,275 | 605,325 | 387,275 | 605,325 | 387,275 |
|  | χ^2^ | 442,599 | 276,349 | 442,599 | 276,349 | 442,599 | 276,349 | 442,599 | 276,349 |
|  | χ^3^ | 119,137 | 56,883 | 119,137 | 56,883 | 119,137 | 56,883 | 119,137 | 56,883 |
|  | χ^4^ | 49,869 | 20,911 | 49,869 | 20,911 | 49,869 | 20,911 | 49,869 | 20,911 |
|  | *total* | 2,494,188 | 1,562,408 | 2,494,188 | 1,562,408 | 2,494,188 | 1,562,408 | 2,494,188 | 1,562,408 |
| **H-bond** | side-chain | 45,682 | 20,766 | 41,593 | 18,388 | 44,567 | 19,759 | 37,869 | 12,603 |
|  | backbone | 393,766 | 242,269 | 393,731 | 242,044 | 392,743 | 241,731 | 550,962 | 238,936 |
|  | *total* | 439,448 | 263,035 | 435,324 | 260,432 | 437,310 | 261,490 | 588,831 | 251,539 |
| **vdW** | C•sp² C | 228,926 | 199,026 | 228,163 | 197,992 | 228,049 | 198,071 | 276,880 | 211,518 |
|  | C•sp² O | 75,506 | 40,896 | 75,159 | 40,367 | 74,960 | 40,373 | 100,462 | 38,966 |
|  | C•sp³ O | 76,370 | 56,773 | 76,826 | 56,725 | 76,879 | 56,830 | 91,638 | 54,907 |
|  | C•sp² N | 76,195 | 47,534 | 74,522 | 46,327 | 74,600 | 46,526 | 95,520 | 43,845 |
|  | C•S | 35,186 | 31,644 | 35,737 | 32,092 | 35,579 | 31,951 | 47,684 | 31,625 |
|  | *total* | 492,183 | 375,873 | 490,407 | 370,503 | 490,067 | 373,751 | 612,184 | 380,861 |
| **Total** | | **3,425,819** | **2,201,316** | **3,419,919** | **2,193,343** | **3,421,565** | **2,197,649** | **3,695,203** | **2,194,808** |
| Covalent (AF3 only) | | 5,155,647 | – | 5,155,647 | **–** | **–** | **–** | **–** | **–** |

*relative solvent accessibility ≤ 0.25

### Supplementary Table 4. The most favored covalent bond lengths and the median absolute deviation (MAD) of the covalent bond length distributions from Top2018 PDB structures and from AlphaFold3 predictions.

| **Covalent bond type** | ***n*** | **Most favored distance**  **(peak) (Å)** | | **Median abs. deviation**  **(MAD) (Å)** | | **Δ (Å)**  **(PDB – AlphaFold3)** | |
| --- | --- | --- | --- | --- | --- | --- | --- |
|  |  | *PDB* | *AlphaFold3* | *PDB* | *AlphaFold3* | *peak* | *MAD* |
| ALA_C(sp3)_C(sp3) | 77046 | 1.526 | 1.532 | 0.007 | 0.003 | -0.006 | 0.005 |
| ARG_C(sp2)_N(sp2) | 56423 | 1.327 | 1.301 | 0.004 | 0.005 | 0.026 | -0.001 |
| ARG_C(sp3)_C(sp3) | 85193 | 1.530 | 1.534 | 0.008 | 0.004 | -0.004 | 0.004 |
| ARG_C(sp3)_N(sp2) | 28234 | 1.461 | 1.472 | 0.004 | 0.005 | -0.010 | 0.000 |
| ARG_N(sp2)_C(sp2) | 28213 | 1.330 | 1.322 | 0.003 | 0.005 | 0.008 | -0.002 |
| ASN_C(sp2)_N(sp2) | 32088 | 1.328 | 1.310 | 0.005 | 0.003 | 0.019 | 0.002 |
| ASN_C(sp2)_O(sp2) | 32093 | 1.232 | 1.220 | 0.005 | 0.004 | 0.012 | 0.001 |
| ASN_C(sp3)_C(sp2) | 32100 | 1.516 | 1.516 | 0.007 | 0.003 | 0.000 | 0.004 |
| ASN_C(sp3)_C(sp3) | 32131 | 1.531 | 1.536 | 0.005 | 0.003 | -0.005 | 0.002 |
| ASP_C(sp2)_O(sp2) | 86494 | 1.249 | 1.251 | 0.004 | 0.004 | -0.001 | 0.001 |
| ASP_C(sp3)_C(sp2) | 43257 | 1.518 | 1.533 | 0.007 | 0.004 | -0.014 | 0.003 |
| ASP_C(sp3)_C(sp3) | 43334 | 1.532 | 1.537 | 0.005 | 0.004 | -0.005 | 0.002 |
| CYS_C(sp3)_C(sp3) | 10408 | 1.530 | 1.533 | 0.005 | 0.003 | -0.003 | 0.002 |
| CYS_C(sp3)_S(sp3) | 10407 | 1.808 | 1.813 | 0.011 | 0.004 | -0.006 | 0.007 |
| GLN_C(sp2)_N(sp2) | 22078 | 1.329 | 1.311 | 0.005 | 0.004 | 0.018 | 0.001 |
| GLN_C(sp2)_O(sp2) | 22080 | 1.232 | 1.220 | 0.004 | 0.004 | 0.012 | 0.001 |
| GLN_C(sp3)_C(sp2) | 22083 | 1.517 | 1.525 | 0.007 | 0.004 | -0.009 | 0.003 |
| GLN_C(sp3)_C(sp3) | 44370 | 1.530 | 1.535 | 0.007 | 0.003 | -0.005 | 0.004 |
| GLU_C(sp2)_O(sp2) | 67159 | 1.250 | 1.250 | 0.004 | 0.004 | 0.000 | 0.000 |
| GLU_C(sp3)_C(sp2) | 33591 | 1.518 | 1.538 | 0.007 | 0.005 | -0.020 | 0.002 |
| GLU_C(sp3)_C(sp3) | 67873 | 1.531 | 1.537 | 0.007 | 0.004 | -0.006 | 0.003 |
| HIS_C(sp3)_C(sp2) | 17585 | 1.496 | 1.506 | 0.003 | 0.003 | -0.009 | 0.000 |
| HIS_C(sp3)_C(sp3) | 17602 | 1.531 | 1.533 | 0.005 | 0.003 | -0.003 | 0.003 |
| ILE_C(sp3)_C(sp3) | 178515 | 1.532 | 1.544 | 0.010 | 0.005 | -0.012 | 0.006 |
| LEU_C(sp3)_C(sp3) | 285169 | 1.531 | 1.538 | 0.007 | 0.005 | -0.007 | 0.002 |
| LYS_C(sp3)_C(sp3) | 88323 | 1.523 | 1.534 | 0.008 | 0.005 | -0.011 | 0.003 |
| LYS_C(sp3)_N(sp2) | 21664 | 1.490 | 1.482 | 0.007 | 0.005 | 0.008 | 0.002 |
| MET_C(sp3)_C(sp3) | 27145 | 1.530 | 1.533 | 0.008 | 0.003 | -0.003 | 0.004 |
| MET_C(sp3)_S(sp3) | 13561 | 1.805 | 1.815 | 0.009 | 0.004 | -0.010 | 0.005 |
| MET_S(sp3)_C(sp3) | 13559 | 1.792 | 1.809 | 0.009 | 0.004 | -0.017 | 0.005 |
| PHE_C(sp3)_C(sp2) | 33769 | 1.503 | 1.515 | 0.006 | 0.003 | -0.012 | 0.003 |
| PHE_C(sp3)_C(sp3) | 33774 | 1.532 | 1.533 | 0.005 | 0.003 | 0.000 | 0.002 |
| PRO_C(sp3)_C(sp3) | 115333 | 1.532 | 1.528 | 0.018 | 0.003 | 0.004 | 0.015 |
| SER_C(sp3)_C(sp3) | 45376 | 1.531 | 1.531 | 0.005 | 0.003 | 0.000 | 0.002 |
| SER_C(sp3)_O(sp3) | 45359 | 1.417 | 1.414 | 0.005 | 0.003 | 0.003 | 0.002 |
| THR_C(sp3)_C(sp3) | 91840 | 1.527 | 1.534 | 0.011 | 0.004 | -0.006 | 0.006 |
| THR_C(sp3)_O(sp3) | 45918 | 1.433 | 1.407 | 0.003 | 0.003 | 0.026 | 0.000 |
| TRP_C(sp3)_C(sp2) | 12511 | 1.499 | 1.496 | 0.010 | 0.003 | 0.003 | 0.006 |
| TRP_C(sp3)_C(sp3) | 12511 | 1.532 | 1.531 | 0.006 | 0.003 | 0.001 | 0.003 |
| TYR_C(sp3)_C(sp2) | 29736 | 1.512 | 1.515 | 0.006 | 0.003 | -0.003 | 0.003 |
| TYR_C(sp3)_C(sp3) | 29740 | 1.532 | 1.532 | 0.006 | 0.003 | -0.001 | 0.003 |
| VAL_C(sp3)_C(sp3) | 181918 | 1.524 | 1.534 | 0.011 | 0.004 | -0.009 | 0.007 |
| bb*_C(sp2)_C(sp3) | 753667 | 1.525 | 1.533 | 0.006 | 0.004 | -0.008 | 0.002 |
| bb_C(sp2)_O(sp2) | 607111 | 1.232 | 1.230 | 0.005 | 0.003 | 0.002 | 0.002 |
| bb_C(sp3)_N(sp2) | 645554 | 1.459 | 1.464 | 0.006 | 0.004 | -0.005 | 0.002 |
| bb_N(sp2)_C(sp2) | 515861 | 1.330 | 1.334 | 0.004 | 0.005 | -0.004 | -0.002 |

*backbone

### Supplementary Table 5. The most favored covalent bond angles and the median absolute deviation (MAD) of the covalent bond angle distributions from *Top*2018 PDB structures and from AlphaFold3 predictions.

| **Covalent bond type** | ***n*** | **Most favored distance**  **(peak) (Å)** | | **Median abs. deviation**  **(MAD) (Å)** | | **Δ (Å)**  **(PDB – AlphaFold3)** | |
| --- | --- | --- | --- | --- | --- | --- | --- |
|  |  | *PDB* | *AlphaFold3* | *PDB* | *AlphaFold3* | *peak* | *MAD* |
| ARG:CA-CB-CG | 28326 | 113.9 | 112.3 | 1.1 | 0.8 | 1.7 | 0.3 |
| ARG:CB-CG-CD | 28279 | 111.8 | 111.3 | 1.1 | 0.8 | 0.5 | 0.3 |
| ARG:CD-NE-CZ | 28213 | 124.5 | 126.5 | 0.5 | 0.8 | -2.0 | -0.2 |
| ARG:CG-CD-NE | 28234 | 111.8 | 111.7 | 1.2 | 0.7 | 0.2 | 0.5 |
| ARG:NE-CZ-NH | 56423 | 120.0 | 120.2 | 0.5 | 1.1 | -0.1 | -0.6 |
| ASN:CA-CB-CG | 32100 | 112.6 | 112.8 | 0.4 | 0.7 | -0.2 | -0.3 |
| ASN:CB-CG-ND | 32088 | 116.5 | 117.9 | 0.4 | 0.5 | -1.4 | -0.1 |
| ASN:CB-CG-OD | 32093 | 120.9 | 122.5 | 0.4 | 0.7 | -1.6 | -0.3 |
| ASP:CA-CB-CG | 43257 | 112.9 | 113.9 | 0.5 | 1.0 | -1.0 | -0.5 |
| ASP:CB-CG-OD | 86494 | 118.4 | 118.3 | 0.9 | 0.9 | 0.1 | 0.0 |
| CYS:CA-CB-SG | 10407 | 113.9 | 110.6 | 1.1 | 1.2 | 3.3 | -0.1 |
| GLN:CA-CB-CG | 22113 | 113.9 | 112.6 | 1.0 | 0.8 | 1.4 | 0.2 |
| GLN:CB-CG-CD | 22083 | 112.7 | 112.8 | 0.8 | 0.6 | -0.1 | 0.2 |
| GLN:CG-CD-NE | 22078 | 116.5 | 118.4 | 0.3 | 0.5 | -1.9 | -0.2 |
| GLN:CG-CD-OE | 22080 | 120.9 | 122.8 | 0.3 | 0.7 | -1.9 | -0.4 |
| GLU:CA-CB-CG | 33714 | 114.0 | 113.2 | 1.1 | 0.9 | 0.7 | 0.2 |
| GLU:CB-CG-CD | 33591 | 113.3 | 113.2 | 0.8 | 0.9 | 0.1 | 0.0 |
| GLU:CG-CD-OE | 67159 | 118.5 | 118.6 | 0.7 | 0.9 | -0.1 | -0.2 |
| HIS:CA-CB-CG | 17585 | 113.7 | 113.2 | 0.5 | 0.8 | 0.5 | -0.2 |
| ILE:CA-CB-CG | 89258 | 110.4 | 110.4 | 0.7 | 0.8 | 0.0 | -0.1 |
| ILE:CB-CG1-CD | 44624 | 114.0 | 114.0 | 0.7 | 0.5 | 0.0 | 0.2 |
| LEU:CA-CB-CG | 71288 | 116.0 | 114.8 | 1.3 | 0.6 | 1.2 | 0.8 |
| LEU:CB-CG-CD | 142567 | 110.0 | 109.4 | 1.0 | 1.4 | 0.6 | -0.5 |
| LYS:CA-CB-CG | 22067 | 113.9 | 112.5 | 1.0 | 1.0 | 1.4 | 0.0 |
| LYS:CB-CG-CD | 21864 | 111.8 | 111.0 | 1.0 | 1.1 | 0.8 | -0.1 |
| LYS:CD-CE-NZ | 21664 | 111.8 | 111.7 | 1.1 | 0.9 | 0.0 | 0.2 |
| LYS:CG-CD-CE | 21706 | 111.4 | 111.2 | 0.9 | 1.3 | 0.2 | -0.4 |
| MET:CA-CB-CG | 13562 | 114.0 | 112.6 | 1.0 | 0.6 | 1.4 | 0.3 |
| MET:CB-CG-SD | 13561 | 113.0 | 114.3 | 1.4 | 1.0 | -1.3 | 0.4 |
| MET:CG-SD-CE | 13559 | 100.8 | 100.7 | 0.9 | 0.6 | 0.1 | 0.3 |
| PHE:CA-CB-CG | 33769 | 113.8 | 113.7 | 0.6 | 0.7 | 0.1 | -0.1 |
| PRO:CA-CB-CG | 38444 | 104.4 | 103.3 | 0.9 | 0.4 | 1.1 | 0.5 |
| PRO:CB-CG-CD | 38443 | 105.4 | 102.9 | 1.3 | 0.3 | 2.5 | 1.0 |
| PRO:CG-CD-N | 38443 | 103.1 | 105.4 | 0.5 | 0.3 | -2.3 | 0.2 |
| SER:CA-CB-OG | 45359 | 111.0 | 110.9 | 0.8 | 0.6 | 0.1 | 0.1 |
| THR:CA-CB-CG | 45917 | 110.8 | 112.5 | 0.6 | 0.6 | -1.8 | 0.0 |
| THR:CA-CB-OG | 45918 | 109.4 | 109.8 | 0.5 | 0.7 | -0.3 | -0.2 |
| TRP:CA-CB-CG | 12511 | 114.1 | 114.2 | 1.4 | 0.9 | -0.1 | 0.5 |
| TYR:CA-CB-CG | 29736 | 113.4 | 113.8 | 1.4 | 0.8 | -0.4 | 0.6 |
| VAL:CA-CB-CG | 121272 | 110.4 | 111.1 | 0.6 | 0.7 | -0.8 | -0.1 |
| bb*:CA-C-O | 607111 | 120.6 | 120.3 | 0.5 | 0.6 | 0.3 | -0.1 |
| bb:CA-N-C | 515861 | 121.4 | 123.2 | 0.9 | 0.8 | -1.9 | 0.1 |
| bb:CB-CA-C | 684157 | 110.2 | 111.5 | 0.9 | 0.8 | -1.3 | 0.1 |
| bb:CB-CA-N | 607111 | 110.7 | 110.1 | 0.8 | 0.8 | 0.5 | 0.0 |
| bb:N-CA-C | 607111 | 111.1 | 111.6 | 1.6 | 1.2 | -0.5 | 0.4 |

*backbone

### Supplementary Table 6. The most favored van der Waals distance (a) and the median absolute deviation (MAD) (b) of the van der Waals distributions from *Top*2018 PDB structures and from AlphaFold2 AlphaFold3 and ESMFold predictions.

**(a)**

| **vdW type** | | ***n*** | **Most favored distance (peak) (Å)** | | | | **Δpeak (Å)** | | |
| --- | --- | --- | --- | --- | --- | --- | --- | --- | --- |
|  |  |  | *PDB* | *AF2* | *AF3* | *ESM* | *PDB–AF2* | *PDB–AF3* | *PDB–ESM* |
| C•S | CYS:SG | 13699 | 3.96 | 3.89 | 3.91 | 3.97 | 0.07 | 0.05 | -0.01 |
|  | MET:SD | 21487 | 4.02 | 3.96 | 4.12 | 4.30 | 0.05 | -0.10 | -0.28 |
| C•C(sp2) | ARG:CZ | 12729 | 3.81 | 3.84 | 3.95 | 4.02 | -0.03 | -0.14 | -0.21 |
|  | ASN:CG | 14311 | 4.26 | 4.42 | 4.26 | 4.25 | -0.16 | -0.01 | 0.01 |
|  | ASP:CG | 14522 | 4.43 | 4.27 | 4.22 | 4.27 | 0.15 | 0.20 | 0.16 |
|  | GLN:CD | 11327 | 4.16 | 4.28 | 4.31 | 4.46 | -0.12 | -0.15 | -0.30 |
|  | GLU:CD | 12862 | 4.33 | 4.50 | 4.46 | 4.46 | -0.16 | -0.13 | -0.13 |
|  | HIS:CD | 8236 | 3.85 | 3.80 | 3.85 | 3.84 | 0.05 | 0.00 | 0.02 |
|  | HIS:CG | 4425 | 3.84 | 3.80 | 3.89 | 3.90 | 0.03 | -0.05 | -0.06 |
|  | PHE:CD | 25773 | 3.80 | 3.79 | 3.81 | 3.81 | 0.02 | 0.00 | -0.01 |
|  | PHE:CE | 31616 | 3.76 | 3.74 | 3.76 | 3.77 | 0.02 | 0.00 | -0.01 |
|  | PHE:CG | 4514 | 3.71 | 3.69 | 3.72 | 3.72 | 0.02 | -0.01 | -0.02 |
|  | PHE:CZ | 15421 | 3.77 | 3.76 | 3.81 | 3.76 | 0.02 | -0.03 | 0.02 |
|  | TRP:CD | 4845 | 3.69 | 3.67 | 3.67 | 3.74 | 0.02 | 0.01 | -0.06 |
|  | TRP:CE | 6967 | 3.74 | 3.75 | 3.73 | 3.77 | -0.02 | 0.00 | -0.03 |
|  | TRP:CG | 1247 | 3.73 | 3.71 | 3.70 | 3.75 | 0.02 | 0.03 | -0.02 |
|  | TRP:CH | 4579 | 3.79 | 3.73 | 3.73 | 3.84 | 0.06 | 0.06 | -0.05 |
|  | TRP:CZ | 8452 | 3.78 | 3.73 | 3.75 | 3.72 | 0.05 | 0.03 | 0.06 |
|  | TYR:CD | 18991 | 3.79 | 3.77 | 3.80 | 3.84 | 0.02 | -0.01 | -0.05 |
|  | TYR:CE | 18316 | 3.72 | 3.71 | 3.71 | 3.76 | 0.01 | 0.01 | -0.04 |
|  | TYR:CG | 3550 | 3.71 | 3.68 | 3.70 | 3.77 | 0.03 | 0.01 | -0.06 |
|  | TYR:CZ | 6243 | 3.67 | 3.62 | 3.65 | 3.61 | 0.04 | 0.02 | 0.06 |
| C•N(sp2) | ARG:NE | 6294 | 3.73 | 4.46 | 4.01 | 4.63 | -0.73 | -0.28 | -0.89 |
|  | ARG:NH | 16376 | 3.70 | 3.63 | 3.71 | 3.86 | 0.06 | -0.01 | -0.16 |
|  | ASN:ND | 15935 | 3.80 | 3.79 | 3.86 | 3.83 | 0.00 | -0.06 | -0.04 |
|  | GLN:NE | 12472 | 3.84 | 4.06 | 4.03 | 4.35 | -0.22 | -0.20 | -0.52 |
|  | HIS:ND | 6443 | 3.79 | 3.77 | 3.79 | 3.83 | 0.02 | 0.00 | -0.04 |
|  | HIS:NE | 8412 | 3.75 | 3.80 | 3.83 | 3.85 | -0.05 | -0.09 | -0.10 |
|  | TRP:NE | 10263 | 3.79 | 3.83 | 3.82 | 4.00 | -0.05 | -0.03 | -0.21 |
| C•O(sp2) | ASN:OD | 17152 | 3.62 | 3.63 | 3.66 | 4.68 | -0.01 | -0.04 | -1.06 |
|  | ASP:OD | 25044 | 3.64 | 3.57 | 3.66 | 4.80 | 0.07 | -0.02 | -1.16 |
|  | GLN:OE | 13017 | 3.72 | 3.77 | 4.05 | 4.12 | -0.05 | -0.33 | -0.40 |
|  | GLU:OE | 20293 | 3.71 | 3.72 | 3.84 | 4.01 | -0.01 | -0.13 | -0.30 |
| C•O(sp3) | SER:OG | 27276 | 3.63 | 3.67 | 3.68 | 3.91 | -0.03 | -0.05 | -0.28 |
|  | THR:OG | 24623 | 3.64 | 3.70 | 3.90 | 3.91 | -0.06 | -0.26 | -0.27 |
|  | TYR:OH | 24471 | 3.72 | 3.73 | 3.86 | 4.03 | -0.01 | -0.14 | -0.31 |

# (b)

| **vdW type** | | ***n*** | **Median abs. deviation (MAD) (Å)** | | | | **ΔMAD (Å)** | | |
| --- | --- | --- | --- | --- | --- | --- | --- | --- | --- |
|  |  |  | *PDB* | *AF2* | *AF3* | *ESM* | *PDB–AF2* | *PDB–AF3* | *PDB–ESM* |
| C•S | CYS:SG | 13699 | 0.26 | 0.28 | 0.27 | 0.30 | -0.02 | -0.01 | -0.04 |
|  | MET:SD | 21487 | 0.29 | 0.34 | 0.34 | 0.35 | -0.05 | -0.05 | -0.06 |
| C•C(sp2) | ARG:CZ | 12729 | 0.32 | 0.36 | 0.34 | 0.39 | -0.03 | -0.02 | -0.06 |
|  | ASN:CG | 14311 | 0.32 | 0.32 | 0.31 | 0.32 | 0.00 | 0.01 | 0.00 |
|  | ASP:CG | 14522 | 0.34 | 0.34 | 0.33 | 0.33 | 0.01 | 0.01 | 0.02 |
|  | GLN:CD | 11327 | 0.33 | 0.33 | 0.33 | 0.34 | 0.00 | 0.00 | -0.01 |
|  | GLU:CD | 12862 | 0.35 | 0.34 | 0.34 | 0.35 | 0.01 | 0.01 | 0.01 |
|  | HIS:CD | 8236 | 0.27 | 0.30 | 0.30 | 0.33 | -0.03 | -0.02 | -0.06 |
|  | HIS:CG | 4425 | 0.40 | 0.38 | 0.39 | 0.40 | 0.01 | 0.01 | 0.00 |
|  | PHE:CD | 25773 | 0.26 | 0.28 | 0.28 | 0.30 | -0.02 | -0.02 | -0.04 |
|  | PHE:CE | 31616 | 0.20 | 0.21 | 0.22 | 0.26 | -0.01 | -0.02 | -0.06 |
|  | PHE:CG | 4514 | 0.33 | 0.36 | 0.35 | 0.39 | -0.03 | -0.02 | -0.06 |
|  | PHE:CZ | 15421 | 0.21 | 0.22 | 0.22 | 0.27 | -0.01 | -0.01 | -0.06 |
|  | TRP:CD | 4845 | 0.25 | 0.27 | 0.27 | 0.31 | -0.01 | -0.02 | -0.05 |
|  | TRP:CE | 6967 | 0.22 | 0.25 | 0.26 | 0.29 | -0.03 | -0.04 | -0.07 |
|  | TRP:CG | 1247 | 0.38 | 0.42 | 0.42 | 0.42 | -0.04 | -0.03 | -0.03 |
|  | TRP:CH | 4579 | 0.20 | 0.21 | 0.22 | 0.30 | -0.01 | -0.02 | -0.10 |
|  | TRP:CZ | 8452 | 0.20 | 0.23 | 0.23 | 0.31 | -0.03 | -0.03 | -0.10 |
|  | TYR:CD | 18991 | 0.27 | 0.29 | 0.29 | 0.31 | -0.02 | -0.02 | -0.04 |
|  | TYR:CE | 18316 | 0.21 | 0.23 | 0.23 | 0.28 | -0.02 | -0.02 | -0.07 |
|  | TYR:CG | 3550 | 0.39 | 0.41 | 0.41 | 0.43 | -0.02 | -0.02 | -0.04 |
|  | TYR:CZ | 6243 | 0.28 | 0.32 | 0.32 | 0.36 | -0.04 | -0.04 | -0.08 |
| C•N(sp2) | ARG:NE | 6294 | 0.40 | 0.41 | 0.38 | 0.38 | -0.01 | 0.02 | 0.02 |
|  | ARG:NH | 16376 | 0.23 | 0.35 | 0.35 | 0.44 | -0.11 | -0.12 | -0.21 |
|  | ASN:ND | 15935 | 0.34 | 0.38 | 0.37 | 0.40 | -0.03 | -0.02 | -0.05 |
|  | GLN:NE | 12472 | 0.32 | 0.38 | 0.36 | 0.40 | -0.06 | -0.04 | -0.07 |
|  | HIS:ND | 6443 | 0.32 | 0.34 | 0.34 | 0.37 | -0.02 | -0.02 | -0.05 |
|  | HIS:NE | 8412 | 0.34 | 0.36 | 0.35 | 0.38 | -0.03 | -0.01 | -0.04 |
|  | TRP:NE | 10263 | 0.36 | 0.36 | 0.36 | 0.40 | 0.00 | 0.00 | -0.04 |
| C•O(sp2) | ASN:OD | 17152 | 0.40 | 0.43 | 0.43 | 0.45 | -0.03 | -0.03 | -0.05 |
|  | ASP:OD | 25044 | 0.39 | 0.44 | 0.44 | 0.46 | -0.05 | -0.05 | -0.07 |
|  | GLN:OE | 13017 | 0.35 | 0.39 | 0.38 | 0.40 | -0.04 | -0.03 | -0.05 |
|  | GLU:OE | 20293 | 0.34 | 0.40 | 0.39 | 0.41 | -0.06 | -0.05 | -0.06 |
| C•O(sp3) | SER:OG | 27276 | 0.42 | 0.42 | 0.41 | 0.43 | -0.01 | 0.00 | -0.01 |
|  | THR:OG | 24623 | 0.41 | 0.42 | 0.41 | 0.41 | -0.01 | 0.00 | 0.00 |
|  | TYR:OH | 24471 | 0.38 | 0.40 | 0.40 | 0.44 | -0.03 | -0.02 | -0.06 |

### Supplementary Table 7. The most favored hydrogen bond angles (donor–H–acceptor) and the median absolute deviation (MAD) of the hydrogen bond distributions from *Top*2018 PDB structures and from AlphaFold2, AlphaFold3 and ESMFold predictions.

| **H–bond type** | | | ***n*** | **Most favored angle**  **(peak) (º)** | | | | **Δpeak (º)** | | | **Median abs. deviation**  **(MAD) (º)** | | | | **ΔMAD (º)** | | |
| --- | --- | --- | --- | --- | --- | --- | --- | --- | --- | --- | --- | --- | --- | --- | --- | --- | --- |
| *type* | *acceptor* | *donor* |  | *PDB* | *AF2* | *AF3* | *ESM* | *PDB–AF2* | *PDB–AF3* | *PDB–ESM* | *PDB* | *AF2* | *AF3* | *ESM* | *PDB–AF2* | *PDB–AF3* | *PDB–ESM* |
| charge_charge | ASP-OD | ARG-NE | 579 | 161 | 161 | 156 | 150 | 1 | 5 | 5 | 8 | 10 | 11 | 10 | -1 | -3 | -2 |
| charge_charge | GLU-OE | ARG-NE | 564 | 164 | 165 | 165 | 162 | 0 | -1 | -1 | 8 | 10 | 11 | 10 | -2 | -3 | -1 |
| charge_dipole | ASN-OD1 | ARG-NH | 669 | 158 | 156 | 141 | 139 | 2 | 17 | 17 | 9 | 10 | 10 | 10 | -1 | -1 | -1 |
| charge_charge | ASP-OD | ARG-NH | 2153 | 158 | 155 | 137 | 138 | 3 | 21 | 21 | 8 | 10 | 10 | 10 | -3 | -3 | -2 |
| charge_dipole | GLN-OE1 | ARG-NH | 575 | 159 | 156 | 157 | 146 | 3 | 2 | 2 | 7 | 10 | 10 | 10 | -3 | -2 | -3 |
| charge_charge | GLU-OE | ARG-NH | 2004 | 156 | 148 | 156 | 143 | 8 | 0 | 0 | 7 | 10 | 10 | 10 | -2 | -3 | -2 |
| charge_dipole | SER-OG | ARG-NH | 627 | 157 | 148 | 140 | 137 | 9 | 17 | 17 | 9 | 10 | 10 | 10 | -1 | -1 | -1 |
| charge_dipole | THR-OG1 | ARG-NH | 512 | 154 | 149 | 138 | 142 | 5 | 16 | 16 | 9 | 9 | 9 | 8 | -1 | 0 | 0 |
| charge_dipole | TYR-OH | ARG-NH | 532 | 155 | 152 | 141 | 138 | 3 | 14 | 14 | 8 | 11 | 9 | 9 | -2 | -1 | -1 |
| dipole_dipole | ASN-OD1 | ASN-ND2 | 1011 | 164 | 160 | 156 | 150 | 4 | 9 | 9 | 8 | 9 | 9 | 10 | -1 | -2 | -2 |
| dipole_charge | ASP-OD | ASN-ND2 | 1738 | 165 | 162 | 162 | 150 | 3 | 4 | 4 | 7 | 9 | 10 | 10 | -2 | -2 | -2 |
| dipole_dipole | GLN-OE1 | ASN-ND2 | 531 | 167 | 159 | 152 | 153 | 8 | 16 | 16 | 8 | 9 | 10 | 10 | -1 | -2 | -2 |
| dipole_charge | GLU-OE | ASN-ND2 | 1257 | 169 | 167 | 163 | 158 | 2 | 6 | 6 | 8 | 10 | 10 | 10 | -2 | -3 | -2 |
| dipole_dipole | SER-OG | ASN-ND2 | 765 | 161 | 164 | 157 | 156 | -2 | 5 | 5 | 8 | 10 | 10 | 10 | -2 | -2 | -1 |
| dipole_dipole | THR-OG1 | ASN-ND2 | 749 | 168 | 164 | 158 | 147 | 5 | 11 | 11 | 8 | 9 | 10 | 10 | -1 | -1 | -2 |
| dipole_dipole | TYR-OH | ASN-ND2 | 530 | 163 | 149 | 151 | 146 | 14 | 12 | 12 | 9 | 10 | 9 | 10 | -1 | 0 | -1 |
| dipole_dipole | ASN-OD1 | GLN-NE2 | 718 | 162 | 160 | 140 | 146 | 2 | 22 | 22 | 9 | 8 | 11 | 10 | 1 | -2 | -1 |
| dipole_charge | ASP-OD | GLN-NE2 | 1462 | 160 | 159 | 151 | 148 | 1 | 9 | 9 | 7 | 8 | 8 | 7 | -1 | -1 | 0 |
| dipole_charge | GLU-OE | GLN-NE2 | 1072 | 168 | 166 | 166 | 164 | 2 | 2 | 2 | 7 | 9 | 9 | 10 | -2 | -2 | -3 |
| dipole_dipole | SER-OG | GLN-NE2 | 557 | 161 | 158 | 145 | 142 | 3 | 16 | 16 | 9 | 10 | 10 | 9 | -1 | -1 | -1 |
| dipole_dipole | THR-OG1 | GLN-NE2 | 578 | 166 | 164 | 157 | 155 | 2 | 9 | 9 | 8 | 9 | 10 | 10 | -1 | -2 | -2 |

### Supplementary Table 8. The most favored hydrogen bond angles (donor–H–acceptor angle) and the median absolute deviation (MAD) of the hydrogen bond distributions from *Top*2018 PDB structures and from AlphaFold2, AlphaFold3 and ESMFold predictions.

| **H–bond type** | | | ***n*** | **Most favored angle**  **(peak) (Å)** | | | | **Δpeak (Ǎ)** | | | **Median abs. deviation**  **(MAD) (Å)** | | | | **ΔMAD (Å)** | | |
| --- | --- | --- | --- | --- | --- | --- | --- | --- | --- | --- | --- | --- | --- | --- | --- | --- | --- |
| *type* | *acceptor* | *donor* |  | *PDB* | *AF2* | *AF3* | *ESM* | *PDB–AF2* | *PDB–AF3* | *PDB–ESM* | *PDB* | *AF2* | *AF3* | *ESM* | *PDB–AF2* | *PDB–AF3* | *PDB–ESM* |
| charge_charge | ASP-OD | ARG-NE | 579 | 2.84 | 2.79 | 2.87 | 3.29 | 0.05 | -0.03 | -0.45 | 0.08 | 0.11 | 0.25 | 0.24 | -0.03 | -0.17 | -0.16 |
| charge_charge | GLU-OE | ARG-NE | 564 | 2.84 | 2.81 | 2.94 | 3.08 | 0.02 | -0.11 | -0.24 | 0.09 | 0.13 | 0.24 | 0.25 | -0.04 | -0.15 | -0.17 |
| charge_dipole | ASN-OD1 | ARG-NH | 669 | 2.87 | 2.79 | 2.75 | 3.09 | 0.08 | 0.12 | -0.22 | 0.08 | 0.11 | 0.24 | 0.29 | -0.04 | -0.16 | -0.22 |
| charge_charge | ASP-OD | ARG-NH | 2153 | 2.87 | 2.73 | 2.69 | 3.31 | 0.14 | 0.18 | -0.44 | 0.08 | 0.09 | 0.22 | 0.36 | 0.00 | -0.14 | -0.28 |
| charge_dipole | GLN-OE1 | ARG-NH | 575 | 2.91 | 2.80 | 2.93 | 3.21 | 0.11 | -0.02 | -0.30 | 0.08 | 0.12 | 0.26 | 0.34 | -0.03 | -0.18 | -0.25 |
| charge_charge | GLU-OE | ARG-NH | 2004 | 2.89 | 2.73 | 2.73 | 3.21 | 0.15 | 0.15 | -0.33 | 0.09 | 0.09 | 0.24 | 0.32 | 0.00 | -0.15 | -0.23 |
| charge_dipole | SER-OG | ARG-NH | 627 | 2.92 | 2.88 | 3.13 | 3.23 | 0.04 | -0.21 | -0.32 | 0.09 | 0.13 | 0.21 | 0.27 | -0.04 | -0.12 | -0.18 |
| charge_dipole | THR-OG1 | ARG-NH | 512 | 2.92 | 2.90 | 2.97 | 3.33 | 0.02 | -0.05 | -0.41 | 0.09 | 0.12 | 0.19 | 0.28 | -0.03 | -0.10 | -0.19 |
| charge_dipole | TYR-OH | ARG-NH | 532 | 2.95 | 2.92 | 3.33 | 3.28 | 0.03 | -0.38 | -0.33 | 0.09 | 0.15 | 0.22 | 0.27 | -0.07 | -0.13 | -0.19 |
| dipole_dipole | ASN-OD1 | ASN-ND2 | 1011 | 2.91 | 2.86 | 2.99 | 2.98 | 0.05 | -0.08 | -0.07 | 0.08 | 0.09 | 0.15 | 0.21 | -0.01 | -0.08 | -0.13 |
| dipole_charge | ASP-OD | ASN-ND2 | 1738 | 2.91 | 2.79 | 2.79 | 3.11 | 0.12 | 0.12 | -0.20 | 0.09 | 0.07 | 0.17 | 0.24 | 0.01 | -0.08 | -0.15 |
| dipole_dipole | GLN-OE1 | ASN-ND2 | 531 | 2.92 | 2.85 | 3.15 | 3.17 | 0.08 | -0.23 | -0.25 | 0.08 | 0.09 | 0.18 | 0.29 | -0.01 | -0.10 | -0.21 |
| dipole_charge | GLU-OE | ASN-ND2 | 1257 | 2.92 | 2.79 | 2.89 | 3.21 | 0.12 | 0.03 | -0.29 | 0.09 | 0.09 | 0.20 | 0.26 | 0.00 | -0.11 | -0.17 |
| dipole_dipole | SER-OG | ASN-ND2 | 765 | 2.95 | 2.92 | 3.04 | 3.02 | 0.03 | -0.09 | -0.07 | 0.09 | 0.10 | 0.16 | 0.19 | -0.01 | -0.07 | -0.09 |
| dipole_dipole | THR-OG1 | ASN-ND2 | 749 | 2.96 | 2.91 | 2.97 | 3.01 | 0.05 | -0.01 | -0.05 | 0.09 | 0.09 | 0.14 | 0.21 | 0.00 | -0.05 | -0.12 |
| dipole_dipole | TYR-OH | ASN-ND2 | 530 | 2.97 | 2.93 | 2.97 | 3.36 | 0.04 | 0.00 | -0.39 | 0.10 | 0.14 | 0.19 | 0.25 | -0.04 | -0.08 | -0.15 |
| dipole_dipole | ASN-OD1 | GLN-NE2 | 718 | 2.92 | 2.85 | 2.95 | 3.09 | 0.07 | -0.03 | -0.16 | 0.08 | 0.09 | 0.17 | 0.18 | -0.01 | -0.09 | -0.09 |
| dipole_charge | ASP-OD | GLN-NE2 | 1462 | 2.92 | 2.79 | 2.82 | 2.91 | 0.13 | 0.10 | 0.01 | 0.09 | 0.07 | 0.14 | 0.18 | 0.02 | -0.05 | -0.09 |
| dipole_charge | GLU-OE | GLN-NE2 | 1072 | 2.90 | 2.79 | 2.90 | 3.19 | 0.11 | 0.00 | -0.29 | 0.08 | 0.07 | 0.18 | 0.24 | 0.01 | -0.10 | -0.16 |
| dipole_dipole | SER-OG | GLN-NE2 | 557 | 2.96 | 2.90 | 3.12 | 3.29 | 0.05 | -0.16 | -0.33 | 0.09 | 0.13 | 0.17 | 0.22 | -0.04 | -0.08 | -0.13 |
| dipole_dipole | THR-OG1 | GLN-NE2 | 578 | 2.97 | 2.94 | 3.00 | 3.32 | 0.03 | -0.04 | -0.35 | 0.10 | 0.13 | 0.18 | 0.22 | -0.03 | -0.08 | -0.12 |

### Supplementary Table 9. Secondary structure assignments from *Top*2018 PDB structures and from AlphaFold2 and AlphaFold3 predictions. Percent (%) correct is defined as the number of residues whose AlphaFold‐assigned secondary structure matches the corresponding PDB assignment, divided by the total number of PDB residues.

| **2º structure type** | **Count (PDB)** | **Count (AlphaFold2)** | **% correct**  **(AlphaFold2)** | **Count**  **(AlphaFold3)** | **% correct**  **(AlphaFold3)** | **Count**  **(ESMFold)** | **% correct**  **(ESMFold)** |
| --- | --- | --- | --- | --- | --- | --- | --- |
| α-helix | 242198 | 244400 | 96.6 | 244759 | 96.5 | 247522 | 96.1 |
| β-sheet | 174650 | 174803 | 96.0 | 175368 | 96.0 | 175002 | 94.6 |
| Other 2º structures | 191071 | 191536 | 92.1 | 190910 | 91.4 | 189579 | 83.6 |
| Coil | 145746 | 142926 | 90.3 | 142628 | 89.6 | 141562 | 86.8 |

### Supplementary Table 10. Multi-temperature X-ray diffraction datasets used to test AlphaFold3’ ability to generate ensembles.

| **PDB ID** | **Protein** | **Temp (K)** | **Resolution** | **Reference** |
| --- | --- | --- | --- | --- |
| 7LN7 | Proteinase K | 277 | 1.02 | ^115^ |
| 8SOG | Proteinase K | 313 | 1.13 | ^55^ |
| 8SQV | Proteinase K | 333 | 1.22 | ^55^ |
| 7LFG | Thaumatin | 277 | 1.22 | ^115^ |
| 5KW3 | Thaumatin | 278 | 1.55 | ^59^ |
| 7LLP | Lysozyme | 277 | 1.13 | ^115^ |
| 5KXO | Lysozyme | 278 | 1.2 | ^59^ |
| 6UCW | KSI | 250 | 1.25 | ^57^ |
| 6U1Z | KSI | 280 | 1.5 | ^57^ |
| 7MHG | Mpro | 240 | 1.53 | ^116^ |
| 7MHH | Mpro | 277 | 2.19 | ^116^ |
| 7MHI | Mpro | 298 | 1.88 | ^116^ |
| 7MHK | Mpro | 310 | 1.96 | ^116^ |
| 6B8T | PTP1B | 240 | 1.85 | ^56^ |
| 6B8X | PTP1B | 278 | 1.74 | ^56^ |
| 5TOF | ubiquitin variant u7ub25 | 298 | 1.12 | ^58^ |
| 5TOG | ubiquitin variant u7ub25.2540 | 298 | 1.08 | ^58^ |
| 5KUZ | Cyclophilin A | 278 | 1.7 | ^59^ |
| 4YUJ | Cyclophilin A | 240 | 1.42 | ^60^ |
| 4YUK | Cyclophilin A | 260 | 1.48 | ^60^ |
| 4YUL | Cyclophilin A | 280 | 1.42 | ^60^ |
| 4YUM | Cyclophilin A | 300 | 1.5 | ^60^ |
| 4YUN | Cyclophilin A | 310 | 1.58 | ^60^ |
| 4YUO | Cyclophilin A | 273 | 1.2 | ^60^ |
| 4YUP | Cyclophilin A | 273 | 1.75 | ^60^ |

71.	Database, A. P. S. AlphaFold Protein Structure Database. https://alphafold.ebi.ac.uk/faq.

72.	GitHub - google-deepmind/alphafold: Open source code for AlphaFold 2. *GitHub* https://github.com/google-deepmind/alphafold.

90.	Manshour, N., Ren, J. Z., Esmaili, F., Bergstrom, B. & Xu, D. Comprehensive Evaluation of AlphaFold-Multimer, AlphaFold3 and ColabFold, and Scoring Functions in Predicting Protein-Peptide Complex Structures. (2025).

113.	Evans, R., M., O. ’neill, A., P., N., A. & A., S. Protein complex prediction with AlphaFold-Multimer. *biorxiv* (2022).

114.	Mehdiabadi, M., Tosatto, S. C. E. & Piovesan, D. Modeling intrinsically disordered regions from AlphaFold2 to AlphaFold3. *Protein Sci* **35**, e70426 (2026).

115.	Yabukarski, F., Doukov, T., Mokhtari, D. A., Du, S. & Herschlag, D. Evaluating the impact of X-ray damage on conformational heterogeneity in room-temperature (277 K) and cryo-cooled protein crystals. *Acta Crystallogr D Struct Biol* **78**, 945–963 (2022).

116.	Ebrahim, A. *et al.* The temperature-dependent conformational ensemble of SARS-CoV-2 main protease (Mpro). *IUCrJ* **9**, 682–694 (2022).
